# Diazotrophic bacteria from maize exhibit multifaceted plant growth promotion traits in multiple hosts

**DOI:** 10.1101/2020.05.17.100859

**Authors:** Shawn M. Higdon, Tania Pozzo, Emily J. Tibbett, Colleen Chiu, Richard Jeannotte, Bart C. Weimer, Alan B. Bennett

**Affiliations:** Department of Plant Sciences, University of California, Davis, California 95616; Department of Population Health and Reproduction, University of California, Davis, California 95616

## Abstract

Sierra Mixe maize is a geographically remote landrace variety grown on nitrogen-deficient fields in Oaxaca, Mexico that meets its nutritional requirements without synthetic fertilizer by associating with free-living diazotrophs comprising the microbiota of its aerial root mucilage. We selected nearly 500 diazotrophic bacteria isolated from Sierra Mixe maize mucilage and sequenced their genomes. Comparative genomic analysis demonstrated that isolates represented diverse genera and possessed multiple marker genes for mechanisms of direct plant growth promotion (PGP). In addition to nitrogen fixation, we examined deamination of 1-amino-1-cyclopropanecarboxylic acid, biosynthesis of indole-3-acetic acid, and phosphate solubilization. Implementing *in vitro* colorimetric assays revealed each isolate’s potential to confer the alternative PGP activities that corroborated genotype and pathway content. We examined the ability of mucilage diazotrophs to confer PGP by direct inoculation of clonally propagated potato plants *in planta*, which led to the identification of bio-stimulant candidates that were tested for PGP by inoculating a conventional maize variety. The results indicate that, while many diazotrophic isolates from Sierra Mixe maize possessed genotypes and *in vitro* phenotypes for targeted PGP traits, a subset of these organisms promoted the growth of potato and conventional maize using multiple promotion mechanisms.

## Introduction

Plants are sessile organisms that interact with bacteria in their surrounding environment to alleviate abiotic and biotic stresses. Utilizing small molecules as chemical signals secreted in root exudates, plants may acquire an associated microbiome enriched for plant growth promoting bacteria (PGPB) that exhibit an array of plant growth promotion (PGP) mechanisms [1]. While some members of plant microbiomes facilitate PGP indirectly by inhibiting the growth of opportunistic microbes acting as parasites, others exhibit direct mechanisms of PGP by providing bioactive metabolites utilized by the host [2]. Well described traits for direct forms of PGP include biological nitrogen fixation (BNF), biosynthesis of auxins such as Indole-3-Acetic acid (IAA), promotion of underground root growth via utilization of 1-amino-1-cyclopropanecarboxylic acid (ACC) as a nitrogen source through the action of ACC deaminase, and the ability to liberate phosphate for plant uptake from biologically inert forms in the environment [3–5]. As the number of PGPB identified with the ability to confer these beneficial traits continues to rise through advances in next-generation sequencing, microbe-assisted crop production and development of crop bio-stimulants are emerging as feasible solutions to achieve global food security without marginalizing the environment [6–8].

The Sierra Mixe landrace of maize cultivated for generations on nutrient depleted fields without synthetic fertilizer was recently reported to derive 28-82% of its nitrogen from air and harbor PGPB capable of BNF (diazotrophs) within aerial root mucilage [9]. While Van Deynze et al. used low coverage shotgun metagenomic analysis of mucilage to uncover the presence of nitrogen fixation (*nif*) genes previously described as essential for diazotrophs (*nifHDKENB*) [10], whole genome sequence (WGS) analysis and BNF activity assay of mucilage bacterial isolates verified the presence of diazotrophic PGPB in the mucilage microbiome [11]. Taxonomic analysis of the mucilage bacterial isolates also revealed that isolated diazotrophs were predominantly classified to proteobacterial genera that comprise plant-associated microbiomes, exhibit endophytic lifestyles, and confer mechanisms of direct PGP independent of BNF [12]. Collectively, the evidence that Sierra Mixe maize thrives under cultivation in the absence of synthetic fertilizers and the characterization of confirmed diazotrophs classified to genera known to confer an array of PGP mechanisms suggests that the mucilage-derived bacterial isolates may be a useful tool to develop novel bio-stimulants for conventional maize varieties or other crops.

In addition to establishing that the mucilage bacterial isolates are diazotrophic, we hypothesized that these mucilage isolates from Sierra Mixe maize may exhibit PGP mechanisms beyond BNF. Secondarily, we hypothesized these capabilities are not limited to maize but rather the isolates are capable of conferring PGP to other crop systems. We scanned genomes from ∼500 diazotrophs for marker genes indicative of direct PGP traits that included IAA biosynthesis (*ipdC/ppdC*), utilization of ACC as a nitrogen source (*acdS*) and phosphate solubilization (*pqqBCDEF* and *pqq-DH*). Determination of direct PGP activities using *in vitro* colorimetric assays verified that some isolates possessed the genotype and phenotype for these PGP traits. *In planta* inoculation of clonally propagated potato (*Solanum tuberosum*) assessed the potential of PGP activity to non-cereal crops and identified candidate PGPB among the isolates. To evaluate candidate PGPB as bio-stimulants outside the laboratory, we inoculated nutrient stressed potato plants in the greenhouse with synthetic communities (SynComs) of mucilage isolates. This revealed a single mucilage isolate that also demonstrated PGP activity in conventional maize when inoculated under nitrogen stress. Our results demonstrate that diazotrophs isolated from Sierra Mixe maize possess the genomic determinants for multiple forms of direct PGP, the associated activities and, surprisingly, that many of the isolates have the ability to confer PGP in other crop systems.

## Materials and methods

### Genome analysis and PGP gene mining

Whole genome sequences for mucilage isolates previously reported were used for this analysis [11]. The ‘hmmscan’ feature of HMMer v3.1b was used to identify protein coding sequences identified in each isolate genome annotation generated with Prokka 1.12 [13, 14]. TIGRFAMs used with ‘hmmscan’ to examine genome presence of homologous sequences to marker genes for alternative PGP mechanisms included: TIGR03394, TIGR01274, TIGR02108, TIGR02111, TIGR03859, TIGR02109, TIGR02110 and TIGR03074 [15]. Dendrograms were made by computing genomic distance between MinHash sketches (k-mer length of 31) of WGS assemblies for each mucilage isolate generated with Sourmash v3.1.0 [16]. LCA classification of MinHash sketches was performed using the GTDB v89 [17] database available at: (https://osf.io/gs29b/). Code used to generate MinHash sketches of isolate WGS assemblies is provided within the Snakemake workflow hosted on Github: (https://github.com/shigdon/snakemake_mucilage-isolates). Analysis of homologous sequences from isolate genomes that matched targeted TIGRFAMs was conducted in R 3.5.1 using a combination of base and tidyverse 1.2.1 packages [18]. Visualization of the results as a circular dendrogram with heatmaps of gene counts was accomplished with a combination of ComplexHeatmap 1.20.0 and ggtree 1.14.6 R packages [19, 20]. Source code for bacterial genome mining analyses are available at: (https://github.com/shigdon/R-Mucilage-isolate-pgp-genes).

### BNF assay

Data from the previously described ^15^N incorporation assay of mucilage isolates to assess the nitrogen fixing capabilities of mucilage isolates was re-analyzed alongside data generated from alternative non-*nif* bioassays reported in this study [11]. Briefly, microbial isolates were independently cultured in an N-deficient liquid medium under both an atmosphere enriched for ^15^N_2_ gas and a control atmosphere of compressed air. Cell cultures were pelleted, and the supernatant was subjected to LC-TOF/MS analysis to identify and quantify the intensity of N-containing masses that were common to both the experimental and control atmospheric conditions. Peak intensities for common N-containing masses under both conditions were summed and used to generate ^15^N/^14^N ratios that served as metrics to score the relative N-fixation capability of isolates in the collection.

### ACC utilization assay

To screen isolates for the ability to utilize 1-amino-1cyclopropane carboxylic acid (ACC) as an independent source of nitrogen through ACC deaminase activity, a protocol in 96-well format was established by using previously described methods [21–23]. Each bacterial isolate was re-activated from cryogenically stored Microbank vials by using sterile inoculation loops to transfer a single cryobead to designated wells within a 2.2 mL deep block 96-well plate filled with 0.6 mL of liquid LB medium. Each isolate was cultured with shaking at 150 rpm under an incubation temperature of 28°C until mid-late log phase of growth was achieved. Bacterial cultures were pelleted by centrifugation at 2,178 × *g* for 20 minutes using the Sorvall legend RT benchtop centrifuge. The LB was removed and pellets were washed twice with 400 µL of 0.9% NaCl saline solution (w/v). To perform the assay, each well of a 96-deep well plate was filled with 200 µL of DF salts minimal medium (4g KH_2_PO_4_ · L^-1^, 6g Na_2_HPO_4_ · L^-1^, 0.2g MgSO_4_-7H_2_O · L^-1^, 2g glucose · L^-1^, 2 g gluconic acid · L^-1^, 2 g citric acid · L^-1^, 1 mg FeSO_4_-7H_2_O · L^-1^, 10 µg H_3_BO_3_ · L^-1^, 11.2 µg MnSO_4_-H_2_O · L^-1^, 125 µg ZnSO_4_-7H_2_O · L^-1^, 78.2 µg CuSO_4_-5H_2_O · L^-1^, 10 µg MoO_3_ · L^-1^) supplemented with 3 mM of fresh ACC (final concentration) and inoculated with a 10 µL volume of the washed cells (resuspended in 400 µL of 0.9% saline solution). Mock-inoculated wells (10 µL of sterile water) served as a negative control for microbial inoculation and each inoculation treatment was assayed with triplicate biological replication. Following inoculation of the assay medium, the initial OD_600_ values were recorded using the Synergy Mx microplate reader (Biotek) and the plates were incubated statically for 4 days at 28°C. Final OD_600_ values were recorded and the relative growth rate on ACC was calculated as the ratio of the day 4 OD_600_ / initial OD_600_.

### Auxin biosynthesis assay

Measuring the production of auxin by each pure culture was achieved by establishing a colorimetric screening assay adapted from previously described methods in 96-well plate format that utilize the Salkowski reagent originally described by Gordon and Weber [24–26]. Each bacterial isolate was revived from cryogenic storage, cultured, washed and resuspended in 400 µL of 0.9% saline solution (w/v) as described for the ACC utilization assay. To perform the colorimetric assay, each well of a 96-deep well plate was filled with 1 mL of liquid LB medium containing a final concentration of 500 µg/mL L-Tryptophan (LB+TRP medium). The LB+TRP medium was inoculated with 50 µL of the 400 µL washed bacterial cell suspension and the cultures were statically incubated for four days at 28°C. Afterwards, cells were pelleted by centrifugation at 3,020 × *g* for 25 minutes and 100 µL of supernatant from each culture was combined and gently mixed with 100 µL of the Salkowski reagent in a clear flat-bottom 96-well plate. The plates were incubated for 30 minutes in darkness prior to measuring absorbance at 535 nm using the Synergy Mx microplate reader, and each culture was assayed in triplicate alongside a mock-inoculated negative control (50 µL sterile water). Quantification of IAA biosynthesis was done by using a linear standard curve (R^2^ = 0.99) of IAA suspended in LB+TRP medium.

### Phosphate solubilization assay

Screening of each microbial isolate’s ability to liberate soluble phosphate from inorganic forms was carried out in 96-well plate format by establishing an assay adapted from previously described methods [27, 28]. Each microbial isolate was revived, cultured, washed and resuspended in 400 µL of 0.9% saline solution (w/v) as described for the aforementioned colorimetric assays. The wells of a 2.2 mL 96-deep well plate were filled with 1 mL of the NBRIP medium described by Hu et al. that was modified by replacing calcium phosphate tribasic (Ca_3_(PO_4_)_2_) with hydroxy apatite (Ca_5_HO_13_P_3_) [29, 30]. Each well was inoculated with either 50 µL of the 400 µL washed cell suspension or sterile saline solution (negative control), and the 96-deep well plates were incubated for 4 days at 28°C without agitation. Following incubation, deep-well plates were centrifuged at 3,020 × *g* for 20 minutes and the supernatant was diluted 100-fold by combining 2 µL of supernatant with 198 µL of sterile water in each well of a clear flat bottom 96-well microplate. Triplicate biological replicates of the diluted supernatant were each combined with 50 µL of fresh working reagent for the assay, which was prepared by combining stock solutions of 51 mM ammonium molybdate, 4.9 N sulfuric acid, 0.9 µM ascorbic acid and 1.2 mM antimony potassium tartrate at a ratio of 1.5:5:3:0.5, respectively. Each plate was then mixed well with a microplate shaker, allowed to equilibrate for 30 minutes, and subjected to absorbance measurement at a wavelength of 630 nm using a Synergy Mx microplate reader. The amount of phosphate liberated by each isolate was quantified by comparing the absorbance values to a linear standard curve generated by measuring the absorbance of KH_2_PO_4_ in diluted the modified NBRIP medium.

### *In vitro* PGP assay data analysis

Data collected from the ACC deaminase utilization, BNF, IAA biosynthesis, and Phosphate solubilization assays were analyzed together using an R environment built with R version 3.5.1. Box plots (S2 Fig) were generated by formatting assay data using base and tidyverse 1.2.1 R packages [18]. Circular heatmaps featuring dendrograms of mucilage isolates were created with R packages that include ComplexHeatmap 1.20.0 and ggtree 1.14.6 [19, 20]. Statistical analysis of linear models was carried out using base R stats packages and the Estimated Marginal Means (emmeans 1.4.1, aka Least-Squares Means) R package [31]. The R code used to analyze data and plot results for each isolate genome in R, along with associated meta and raw data files, is provided at: (https://github.com/shigdon/R-Mucilage-isolate-pgp-assay).

### Bacterial culture for *in planta* potato inoculation assay

Cell cultures were initiated by diluting 50 µL of bacterial glycerol stock solutions to a final volume of 1 mL using liquid LB media and were incubated at 28°C for three days. Bacterial cell suspensions were washed using a 0.9% saline solution (w/v) and a subsample of 10 µL was diluted with 200 µL of 0.9 % saline solution (w/v) using a 96-well clear bottom polypropylene microplate. Cell densities for each culture were measured using OD_600nm_ with a Biotek Synergy MX plate reader. Preparation of the final inoculum for each isolate applied to the base of potato plantlets was achieved by diluting 25 µL of each saline-washed bacterial cell suspension in 475 µL of the 0.9% saline solution (w/v).

### *In planta* potato plantlet inoculation assay to screen for PGP

To assess the growth of potato plantlets, potato nodal cutting (PNC) media was used following protocols described by Johnston-Monje [32]. The PNC medium was transferred to 22×11 round bottom glass tubes (25 mL) and placed on a slant until solidified with subsequent potato plantlet transplantation into each tube. Russet Burbank-Idaho potato plantlets (New Brunswick Plant Propagation Center, Canada) were used for the inoculation assay. Triplicate biological replication of potato plantlet inoculation with 100 µL of individual bacterial isolate cell suspensions were done by pipetting the inoculum to the surface of each PNC medium slant containing a transplanted potato plantlet. Mock-inoculation control solutions were done using 100 µL of the 0.9% saline solution (w/v). The vials were capped and wrapped with surgical tape to minimize contamination. Inoculated potato plantlets were incubated in a growth chamber as described by Johnston-Monje [32] with 60-70% relative humidity for three weeks. Initial weights of inoculation vials were recorded and after three weeks of incubation, wet and dry weights were recorded for shoot and root biomass.

### Bacterial culture for SynCom inoculation experiment with potato grown in the greenhouse

Mucilage isolates were reactivated in a laminar flow hood by streaking cell suspensions from Microbank cryogenic storage onto LB or TY agar petri dishes incubated statically at 28°C for 24 hours. Each isolate was sub-cultured by taking a single colony of cells from each plate and inoculating it into 300 mL of TY liquid medium followed by incubation at 28°C for 24 hours with shaking at 200 rpm. Once the OD_600_ value of each liquid culture reached 1.0, the total volume of each culture was divided in half and each half was subjected to centrifugation for 10 minutes at 4°C, with a *rcf* (× *g*) value of 16,000. After decanting TY medium, cell pellets of each microbial isolate were gently resuspended in either 150 mL of 0.5X Hoagland liquid medium, or 0.5X N-free Hoagland liquid medium. SynComs were generated by combining 34 mL of each isolate’s Hoagland cell suspension to create inoculation treatments consisting of 3 mono-cultured mucilage isolate cells pooled in equal volumes. Mono-isolate inoculations of RLS potato plants were done using equal volumes of SynCom inoculation volumes and the isolates were cultured using the same method previously described.

### Mucilage isolate SynCom inoculation experiment with potatoes grown in the greenhouse

Clonally propagated potato plantlets of the Russet Burbank and Red La Soda (New Brunswick Plant Propagation Centre, Canada). Sterilized 150×22 mm test tubes were filled with 15 mL of a perlite:vermiculite (50:50) mixture in a sterile environment. Each prepared test tube was filled with 10 mL of inoculant comprised either by SynCom cell suspensions (Hoagland, N+ or N-) or non-inoculated Hoagland (N+ or N-) liquid medium. Potato Plantlet cuttings were transferred into the test tubes using sterilized scissors and forceps by nestling the stem of the plantlets 2 mm into the growth medium (substrate + Hoagland cell suspension). Inoculated plants in test tubes were sealed with micropore tape and incubated for 10 days. Incubation conditions were at 28 °C with a 16:8 photoperiod and 50% Relative Humidity. Once plantlets had successfully rooted into the test tube inoculation medium, plants were transplanted into 5-liter pots in the greenhouse and a second 10 mL inoculation treatment of SynCom or control solution was applied to the stem base of each plant. Pots were filled with a customized medium consisting of Perlite, Calcined Clay and Vermiculate at a ratio of (1:1:1). Greenhouse plants were watered using fertilized irrigation consisting of modified Hoagland formulations in RO water spanning two treatments: a high nitrogen treatment with 150 ppm N and a low nitrogen treatment of 15 ppm N [33]. Each pot received 500 mL of fertilized water per day with an increase to 1 L beginning after 60 days in the greenhouse. The treatment groups (inoculation x N-fertilization) were arranged using a randomized complete block design (RCBD) to account for positional effects from environmental variation in the greenhouse. Design of the factorial experiment consisted of 112 plants comprising four complete block replications that each included the two levels of fertilizer treatment, two potato genotypes, and seven inoculation treatments. After 90 days, plants were destructively sampled to measure shoot, root and tuber weights and record tuber number.

### Deconstructed SynCom inoculation experiment of potatoes grown in the greenhouse

Red La Soda potato plants for the deconstructed SynCom experiment were acquired, inoculated, incubated and transplanted to the greenhouse using the same resources, procedures and environmental conditions described for the initial SynCom inoculation experiment. However, rather than pooling three independent cultures of different isolates to form SynComs, potato plants were inoculated with 10 mL cell suspensions of individual isolate cultures or mock-inoculated controls (sterile 0.5X N^+^ or N^-^ Hoagland medium). Isolate cells were cultured and resuspended in either replete 0.5X Hoagland medium or 0.5X N-free Hoagland medium using identical procedures as those described for the initial SynCom inoculation experiment. Inoculated plants were arranged using an RCBD that included 6 full replications of each factorial treatment group covering two levels of fertilization (150 ppm N or 15 ppm N), and four different inoculation treatments (either one of three mono-isolate inoculations or a mock-inoculation). Incubated plants were grown in the greenhouse for 120 days and subjected to destructive sampling to measure shoot, root and tuber weights and record tuber number.

### Maize seed sterilization and germination

Maize seeds of the domesticated hybrid variety B73 x Mo17 (Plant ID: Ames19097) (U.S. National Plant Germplasm System, Agricultural Research Service, U.S. Department of Agriculture). Seeds were surface sterilized by subjecting 10 seeds that were placed in 50 mL conical tubes (Falcon) along with 15 mL of a 6 % (v/v) bleach solution with 0.1 % Tween 20 to horizontal shaking at a 45° angle for 20 minutes under a controlled temperature of 28°C. Following the sterilization wash, five replications of seed rinsing was carried out by horizontal shaking the seeds with an equivalent volume of fresh Milli-Q water for each rinse. Maize seeds were germinated in the laboratory by sowing at a depth of three centimeters into 2×6 plastic tray inserts that were filled with a customized medium comprised of Perlite, Calcined Clay and Vermiculate at a ratio of (1:1:1). Trays of sown seeds were placed in a controlled environment growth chamber that was held at 28°C with a relative humidity setting of 50% and a photoperiod of 16:8 (light:dark).

### Inoculation experiments with conventional maize in the greenhouse

The mucilage microbial isolates used for the B73 x Mo17 maize inoculation experiment were re-activated from cryo-storage by streaking onto LB-agar medium plates that were statically incubated at 28°C for 24 hours. Single colonies were sub-cultured in baffled Erlenmeyer flasks containing 200 mL of TY medium shaking (150 rpm) at 28°C. Once the cultures reached an OD_600_ value of 1.0 (∼ 1×10^9^ cells per 1 mL of liquid culture medium), the cells were pelleted by centrifugation for 30 minutes using the Sorvall Legend RT benchtop centrifuge under the conditions of 1,556 × *g* and 4°C. The TY medium was decanted, and the cells were resuspended using an equivalent volume of 0.5X N-free Hoagland medium. The N-free Hoagland cell suspensions were then applied to germinated maize seedlings that had developed their first true leaf using a soil-drench strategy in which 8 mL of the suspension was directly applied to the soil medium using a pipette. Inoculated seedlings were kept in the laboratory growth chamber for 2 days prior to greenhouse transplantation. Seedlings were transplanted into 7 gallon pots containing a customized soil mixture of Peat moss, Sand and Porous Calcined Clay (Greens Grade) and at a ratio of 3:2:1. Following transplantation, 8 mL of microbial cell culture suspended in N-free Hoagland liquid medium was applied to the base of each plant stalk. Plants were fed using RO water injected with modified Hoagland fertilizer solutions to deliver two nitrogen treatments; one high nitrogen treatment (150 ppm N) and one low nitrogen treatment (15 ppm N) [33]. Plants of each treatment group (inoculation x N-fertilization) were arranged using an RCBD to account for positional effects in the greenhouse.

### Statistical analysis and visualization of inoculation experiment results

Plant response variables were analyzed using R version 3.5.1. Statistical analysis of plant biomass accumulation measurements (i.e. gravimetric weights of shoots, underground roots, and tubers) and recorded tuber number were fit to linear models and subjected to student t-tests using the Estimated Marginal Means 1.4.1 (aka Least-Squares Means) R package [31]. Mean estimates for all inoculation groups were subjected to pairwise t-tests with *p*-value correction for multiple testing implemented with Dunnett’s test (comparison of ≥19 microbial inoculation treatments to a single mock-inoculated control) and/or the Tukey HSD method (all pairwise comparisons of 6 SynComs and a mock-inoculated control group, and all pairwise comparisons of 3 mono-isolate inoculations). Observed increases in response variables with microbial inoculation treatments were considered to be statistically significant relative to non-inoculated controls at adjusted p-values ≤ 0.01. Plots of mean estimates with confidence intervals were generated using base R functions. Bar plots were generated using the R packages ggplot2 3.2.1 and cowplot 1.0.0 [34]. Standard error bars were computed with custom R functions where the source code and raw data used for the analysis of the *in planta* potato inoculation experiments is available at https://github.com/shigdon/R-Mucilage-isolate-potato-inplanta-assay. Source code and raw data used for analysis of the SynCom potato inoculation experiments is available at https://github.com/shigdon/R-Mucilage-isolate-potato-SynCom. Source code and raw data used for analysis of the SynCom F deconstruction experiment is available at https://github.com/shigdon/R-Mucilage-isolate-potato-SynComF. Source code and raw data used for analysis of the B73xMo17 maize inoculation experiment is available at https://github.com/shigdon/R-Mucilage-isolate-maize-inoculation.

## Results

### Diazotroph genomes possessed marker genes for additional PGP functions

Querying open reading frames (i.e. genes) of mucilage diazotroph genomes for marker genes associated with forms of direct PGP alternative to BNF revealed that members of the isolate collection possessed the genetic underpinnings to confer ACC deamination, biosynthesis of IAA, and phosphate solubilization (S1 Table). Comparison of these results using a heatmap of gene presence counts plotted alongside a clustered dendrogram based on WGS variation (i.e. genome distance) revealed clusters among the non-*nif* PGP gene profiles of each diazotroph genome (S1 Fig). Cross-referencing previously reported taxonomy and essential *nif* gene content with results from mining each diazotroph genome for non-*nif* PGP marker genes produced a summary of PGP genome content tallied by genus (Table 1) [11]. This demonstrated that diazotroph genomes that possessed metabolic genes for BNF also possessed genomic traits for other direct PGP products.

**Table 1.** Number of isolates with PGP marker genes summarized by genus classification. ^a^Genetic markers for Biological Nitrogen Fixation (BNF) phenotypically positive isolates comprising the model for diazotrophic bacteria proposed by Dos Santos et al. [10]. Values were generated with previously reported genome mining data [11]. *^b^*Genetic marker for 1-amino-1-cyclopropanecarboxylic acid deaminase (ACCD). *^c^*Genetic marker for biosynthesis of the plant auxin indole-3-acetic acid (IAA). *^d^*Genetic markers for the mechanism to solubilize phosphate (PO_4_) from inorganic forms mediated by the genetic operon of *pqq* genes for biosynthesis of pyrroloquinoline quinone (PQQ) and PQQ-associated dehydrogenase (*pqq*-*DH*).

| Genus | BNF<sup>a</sup> |  |  |  |  |  | ACCD<sup>b</sup> | IAA<sup>c</sup> | PO<sub>4</sub><sup>d</sup> |  |  |  |  |  |
| --- | --- | --- | --- | --- | --- | --- | --- | --- | --- | --- | --- | --- | --- | --- |
|  | *nifH* | *nifD* | *nifK* | *nifE* | *nifN* | *nifB* | *acdS* | *ipdC* | *pqqB* | *pqqC* | *pqqD* | *pqqE* | *pqqF* | *pqq-DH* |
| *Acidovorax* | 2 | 0 | 0 | 0 | 0 | 0 | 2 | 2 | 2 | 2 | 2 | 2 | 0 | 2 |
| *Acinetobacter* | 10 | 0 | 0 | 0 | 0 | 0 | 2 | 11 | 10 | 10 | 10 | 10 | 0 | 11 |
| *Agrobacterium* | 0 | 0 | 0 | 0 | 0 | 0 | 0 | 11 | 0 | 0 | 0 | 0 | 0 | 11 |
| *Atlantibacter* | 0 | 0 | 0 | 0 | 0 | 0 | 3 | 3 | 0 | 0 | 0 | 0 | 0 | 3 |
| *Bacillus* | 1 | 0 | 0 | 0 | 0 | 0 | 0 | 1 | 0 | 0 | 0 | 0 | 0 | 0 |
| *Citrobacter* | 0 | 0 | 0 | 0 | 0 | 0 | 10 | 10 | 0 | 0 | 0 | 0 | 0 | 3 |
| *Curtobacterium* | 1 | 0 | 0 | 0 | 0 | 0 | 0 | 3 | 0 | 0 | 0 | 0 | 0 | 0 |
| *Enterobacter* | 18 | 18 | 18 | 18 | 18 | 18 | 71 | 71 | 0 | 0 | 0 | 3 | 0 | 71 |
| *Erwinia* | 0 | 0 | 0 | 0 | 0 | 0 | 4 | 4 | 4 | 4 | 4 | 4 | 4 | 4 |
| *Escherichia* | 0 | 0 | 0 | 0 | 0 | 0 | 1 | 1 | 0 | 0 | 0 | 0 | 0 | 1 |
| *Hafnia* | 0 | 0 | 0 | 0 | 0 | 0 | 0 | 3 | 0 | 0 | 0 | 0 | 0 | 0 |
| *Herbaspirillum* | 1 | 0 | 0 | 0 | 0 | 0 | 1 | 1 | 0 | 0 | 0 | 0 | 0 | 0 |
| *Klebsiella* | 26 | 26 | 26 | 26 | 26 | 26 | 26 | 26 | 0 | 0 | 0 | 0 | 0 | 26 |
| *Kosakonia* | 9 | 9 | 9 | 9 | 9 | 9 | 9 | 9 | 0 | 0 | 0 | 0 | 0 | 9 |
| *Lactococcus* | 0 | 0 | 0 | 0 | 0 | 0 | 0 | 23 | 0 | 0 | 0 | 17 | 0 | 0 |
| *Leifsonia* | 1 | 0 | 0 | 0 | 0 | 0 | 0 | 1 | 0 | 0 | 0 | 0 | 0 | 0 |
| *Lelliottia* | 0 | 0 | 0 | 0 | 0 | 0 | 23 | 23 | 0 | 0 | 0 | 0 | 0 | 23 |
| *Metakosakonia* | 25 | 25 | 25 | 25 | 25 | 25 | 26 | 26 | 0 | 0 | 0 | 1 | 0 | 26 |
| *Microbacterium* | 0 | 0 | 0 | 0 | 0 | 0 | 2 | 2 | 0 | 0 | 0 | 0 | 0 | 0 |
| *Micrococcus* | 2 | 0 | 0 | 0 | 0 | 0 | 0 | 2 | 0 | 0 | 0 | 0 | 0 | 0 |
| *Morganella* | 0 | 0 | 0 | 0 | 0 | 0 | 1 | 1 | 0 | 0 | 0 | 0 | 0 | 0 |
| *Pantoea* | 0 | 0 | 0 | 0 | 0 | 0 | 14 | 14 | 14 | 14 | 14 | 14 | 14 | 14 |
| *Pseudomonas* | 32 | 2 | 2 | 2 | 2 | 2 | 28 | 33 | 33 | 33 | 33 | 33 | 30 | 33 |
| *Rahnella* | 24 | 24 | 24 | 24 | 24 | 24 | 72 | 72 | 66 | 66 | 66 | 66 | 11 | 72 |
| *Raoultella* | 70 | 70 | 70 | 70 | 70 | 70 | 70 | 70 | 2 | 1 | 1 | 0 | 0 | 69 |
| *Rhodococcus* | 0 | 0 | 0 | 0 | 0 | 0 | 1 | 1 | 0 | 0 | 0 | 1 | 0 | 0 |
| *Serratia* | 0 | 0 | 0 | 0 | 0 | 0 | 17 | 17 | 0 | 0 | 0 | 1 | 0 | 17 |
| *Staphylococcus* | 0 | 0 | 0 | 0 | 0 | 0 | 0 | 1 | 0 | 0 | 0 | 0 | 0 | 0 |
| *Stenotrophomonas* | 15 | 0 | 0 | 0 | 0 | 0 | 0 | 15 | 0 | 0 | 0 | 1 | 0 | 14 |
| unassigned | 22 | 20 | 20 | 20 | 20 | 19 | 31 | 35 | 2 | 2 | 3 | 3 | 1 | 32 |
| **Total** | **259** | **194** | **194** | **194** | **194** | **193** | **414** | **492** | **133** | **132** | **133** | **156** | **60** | **441** |

While none of the diazotrophs contained homologous sequences to all of the genes for PGP markers queried, comparative analysis of PGP gene profiles of the diazotroph genomes revealed that isolates classified to proteobacterial genera were predominant among those with gene matches for both essential *nif* genes and other PGP traits. Among the 492 diazotroph genomes queried, 52% exhibited presence of the *nifH* gene (Table 1). Of these 259 genomes, 193 diazotrophs classified to *Enterobacter, Klebsiella, Kosakonia, Metakosakonia, Pseudomonas, Rahnella* and *Raoultella* had homologous sequences for the complete Dos Santos model of essential *nif* genes (Table 1, S1 Table). Interestingly, all diazotrophic genomes in this subset, referred to as the Dos Santos Positive (DSP) group [11], contained homologous coding sequences for the *acdS* and *ipdC/ppdC* genes. Although DSP isolate genomes had homologous sequences to the *acdS* and *ipdC/ppdC* genes, matches for these gene models were identified in the population of diazotrophic genomes surveyed at frequencies of 84% and 100%, respectively. In the case of PQQ genes, roughly 28% of all isolates examined contained homologous sequences to *pqqBCDE*, 12% had coding sequences homologous to *pqqF* and 90% had matches to *pqq-DH*. With the exception of *pqq-DH*, DSP isolates possessed sequences matching models for at least one of the *pqqBCDEF* genes but at a much lower frequency of 16.5%. Taken together, these observations warranted phenotypic assessment of each mucilage diazotroph for the targeted PGP traits.

### *In vitro* assays confirmed diazotrophic isolate potentials for known PGP traits

#### Mucilage isolates formed three groups of diazotrophs capable of BNF

Prior studies established that all of these mucilage isolates are diazotrophs using experimentally determined ^15^N/^14^N ratios (BNF ratios). Combining these phenotypic results with *nif* gene analysis revealed three distinct genotypic groups (NIF groups) based on possession of homologous sequences to essential *nif* genes (*nifHDKENB*) in the Dos Santos model [10, 11]. These NIF groups consisted of DSP isolates with genome sequences matching all six essential *nif* genes, the group consisting of genomes with incomplete versions of the Dos Santos model (Semi Dos Santos – SDS), and another that completely lacked matches to all essential *nif* genes (Dos Santos negative – DSN). This observation led to the hypothesis that diazotrophy is not likely to be the only growth stimulating capability of these organisms that may have multiple genomic underpinnings. Comparison of BNF ratio values with additional growth performance bioassays for alternative modes of direct PGP confirmed multi-trait phenotypes for many mucilage diazotrophs in the context of the three NIF groups (Fig 1, S2 Table). Plotting the distribution of BNF ratios by isolates categorized by NIF group membership revealed interquartile ranges with low variation in the dataset (S2 Fig A). Applying stringent criteria for diazotrophic isolates BNF phenotypes to be considered as outliers, defined as BNF ratios falling in the upper quartile beyond three times the interquartile range (IQR), identified 13 DSN isolates, 4 DSP isolates and 4 SDS isolates that displayed high BNF efficiencies.

**Fig 1.**
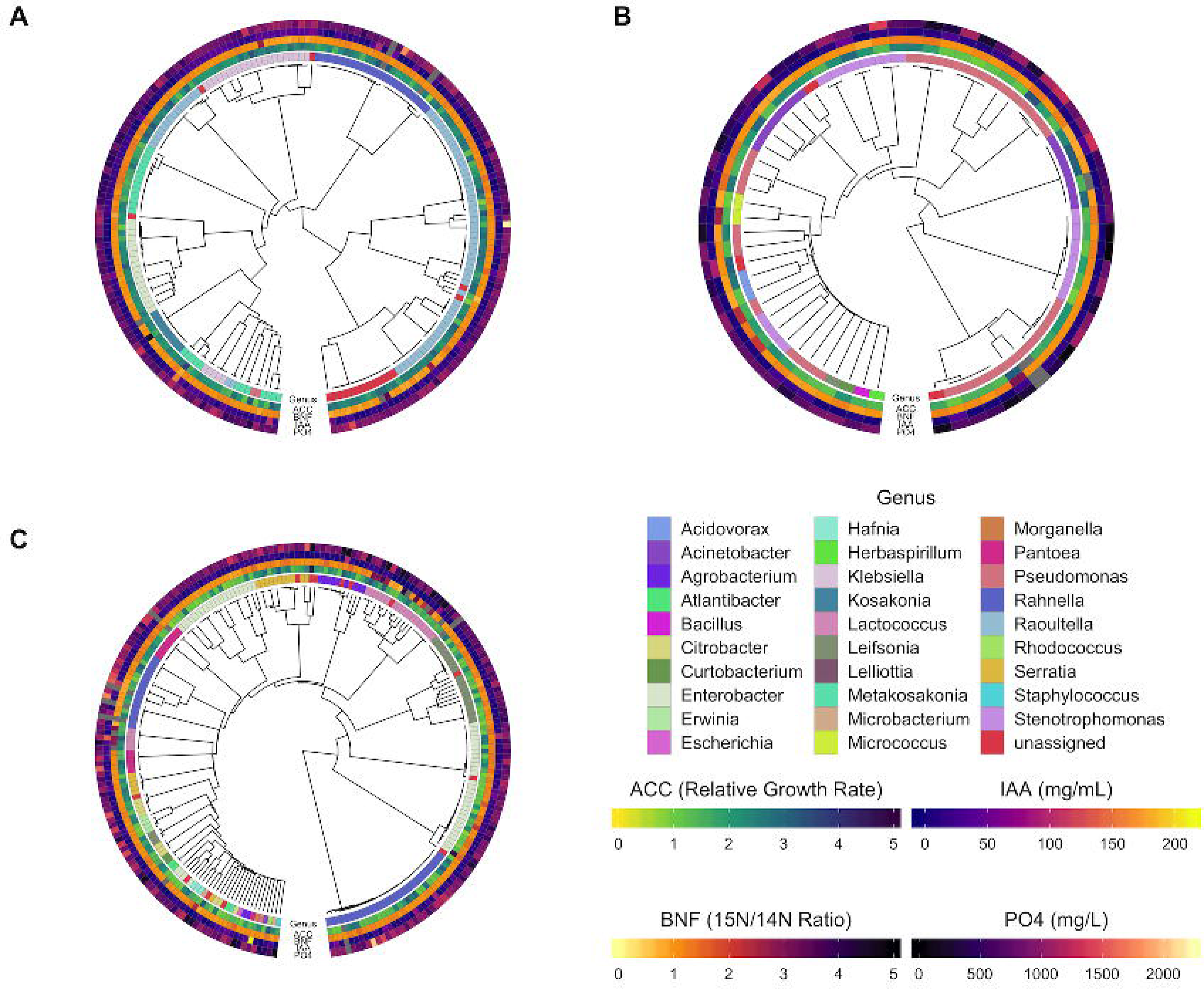
Dendrograms of mucilage diazotroph genomes with PGP assay phenotypes. Phenotypes for the PGP traits were presented in the context of the three NIF groups using heatmaps plotted alongside dendrograms of whole genome distances computed with Sourmash [16]. A) Diazotrophs of the Dos Santos Positive (DSP) group; B) Diazotrophs of the Semi-Dos Santos (SDS) group; C) Diazotrophs of the Dos Santos Negative (DSN) group. Previously reported BNF ratio values [11] were presented alongside results from *in vitro* biochemical assays for alternative forms of direct PGP. Phenotypic assays of all diazotrophs included assessment of biological nitrogen fixation (BNF), utilization of 1-amino-1-cyclopropane carboxylic acid (ACC) as a nitrogen source, biosynthesis of Indole-3-Acetic Acid (IAA) and liberation of phosphate (PO_4_). For the BNF assay, the ^15^N/^14^N ratio (BNF ratio) reflected the BNF capability of each isolate, as determined by summing peak intensities from N-containing biomarkers (see Methods) commonly identified by LC-MS of supernatant after cell culture on N-deficient medium in a ^15^N enriched atmosphere and air (control). ACC utilization reflected the relative growth rate of each diazotrophic isolate following culture on N-deficient medium with ACC provided as the individual N-source. IAA phenotypes represented the concentration of IAA detected by colorimetric assay in mg/mL following incubation with L-tryptophan. Phosphate solubilization phenotypes demonstrated the phosphate concentration in mg/L detected by colorimetric assay of supernatant after culturing each diazotroph in phosphate-deficient medium supplemented with hydroxyapatite.

Further analysis of the BNF ratios with a linear model of the data (BNF ratio ∼ 1 + NIF group) tested the hypothesis that differences in *nif* gene content (i.e. NIF group assignment) contributed to the observed variance in BNF efficiency. Analysis of variance (ANOVA) using the linear model revealed that NIF group assignment significantly (*p* = 0.027) contributed to variance in the BNF ratios observed among all diazotrophic isolates assayed. Subsequent pairwise comparisons of mean BNF ratios estimated for each NIF group revealed a significant positive difference of 0.11 in average BNF ratio values for DSN isolates over DSP isolates (*p* = 0.02, S3 Fig). Although the effect size of the difference between mean BNF ratio value between the DSN and DSP NIF groups was relatively small, this result confirmed that isolates without genomic possession of *nifHDKENB* displayed the highest observed BNF ratios on average relative to the SDS and DSP NIF group isolates that possessed some or all of the *nif* genes proposed to be essential for BNF activity [10], respectively. Taken together, these results confirmed the tested hypothesis as variation in *nifHDKENB* gene configurations significantly impacted the distribution and average of BNF ratios observed among the three NIF groups.

#### Mucilage isolates were successfully cultured with ACC as the source of nitrogen

Implementing a bioassay to evaluate the ability of each diazotrophic isolate to grow when cultured in a defined medium with ACC provided as the nitrogen source generated *in vitro* phenotypes that characterized their potential to confer ACC deaminase functionality. Results from this *in vitro* assay revealed that diazotrophic isolates across all three NIF groups displayed ACC utilization (Fig 1). Assessing isolate utilization of ACC based on NIF group membership showed differences in ACC utilization efficiency (S3 Table). Visualizing the distribution of data acquired from each isolate categorized by NIF group membership showed that the DSP group had a higher relative growth rate (RGR) overall when compared to the DSN and SDS groups (S2 Fig B). The observed RGR of each diazotroph comprising the DSP group spanned the inclusive range from 0.8 to 3.7 with a median RGR of 2.31 (S3 Table). SDS isolates exhibited RGR values of a similar range from 0.91 to 3.32 but had a lower median RGR at 1.63. Interestingly, the DSN group had a larger RGR range (0.76 to 4.74) but also had a lower median RGR of 1.5, which was the lowest of the three NIF groups.

To assess the significance of the observed differences in ACC utilization efficiency among the three isolate NIF groups, RGR values were fit to a linear model (RGR ∼ 1 + NIF group) to conduct ANOVA and test the hypothesis that NIF group membership impacted isolate utilization of ACC. Results from the ANOVA revealed that NIF group membership had a significant impact on isolate utilization of ACC (*p* = 4.96e^-16^). Additional pairwise comparisons of mean RGR differences for each NIF group showed that DSP isolates exhibited significantly higher (*p*-adj < 0.0001) amounts 0.52 and 0.44 over DSN and SDS isolates, respectively (S3 Fig). Collectively, these results demonstrated that DSP isolates had the highest growth rate among the three NIF groups when ACC was provided as the nitrogen source and confirmed the tested hypothesis.

#### Mucilage diazotrophs produced IAA from tryptophan

Examining mucilage diazotrophs for the ability to synthesize IAA *in vitro* demonstrated that diazotrophs from all three NIF groups produced this compound when provided with the metabolic precursor Trp (Fig 1). Assessing the ability of mucilage diazotrophs to produce IAA by NIF group revealed that isolates in the DSP group generated the highest median IAA concentration of 17.35 mg/mL, followed by median concentrations of 13.51 mg/mL and 8.4 mg/mL for the DSN and SDS groups, respectively (S3 Table). Observed median values followed trends similar to the calculated mean IAA concentrations among the isolate NIF groups. However, visualizing the distribution of the IAA bioassay data revealed 9 DSN isolates and 3 DSP isolates as high-producing outliers within their respective NIF groups using the same parameters for outlier designation as in the BNF assay data (S2 Fig C). These outliers contributed to the extended range of IAA concentrations observed for the DSN and DSP groups, which contained isolates with respective maximum values of 210.76 and 158.0 mg/mL IAA that were higher than the SDS group maximum of 92.2 (S3 Table). These results showed that while the largest NIF group of DSN isolates had the highest outlier number for IAA producing diazotrophs, the IAA concentrations generated by a substantial proportion of DSP isolates were sufficient in producing a higher average concentration for the group despite its smaller group size.

Linear modeling of the IAA biosynthesis assay ([IAA] ∼ 1 + NIF group) enabled testing the hypothesis that NIF group membership had a directional impact on the levels of IAA produced by the isolates. Conducting ANOVA on the dataset showed significant differences in isolate IAA production attributed by NIF group membership (*p* = 0.03). Additional pairwise comparison of differences between the NIF groups revealed that DSP isolates exhibited a significant increase of 7.84 mg/mL (*p-*adj = 0.025) in average IAA production relative to the SDS NIF group (S3 Fig). Taken together, results from these statistical analyses confirmed the tested hypothesis by demonstrating that DSP isolates with the complete set of essential NIF genes produce higher levels of IAA on average compared to the SDS group.

#### Mucilage diazotrophs liberate soluble phosphate from an inorganic mineral complex

Testing *in vitro* cultures of each isolate for the presence of soluble phosphate after incubation with hydroxyapatite provided evidence that mucilage diazotrophs possess the functionality to liberate forms of phosphate contained in complex, inorganic mineral complexes. This presented the case for dual PGP functionality by the majority of mucilage diazotrophs in all three NIF groups that exhibited strong phenotypes in both the BNF and PO_4_ bioassays (Fig 1). Similar to trends observed for median concentration with the bioassay for IAA biosynthesis, the DSN and DSP groups displayed respective PO_4_ concentration medians of 852.76 and 839.47 mg/L while the median for the SDS group was lower at 702.1 mg/L (S3 Table). Unlike results from the IAA assay, outliers among the NIF groups for the PO_4_ solubilization assay with values in the upper quartile above three times the IQR were less frequent and individual isolates of the DSP and DSN groups produced PO_4_ concentration maxima of 2214.7 and 2051.1 mg/L, respectively (S2 Fig D, S3 Table). These results indicated that when considered at the NIF group level, the DSN and DSP isolates showcased higher PO_4_ liberation than did SDS isolates.

Fitting observations of isolate PO_4_ liberation levels to a linear model ([PO_4_] ∼ 1 + NIF group) facilitated testing of the hypothesis that NIF group membership impacted the ability of isolates to solubilize inorganic phosphate. Results from ANOVA affirmed that differences in NIF group assignment significantly contributed to observed variance in phosphate liberation from hydroxyapatite (*p* = 6.721e^-06^). Pairwise comparison of differences in average concentrations of liberated PO_4_ between the NIF groups confirmed that both DSP and DSN isolates displayed significant increases over those comprising the SDS group levels of 175.6 mg/L and 186.9 mg/L (*p-*adj < 0.0001), respectively (S3 Fig). These statistical analyses validated the tested hypothesis by providing a high level of confidence that isolates of the DSN and DSP groups outperform members of the SDS group with respect to PO_4_ solubilization.

### *In planta* inoculations of potato revealed mucilage isolates capable of PGP

Based on the observations that diazotrophic isolates from Sierra Mixe maize possessed genetic markers for multiple PGP traits and demonstrated the ability to carry out the associated PGP functionalities by *in vitro* phenotypic assay, we hypothesized that the isolates had the ability to promote the growth of non-cereal crops. Conducting *in planta* inoculation experiments with clonally propagated potato plantlets to identify isolates capable of mixed PGP confirmed this hypothesis by revealing diazotrophic isolates from diverse genera that demonstrated potato PGP. After screening plants inoculated with mucilage isolates for PGP against mock-inoculated controls, linear modeling of data for fresh and dry plant tissue sample weight along with subsequent hypothesis testing identified 16 diazotrophic isolates that significantly (*p*-adj ≤ 0.01, 19 ≤ N microbe inoculation treatments compared to a single mock-inoculated control ≤ 38) increased accumulation of either the shoot biomass, root biomass, or both (Table 2). Taxonomic classification of the corresponding WGS assemblies demonstrated that the majority of these potato PGP isolates resembled genera from the *Enterobacteriaceae*. However, BCW-200533 and BCW-201835 also promoted the growth of potato were identified to be *Rhodococcus* and *Staphylococcus*, respectively. Enterobacterial diazotrophs from the DSP group that conferred potato PGP included five isolates identified as *Metakosakonia*, *Rahnella*, or *Raoultella*. Conversely, the remaining 11 isolates were identified as *Enterobacter*, *Lelliottia*, *Pantoea*, *Rahnella*, *Rhodococcus*, *Serratia*, and *Staphylococcus* that were representatives of the DSN group (Table 2, S2 Table).

**Table 2.**
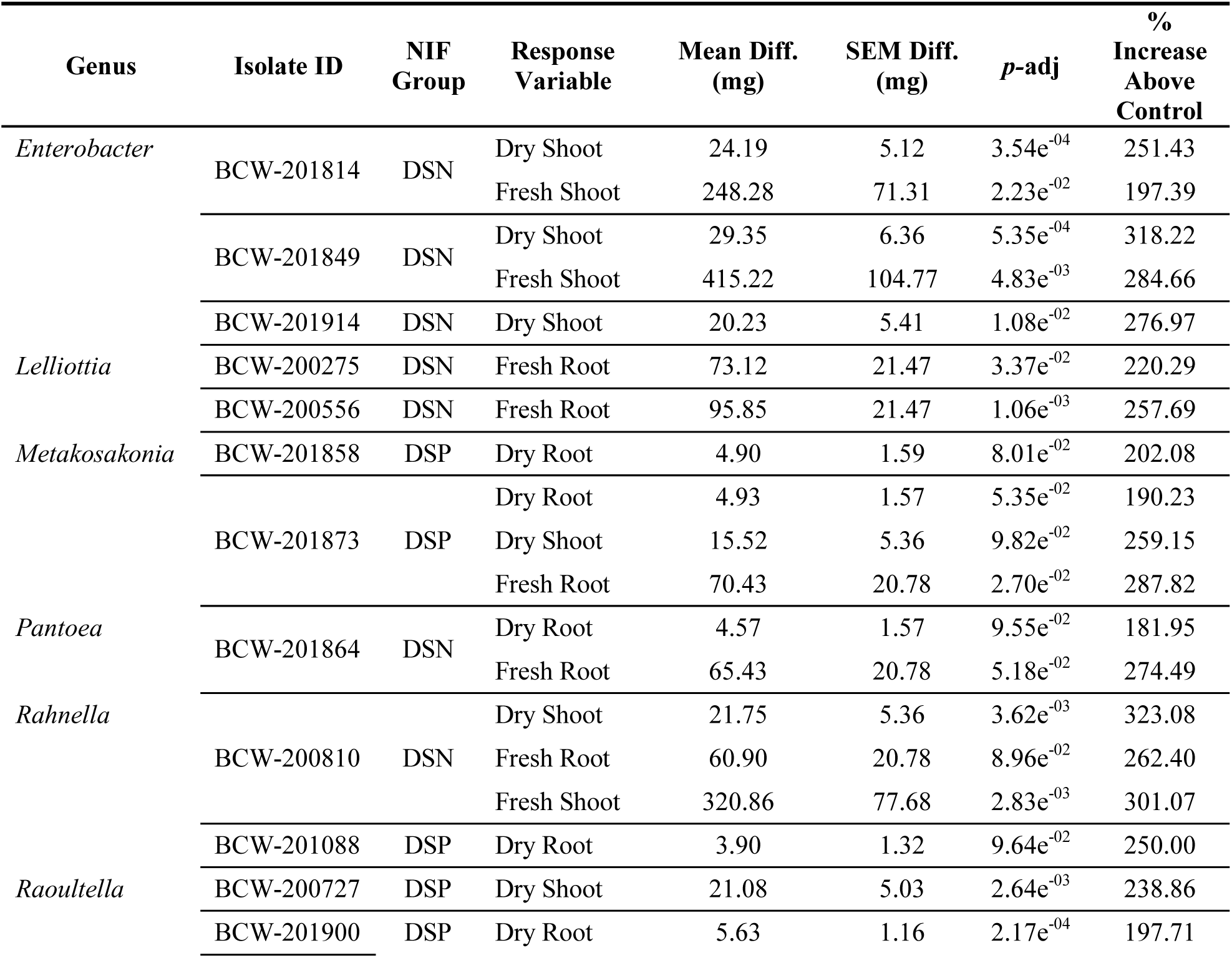
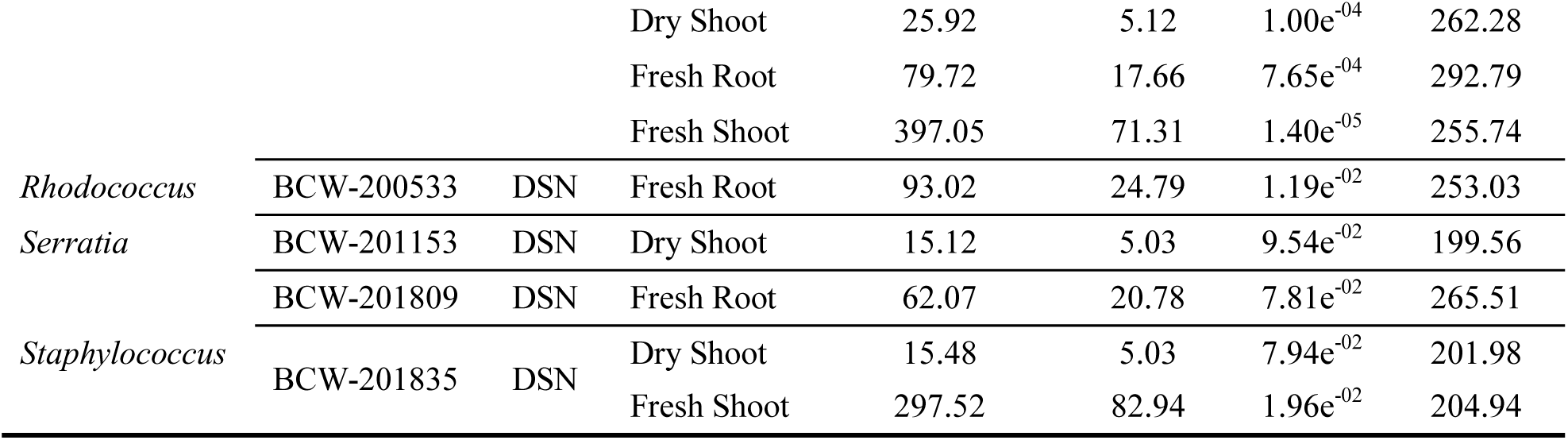
Potato growth responses to in planta inoculation with mucilage isolates. Mucilage isolates were screened in groups with independent controls (*n =* 3, see Methods). Controls consisted of plantlets that were inoculated with 100 µL of a liquid saline solution 0.9% (w/v). Weight differences for each measured response were determined in R by fitting weight measurement data to linear models (weight ∼ 1 + Inoculation Treatment), performing mean estimations and multiple *t*-tests for contrasts between inoculation treatments and the mock-inoculated control. SEM diff. indicates the standard error of mean difference estimates between each inoculation treatment and the mock-inoculated control. Adjusted *p*-values (*p-*adj) correspond to confidence levels for mean difference estimates corrected for multiple testing against single controls using Dunnett’s method. Percent increase above control values indicate the percentage of weight for the designated response variable with respect to corresponding values observed with the mock-inoculated control for each batch of inoculations. Abbreviations for NIF group assignments include Dos Santos Positive (DSP), and Dos Santos Negative (DSN).

Analyzing both fresh and dry shoot weight measurements taken after independent incubation with each mucilage diazotroph identified eight isolates capable of potato PGP. While visualizing average weights for each mono isolate inoculation experiment showed a range of shoot growth responses with respect to mock-inoculated control plants (S4-S16 Fig), linear modeling of the data and testing the hypothesis that observed increases in weight were caused by microbial inoculation identified a subset of isolates with statistically supported difference estimations (*p*-adj ≤ 0.01). This allowed for rejection of the null hypothesis that observed increases were random. Fresh shoot weight differences found 5 mucilage isolates that conferred shoot growth promotion with effect sizes ranging between 200 and 300 % of mock-inoculated controls (Table 2). Following desiccation of each shoot sample, repeating the analysis with dried tissue weights showed 9 diazotrophs with recorded weights significantly higher than those of corresponding control plants. The dried shoot weights for plants receiving these beneficial inoculums had an observed effect size that ranged from 199 to 318 % of controls. Comparison of differences between PGP isolate identification based on analysis of fresh or dry shoot samples showed that 5 of the 8 isolates identified had statistically significant (*p*-adj ≤ 0.01) PGP effects on potato shoot weight for both fresh and dry response variables (*Enterobacter* BCW-201814, *Enterobacter* BCW-201849, *Rahnella* BCW-200810, *Raoultella* BCW-201900 and *Staphylococcus* BCW-201835).

Applying the procedure for statistical analysis used with shoot weight measurements to root weight measurements taken after *in planta* inoculation and incubation confirmed that 10 mucilage isolates had the ability to confer root growth promoting effects in potato. Although visualizing mean estimates and associated standard error for root weights presented a wide range of inoculation effects with numerous degrees of variation (S4 – S16 Fig), further analysis of fresh and dry root weight measurements uncovered statistically supported observations of growth promotion by 8 and 5 different isolates, respectively. Mucilage isolates with potato root PGP phenotypes determined using fresh root weights had effect sizes between 220 and 293 % of mock-inoculated controls, and those observed with dry root samples presented a lower range of 181 – 250 % (Table 2). Among the 10 isolates found to confer potato root PGP, corresponding promotion effects for both fresh and dry weight measurements occurred following inoculation with 3 isolates (*Metakosakonia* BCW-201873, *Pantoea* BCW-201864 and *Raoultella* BCW-201900). Furthermore, three of the 16 isolates determined to confer potato PGP by *in planta* inoculation exhibited potato PGP effects on both shoot and root tissue (*Metakosakonia* BCW-201873, *Rahnella* BCW-200810 and *Raoultella* BCW-201900).

### SynCom inoculation of potato in the greenhouse confirmed mucilage isolate PGP

Inoculating Russet Burbank (RB) and Red La Soda (RLS) potatoes in the greenhouse with synthetic microbial communities (SynComs) comprised of isolates found to increase potato biomass *in planta* provided further evidence for PGP by mucilage diazotrophs in conditions of replete fertilization and nitrogen stress. Selection of SynCom members relied on integration of PGP gene mining results with individual performances on the *in planta* potato PGP inoculation assay. This resulted in a PGPB candidate list consisting of three DSP and three DSN diazotrophic isolates that exhibited both diversity in genotype for targeted PGP genes and the measured capability to promote potato root and/or shoot growth (Fig 2A). Subsequent randomization and nested pooling of these isolates generated six SynComs for greenhouse potato inoculation (Fig 2B). This strategy allowed for increased replication of inoculation with each mucilage isolate and assessment of microbe-microbe interaction effects on potato PGP.

**Fig 2.**
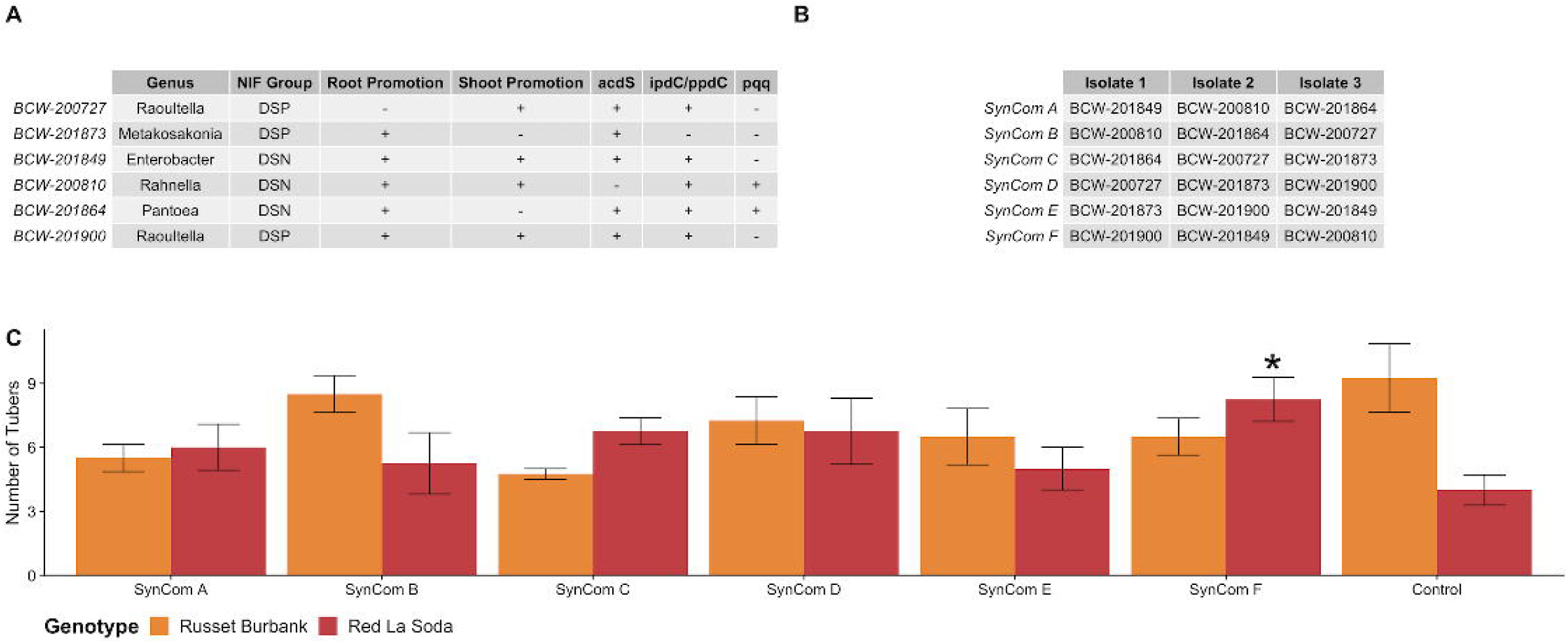
Design and impact of mucilage isolate SynCom inoculations for potato PGP. Factors for incorporating mucilage isolates in the experimental design included strong potato growth promotion phenotypes from the *in planta* inoculation assay along with variation of PGP gene possession among the targeted traits for the collection. A) SynCom isolate selection criteria and specifications that include: percent growth promotion of potato roots (R) and shoots (S) from the *in planta* potato inoculation assay, nitrogen fixation group (NIF Group) assignment based on *nif* gene mining results, and the presence (+) or absence (-) of targeted marker genes for ACC deaminase (*acdS*), biosynthesis of IAA (*ipdC/ppdC*), and genes comprising the PQQ operon (*pqq*). B) Nested pooling design of mucilage isolate SynComs for the greenhouse inoculation experiment with potato. C) Bar plot indicating the average number of tubers produced by potato plants of either the Russet Burbank (RB) or Red La Soda (RLS) genotypes after SynCom inoculation treatment with replete fertilization (*n* = 4). The value annotated with an asterisk (*) indicates a difference estimate determined to be significant after comparison to the mock-inoculated control group (*p*-adj = 0.06).

Although SynCom inoculations of plants grown in the greenhouse demonstrated trends for increased shoot biomass production in both potato varieties relative to mock-inoculated controls that received a sterile Hoagland medium solution, linear modeling of the shoot weight data and hypothesis testing for pairwise comparisons of growth responses to all inoculation treatments found no significant differences. In the case of plants receiving replete fertilization, RB potatoes inoculated with SynComs A, B, C, D, and F had higher average weights for shoot biomass accumulation relative to mock-inoculated controls at differences of 2.3, 9.3, 6.8, 12 and 16.8 grams, respectively (S17 Fig A). Inoculation of RLS plants also treated with complete fertilizer experienced respective gains in average shoot weight over mock-inoculated controls at levels of 2.5, 9.7, 8.5 and 4.3 grams for SynComs B, D, E and F (S17 Fig A). While SynComs A and C did not produce an increase in average shoot weight for potatoes of both varieties that experienced nitrogen stress, inoculation with SynComs D, E and F increased the average shoot weight of stressed RB (5.2, 5.8, 2.3 grams, respectively) and RLS plants (9.7, 8.5, 4.3 grams, respectively) relative to mock-inoculated controls (S17 Fig D). Despite the high level of variance among the shoot weight measurements in the experiment, these results indicated that mucilage isolates have the potential to increase the shoot biomass of potato plants based on the mean shoot weight difference estimates observed between SynCom treated groups and the mock-inoculated control.

Implementing statistical analysis of root weight measurements also yielded non-significant mean difference estimates between inoculated and mock-inoculated groups in both potato varieties. However, the observed average root weights over the replications also demonstrated a trend for root PGP as a result of SynCom treatment. Inoculated RB plants receiving replete fertilization and treatment with SynComs D and F displayed root weight averages that were 133 and 123% of that observed with the control group, respectively (S17 Fig B). Likewise, RB plants under nitrogen stress treated with SynComs E and F had mean root weights measured to be 130 and 124% of the control group average (S17 Fig E). RLS plants receiving replete fertilizer treatment and inoculation with SynComs C, D and F also produced greater average root weights relative to the control group at respective percentages of 125, 120, and 126%. SynComs A and F also generated increases for RLS average root weight at 128 and 134% of the observed control. Surprisingly, the results for each of these factorial conditions demonstrated the commonality of SynCom F inoculation producing an observed increase in average root weight.

Growing potatoes in the greenhouse provided the ability to assess the effect of SynCom inoculation on tuber production. Despite performing calculations that resulted in non-significant mean tuber weight difference estimates for inoculated groups against the mock-inoculated control, SynComs A, D and F exhibited the trend of producing increased average tuber weights for multiple factorial conditions in the experiment. Under replete and deficient fertilization conditions, inoculation of RLS potatoes with SynCom A resulted in positive mean tuber weight differences relative to the control groups at levels of 29 and 23 grams, respectively (S17 Fig C, S17 Fig F). However, inoculation with both SynComs D and F only produced average tuber weight increases relative to the control under replete fertilization. SynCom D inoculation generated respective increases to average tuber weight in RB and RLS potatoes by 24 and 44 grams. In a similar fashion, SynCom F inoculation also yielded increased average tuber weights relative to the control with gains of 26 and 36 grams, respectively. These observed trends for increased average tuber weight over the mock-inoculated control as a result of SynCom inoculation provided additional evidence for potato PGP by isolates contained within SynComs A, D and F.

In addition to measuring average tuber weight, evaluating the number of tubers set by each plant after microbial inoculation and differential fertilization treatments revealed that some mucilage isolates had an impact on tuber development in potatoes. Regarding the tuber number for RB plants that received replete fertilization, SynCom inoculation treatment was not found to increase average tuber number (S17 Fig G). However, measurements for RB potatoes receiving fertilizer stress and inoculation with SynCom E produced a mean difference in average tuber number of 1.75 with respect to the mock-inoculated control (S17 Fig H). With respect to inoculation of nutrient stressed RLS plants, all SynCom treatments did not increase average tuber number relative to the mock-inoculated control. Interestingly, the combination of SynCom inoculation, replete fertilization, and the RLS genotype revealed the trend of increased tuber number relative to the mock-inoculated control (Fig 2C). While increases in tuber number for RLS potatoes receiving proper nutrition occurred for all SynCom inoculated groups, SynCom F produced the largest effect size with a significant difference of 4.25, amounting to 206% of the control (*p*-adj = 0.06).

The observed trend of treatment with SynCom F resulting in increases to potato growth warranted further investigation of its individual components. Deconstruction of SynCom F provided the means to assess the relative contribution of each mucilage diazotroph composing the original inoculation treatment. Adherent to the previously imposed differential fertilization treatments (replete or nutrient stressed) and greenhouse conditions, a subsequent experiment with independent inoculations of RLS potato plants with BCW-200810 (*Rahnella* sp.), BCW-201849 (*Enterobacter* sp.) and BCW-201900 (*Raoultella* sp.) revealed several PGP capabilities for diazotrophic isolates of SynCom F when compared to a mock-inoculated control group.

Inoculation of RLS potatoes in the greenhouse with individual components from SynCom F resulted in significant increase of average shoot and root weights relative to mock-inoculated controls (*p*-adj ≤ 0.05). While the estimates for the average shoot weight of RLS potatoes under replete fertilization treatments demonstrated consistent levels across the inoculated and mock-inoculated groups, plants subjected to fertilizer stress that received inoculation with BCW-200810, BCW-201849 and BCW-201900 showcased respective averages determined to be 125, 133 and 144% of the mock-inoculated control. Among these differences, mean estimates for shoot weight between control plants and those inoculated with BCW-201849 and BCW-201900 were significantly different (adjusted *p*-values of 0.03 and 0.01, respectively) (Fig 3B). Growing RLS potatoes in the presence of all three mucilage isolates from SynCom F produced increases to average root biomass across both fertilization treatments as well. Despite showing mean root weight difference estimates from the mock-inoculated plants that were not significantly different, BCW-200810 inoculated plants experiencing replete fertilizer showcased an average root biomass increase of 47% and those under nutrient stress yielded a 33% improvement (Fig 3C, Fig 3D). Comparatively, confidence levels for increases in average root growth in response to inoculation with BCW-201849 and BCW-201900 under both levels of fertilization were significantly higher. Notably, BCW-201849 and BCW-201900 inoculated plants generated respective differences to mock-inoculated plants of 26.2 (188% of control, *p*-adj = 0.04) and 25.2 (185% of control, *p*-adj = 0.05) grams under replete fertilizer treatment. RLS plants inoculated with these two isolates also presented increases in average root weight when grown under fertilizer stress, where BCW-201849 and BCW-201900 had increased respective averages of 8 and 7.8 grams (160% of control, *p*-adj = 0.1).

**Fig 3.**
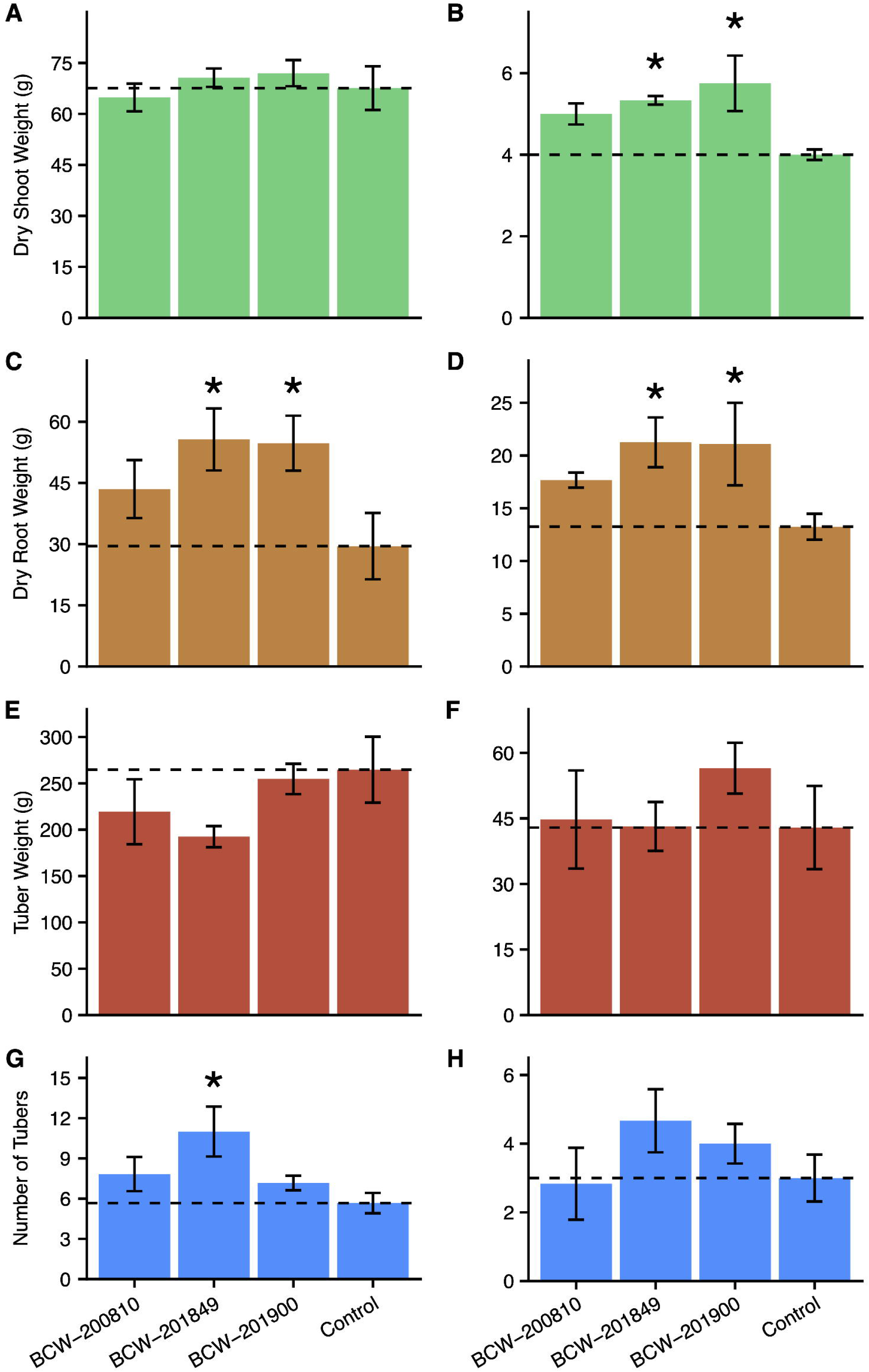
Growth responses for potato plants inoculated with individual isolates of SynCom F. Results from the destructive sampling of Red La Soda potatoes grown either with replete fertilization or nutrient deficiency, and inoculation with one of four treatments: inoculation with *Rahnella* BCW-200810, *Enterobacter* BCW-201849, *Raoultella* BCW-201900, or a mock-inoculation of 0.5X Hoagland liquid medium without the presence of cultured bacterial cells (control). Response variables of inoculated plants grown under replete fertilizer conditions included dry shoot weight (A), dry root weight (C), tuber weight (E), and number of tubers (G). Measurements for inoculated plants subjected to nutrient stress also included dry shoot weight (B), dry root weight (D), tuber weight (F), and number of tubers (H). Bars indicate standard error of the mean (SEM) recorded weight value for each response variable (*n* = 6). Values annotated by an asterisk (*) had mean difference estimates relative to mock-inoculated controls with Dunnett adjusted *p*-values < 0.05 (with the exception of panel D, where *p*-adj = 0.1).

Observed results from inoculating RLS potatoes with individual isolates from SynCom F also included a small impact on average tuber weight under nutrient stress and a trend for increased number of tubers. The average tuber weight responses for inoculated plants grown without fertilizer stress showed no signs of growth promotion relative to those that received mock-inoculation treatment (Fig 3E). RLS plants inoculated with BCW-201900 resulted in an average tuber weight increase of 13.6 grams in comparison to the mock-inoculated control, but hypothesis testing with this result provided a low degree of confidence in the estimate (Fig 3F). In congruence with results from inoculation with SynCom F in the preceding experiment, BCW-201849 inoculation of RLS potatoes grown with replete fertilizer increased the average number of tubers developed by 94% relative to the mock-inoculated control group (Fig 3G, *p*-adj = 0.02). Additionally, BCW-201849 and BCW-201900 inoculation groups presented a similar trend by increasing the tuber number of RLS plants that experienced fertilizer stress by comparison to plants that did not receive microbial inoculation treatments (Fig 3H).

### Mucilage derived *Raoultella* conferred PGP to conventional maize under nitrogen stress

Given the demonstrated consistency of the diazotrophic mucilage isolate BCW-201900 (*Raoultella* sp.) in its ability to test positive for PGP genotypes, *in vitro* assays for PGP phenotypes, promotion of potato growth *in planta*, and exhibit potato PGP in the greenhouse, we decided to assess its ability to confer PGP in conventional maize. This prompted investigation of the ability for BCW-201900 to confer PGP upon varieties of maize distantly related to the Sierra Mixe landrace. Using a greenhouse setting similar to that of the inoculation experiments conducted with potato, comparison of BCW-201900 inoculated B73 x Mo17 conventional maize hybrids with mock-inoculation control plants demonstrated the mucilage isolate’s ability to modulate maize development under severe nitrogen stress (15 ppm total N delivered per feeding). Specifically, BCW-201900 inoculated plants had accelerated shoot development and anthesis in response to the absence of adequate nitrogen fertilization compared to B73 x Mo17 plants that received mock-inoculation treatment (Fig 4A). Weekly growth monitoring also revealed that BCW-201900 inoculated maize plants experienced progressive growth acceleration when compared to mock-inoculated controls. Statistical analysis of average height measurements for inoculated vs control groups at each sampling time revealed significant differences for height increases when inoculated with BCW-201900. Notably, inoculated plants exhibited average height increases of 20.5 cm at 4 weeks (*p*-adj < 0.01), 21.6 cm at 7 weeks (*p*-adj < 0.01), and 60.4 cm after 8 weeks (*p*-adj < 0.0001) in the greenhouse (Fig 4B).

**Fig 4.**
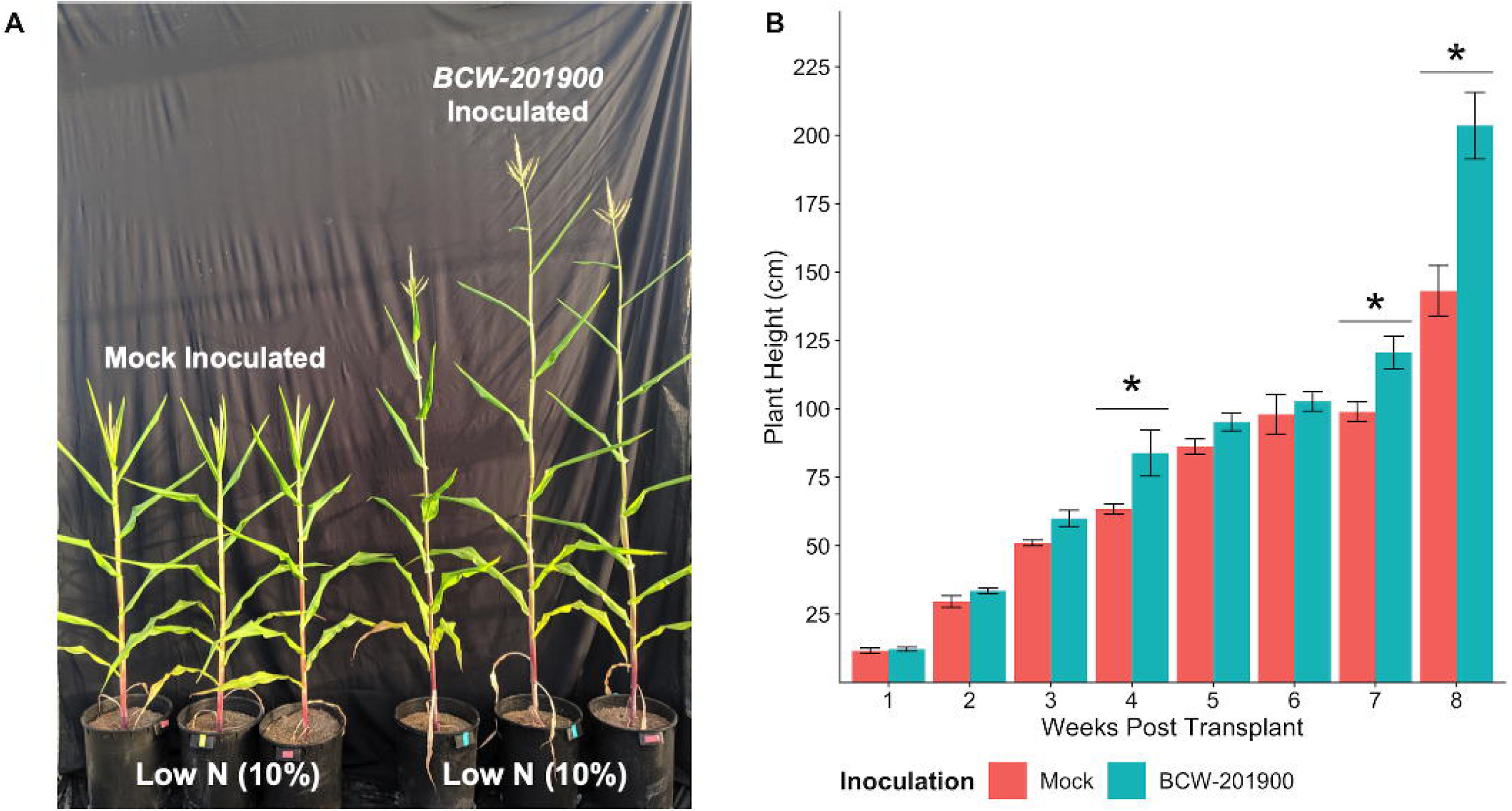
Inoculation of conventional maize with *Raoultella* isolated from Sierra Mixe maize. Results from inoculating plants of the B73 x Mo17 conventional maize variety with *Raoultella* isolate BCW-201900 and subjecting them to severe nitrogen stress (15 ppm N delivered via fertilized irrigation). Plants were grown alongside mock-inoculated control plants of the same variety (*n* = 4). A) Plants that received mock inoculation (left) or BCW-201900 inoculation (right) were photographed after 60 days in the greenhouse. B) Average plant height of BCW-201900 inoculated and mock-inoculated control groups. Measurements were taken on a weekly basis. Data points were subjected to linear modeling and hypothesis testing in R using the emmeans 1.4.1 package. The *p*-values associated with significant differences were adjusted using the Tukey HSD method. Significant differences for comparisons denoted by asterisks (*) indicate *p*-adj < 0.01.

## Discussion

### Mucilage diazotrophs exhibited alternative PGP traits independent of NIF group

Characterizing the PGP functionalities of diazotrophic bacteria isolated from Sierra Mixe maize relied on the integration of results from WGS analysis and confirmation of PGP phenotypes to provide multiple points of support for the targeted PGP traits. Implementing this strategy was inspired by the high level of genomic and functional evidence previously generated to establish isolates within the collection as diazotrophic [11], as well as our hypothesis that mucilage isolates possessed the ability to confer PGP functionalities in addition to BNF. Identifying homologous coding sequences to HMMs of genes associated with direct PGP mechanisms within the genomes of mucilage diazotrophs provided a base level of support for determining the extent of each isolate’s PGP capabilities and warranted further investigation into their corresponding PGP phenotypes. Accordingly, this study used *in vitro* biochemical assays to quantify isolate activity levels for PGP mechanisms alternative to BNF that included ACC deaminase activity, biosynthesis of IAA, and PO_4_ solubilization, which consequently verified many of the alternative PGP genotypes assigned to mucilage diazotrophs. However, statistical analysis of results from the ^15^N incorporation assay and all three colorimetric PGP assays indicated significant differences in the average capacity for isolates comprising each NIF group to perform the respective PGP functionalities *in vitro* (S3 Fig). This suggests that *nif* gene content may have an impact on the extent to which diazotrophs from different NIF groups are capable of conferring each of the investigated PGP traits in a plant environment.

Elucidating the genomic presence of the genetic marker for ACC deaminase activity in diazotrophic isolates, as well as their associated phenotypes, revealed an abundance of this PGP trait within the colleciton among all 3 NIF groups. ACC is synthesized by the plant enzyme ACC synthase as a metabolic precursor to the plant growth regulating compound (PGR) ethylene that is involved in responses to abiotic and biotic stress in the environment [3]. Plant associated bacteria with the *acdS* gene faciltate PGP by uptake and subsequent deamination of ACC through action of ACC deaminase to form ammonia and ⍰-ketobutyrate, which in turn lowers inhibition of plant growth by ethylene [35]. Drawing motivation from this well established model for ACC deaminase mediated PGP, this study found that the majority of mucilage diazotrophs possessed homologous coding sequences to the *acdS* gene (S1 Fig, S1 Table), and the ability to grow on defined media supplemented with ACC as the independent, limiting source of bio-available nitrogen irrespective to their NIF group assignment (Fig 1, S2 Fig). While possession of homologs for the *acdS* gene was confirmed in isolates classified to roughly two thirds of identified genera within the collection (Table 1), many of the ACC deaminase possessing isolates classified to the same genus had different NIF group membership assignments (S2 Table) based on their respective *nif* gene content (e.g. *Enterobacter, Pseudomonas,* and *Rahnella* isolates). This suggests that mucilage diazotrophs are capable of facilitating PGP through ACC deaminase activity regardless of their designated BNF mechanism.

Examination of isolate genomes for the possession of homologous coding sequences to the *ipdC/ppdC* HMM and ensuing colorimetric assay uncovered that the majority of mucilage diazotrophs exhibit the genetic capacity necessary for IAA biosynthesis. Bacteria that associate with plants in both beneficial and pathogenic contexts are known to employ different metabolic strategies for IAA production [36]. However, previous investigations suggested that utilization of Indole-3-Pyruvate Decarboxylase (*ipdC*) to produce IAA is associated with PGPB rather than phytopathogens via utilization of Trp [37]. In recognition of this distinction, genomic mining of mucilage isolates using *ipdC* as a marker for IAA mediated PGP revealed that all isolates possessed homologous sequences for the gene (S1 Fig, S1 Table). Pairing this discovery with results from the colorimetric assay to quantify IAA production that confirmed isolate production of IAA (Fig 1, S2 Fig, S3 Table) strengthened the assertions that mucilage diazotrophs are capable of facilitating this direct form of PGP, and that the presence of this feature occurred independent of NIF group assignment. Furthermore, when paired with the high abundance of diazotrophs found to display ACC deaminase functionality, the ubiquity of IAA biosynthesis characteristics among mucilage diazotrophs supports designation of the isolate collection as a resource to develop bio-stimulants based on the reported synergism of these two traits for conferring PGP [38].

Surveying mucilage diazotrophs for the possession of genes related to phosphate solubilization and testing their ability to liberate phosphate from a complex, inorganic source provided evidence for a third mode of alternative PGP among isolates of all NIF groups in the collection. Previous descriptions of gluconic acid production by rhizobacteria using the membrane bound holoenzyme PQQ-glucose dehydrogenase (PQQ-DH) as a PGP mechanisms for phosphate solubilization in soil environments [5, 39] prompted mucilage isolate genome mining for protein coding sequences homologous to the associated HMMs and subsequent phenotypic assay. Although the vast majority of diazotrophic isolates had sequences matching the model for PQQ-DH, the lack of abundance in sequences similar to HMMs for genes of the PQQ operon suggested that the majority of mucilage diazotrophs are unlikely to utilize the PQQ-dependent mechanism for phosphate solubilization (S1 Fig, Table 1, S1 Table). This observation was further supported by results from the phosphate solubilization assay, which demonstrated that the mucilage isolates from all three NIF groups were shown to be capable of solubilizing phosphate after being incubated in phosphate-deficient medium containing hydroxyapatite (Fig 1, S2 Fig, S2 Table). While these results validate the presence of mucilage diazotrophs capable of solubilizing phosphate using PQQ-DH in the isolate collection and support their potential to modulate plant growth through via this mechanism, they also invite the likely possibility that many of the isolates employ alternative strategies for phosphate liberation, such as the secretion of organic acids other than gluconic acid [40].

### A subset of mucilage diazotrophs are versatile PGPB lacking host specificity

Considering the isolated region of Sierra Mixe maize cultivation, the broad range of genomic diversity observed within the mucilage diazotroph collection [11], and the confirmation of PGP phenotypes in addition to BNF, we hypothesized that mucilage diazotrophs were capable of conferring PGP to crop systems highly divergent from maize. The use of clonally propagated potato plants was well suited for addressing the hypothesis based on its reduced likelihood of contributing plant genotype impact on measured response variables, and the high fertilizer demand of potato in agricultural production [41]. In accordance with these justifications, the potato inoculation experiments described in this study served the dual purpose of confirming that mucilage diazotrophs are capable of PGP and testing their ability to benefit crop systems distant from their source of isolation.

Unexpectedly, *in planta* inoculation of potato with mucilage diazotrophs provided baseline support for our hypothesis by demonstrating that both DSP and DSN isolates enhanced shoot and root biomass accumulation. The confirmation of PGP phenotypes for DSP isolates classified as *Metakosakonia* and *Raoultella* through observed increases in shoot and root growth of potato (Table 2, S2 Table) suggested their potential to be versatile bio-stimulants for crop production. Concurrently, the abundance of DSN isolates shown to enhance biomass accumulation indicated that possession of the essential *nif* genes was not a requirement for the growth benefits observed with the inoculated plants, and that these isolates likely employed alternative mechanisms to confer potato PGP. Although results from the *in planta* inoculation assay elucidated that mucilage diazotrophs have the capacity to enhance potato development under highly controlled settings, the discovery of DSP and DSN isolates with established phenotypes for multiple forms of direct PGP (S1 Table) validated the chosen strategy for identification of PGPB within the collection.

While establishing *in planta* PGP phenotypes provided a proof of concept for using the diazotrophic isolate collection as a resource to develop crop bio-stimulants, performing potato inoculation experiments in the greenhouse with SynComs comprised by potato PGP isolates increased the level of support for the hypothesis that these isolates function as PGPB of crops distantly related to maize. Designing SynComs with DSP and DSN isolates found both to dramatically increase potato growth *in planta and* possess marker genes for multiple PGP functionalities increased the likelihood of observing potato PGP in a greenhouse setting (Fig 2). Despite high variance among the replications, the observed trends of increased shoot, root and tuber weights for SynCom inoculated plants relative to mock-inoculated controls (S17 Fig) demonstrated that potato PGP conferred by mucilage diazotrophs is feasible beyond the laboratory. The unexpected doubling in the number of tubers formed by RLS potatoes inoculated with SynCom F highlighted the candidacy of isolates comprising the treatment as PGPB for bio-stimulant development. Further dissection of SynCom F revealed definitive potato PGPB from the mucilage isolate collection through independent inoculation of RLS potato with either an *Enterobacter* isolate from the DSN group (BCW-201849), or a *Raoultella* isolate from the DSP group (BCW-201900). Mono-isolate inoculations of RLS potato plants in the greenhouse with these two mucilage isolates led to significant increases in shoot and root biomass weights relative to mock-inoculated control plants (Fig 3) that were not previously observed after the SynCom F inoculation experiment (Fig S17). This suggested that mono-isolate inoculation of RLS potatoes with BCW-201849 and BCW-201900 was a more efficient PGP strategy than co-inoculation of the isolates as a SynCom. Furthermore, the discovery of BCW-201849 complements the report of enhanced root and shoot growth as a result of inoculating potato plants with *Enterobacter* under a field setting [41], with the exception of the mucilage associated *Enterobacter* isolate described here being identified as the causal factor for tuber doubling (Fig 2C, Fig 3G).

### Mucilage derived *Raoultella* demonstrated versatile PGP capability

Previous investigations of bacteria classified to the genus of free living, gram-negative enterobacteria termed *Raoultella* characterized its known association with agricultural crops. After becoming distinguished from the neighboring genus *Klebsiella* through a combination of genomic and biochemical studies [42], reports on plant associations with *Raoultella* spp. began to surface. Through the rise of applying WGS for bacterial isolates from environmental samples, detailed genomic information became available for *Raoultella ornithinolytica* isolated from soil, *Raoultella planticola* isolated from river water, and the diazotrophic endophyte *Raoultella terrigena* isolated from *Nicotiana tabacum* [43–45]. With respect to diazotrophic associations with cereal crops, recent investigation into a diazotrophic strain of *R. planticola* demonstrated that it was capable of conferring BNF-mediated PGP to sugar cane through endophytic colonization [46]. While reference genomes for these species exist in GenBank [47] and the Genome Taxonomy Database (GTDB) [48] used to classify mucilage isolate genomes in this study, the lack of species level assignment to described species of *Raoultella* for many mucilage derived *Raoultella* isolates using WGS similarity indicates the potential genomic novelty for *Raoultella* associated with Sierra Mixe maize and suggests that strain variation is important for these traits to be conferred in the microbiome.

This study characterized the diazotrophic isolate *Raoultella* BCW-201900 as a bacterium capable of multiple PGP functionalities, a growth promoter of potato through *in planta* inoculation, a common element of inoculation treatments that enhanced greenhouse potato growth either in SynComs or independently, and as a modulator of conventional maize development. In addition to its established BNF phenotype and assignment to the DSP NIF group, BCW-201900 also possessed genomic features and generated phenotypes for ACC deaminase activity and IAA production (Fig 1). Among the 16 mucilage isolates determined to be significant potato growth promoters following *in planta* inoculation, BCW-201900 was the only mucilage isolate that increased biomass production with high levels of confidence in all four of the measured response variables (Table 2). Pairing this with its demonstrated ability to enhance the growth of potato shoots, roots and tuber development in the greenhouse under nitrogen stress bolstered the assertion that BCW-201900 is a flexible PGPB capable of conferring PGP to dicotyledonous plants (Fig 3). Further testimony to the adaptable nature of PGP by BCW-201900 arose from observations that BCW-201900 accelerated the development of conventional maize under extreme nitrogen stress (Fig 4). As an added measure of PGPB versatility to the multiple PGP functionalities and growth promotion phenotypes described here, the BCW-201900 genome also possesses homologous sequences for alternative *nif* genes in addition to the essential *nif* operons comprising the Dos Santos model [10, 11]. These revelations demonstrate the consistency of BCW-201900 as a mucilage derived PGPB and support its candidacy for the development of a versatile bio-stimulant in the future.

## Conclusions

These experimental results confirm the two-stage hypothesis that diazotrophic bacteria isolated from aerial root mucilage possess the ability to carry out multiple methods of direct PGP in addition to BNF that result in growth promotion of crops unrelated to Sierra Mixe maize. We identified that the majority of the diazotrophic isolates possess marker genes for ACC deaminase, IAA production and PO_4_ solubilization in addition to *nif* genes and presented evidence of their ability to confer these functionalities *in vitro* at variable capacities that were found to be significantly affected by *nif* gene content. Utilizing these mucilage diazotrophs as inoculants for potato plants and a conventional hybrid variety of maize also confirmed the ability of several isolates to confer PGP phenotypes for crops outside of their native environment. This suggests the potential for mucilage diazotrophs to function as PGPB over a broad host range, as evidenced by our reported discovery of multiple PGP functionalities conferred by *Raoultella* BCW-201900. Overall, our investigation outlined the PGP capabilities presented by this unique collection of diazotrophic bacteria from Sierra Mixe maize, emphasized its potential utility for the development of novel bio-stimulants, and ultimately provides support for the sustainable intensification of agricultural production.

## Acknowledgements

Christopher Durand, Andrew Hutchison and associated staff of the U.C. Davis Department of Plant Sciences CORE greenhouse team were highly involved in setting up and maintaining greenhouse inoculation experiments. We also thank Professor Allen Van Deynze, Director of the Seed Biotechnology Center at U.C. Davis, for his input during the design of greenhouse inoculation experiments. U.C. Davis undergraduate students who assisted in the collection of samples during inoculation experiments included Cleo Fang and Alexander Vavan.

## Supporting Information

**S1 Fig. Dendrogram of diazotroph genomes with heatmaps of non-*nif* PGP gene profiles.** Marker genes for non-*nif* PGP functionalities were detected by scanning total amino sequences from each pure isolate genome that were predicted using Prokka against HMMs obtained from the TIGRFAM database [15]. Query sequences were considered as positive matches to the targeted HMMs if model coverage was greater than or equal to 75 % with an e-value less than or equal to 1e^-9^. HMMs for marker genes corresponding to the non-*nif* PGP traits of interest included the *acdS* gene encoding ACC deaminase, the *ipdC/ppdC* genes encoding indole pyruvate decarboxylase, individual models for the *pqqBCDEF* genes that encode corresponding subunits of the pyrroloquinoline quinone cofactor and the *pqq* associated dehydrogenase (*pqq-DH*). The non-*nif* PGP gene profiles for each pure isolate are presented in the context of a hierarchically clustered dendrogram corresponding to MinHash distances computed with Sourmash [16].

**S2 Fig. Distribution of PGP assay data from mucilage isolates across NIF groups.** Data acquired from *in vitro* assays to assess mucilage isolate phenotypes for targeted PGP traits was analyzed and plotted using R 3.5.1. Datapoints for the assays represented the average response values observed over three biological replications with each isolate. Points annotated with BCW-ID numbers indicate outliers for each assay. Outliers were determined as isolates with observed values greater than subtracting or adding 3 times the calculated interquartile range from the first or third quartile, respectively. Boxplots were made using tidyverse 1.2.1 [18], ggrepel 0.5 and cowplot 1.0.0. Code for the analysis is hosted on Github at: (https://github.com/shigdon/R-Mucilage-isolate-pgp-assay).

**S3 Fig. Pairwise comparisons of average PGP assay performances by NIF group.** Data obtained from *in vitro* phenotypic assays for PGP functionalities were fit to linear models to calculate mean estimates for each NIF group, compute mean difference estimates between NIF groups, and conduct multiple pairwise comparisons between the estimated means to determine statistical significance of the differences. The three NIF groups include Dos Santos Positive (DSP) isolates possessing all six essential *nif* genes, Semi-Dos Santos (SDS) isolates possessing an incomplete set of the six essential *nif* genes, and the Dos Santos Negative (DSN) isolates without any of the six essential *nif* genes. Each x-axis presents the estimated average response by the respective linear model and the y-axes depict the *p*-values of each comparison that were adjusted by the Tukey HSD method using the R package emmeans 1.4.1. A) Average BNF ratios by NIF group corresponding to the ^15^N incorporation metabolomic assay; B) Average relative growth rate by NIF group for isolates cultured with 1-amino-1-cyclopropane carboxylic acid provided as the nitrogen source; C) Average production levels of indole-3-acetic acid by NIF group; D) Average liberation of soluble phosphate from hydroxyapatite by NIF group. Code for the analysis is hosted on Github at: (https://github.com/shigdon/R-Mucilage-isolate-pgp-assay).

**S4 Fig. Biomass weights for in planta inoculation screen of mucilage isolate set 1.** Mono-isolate inoculation of potato plantlets with 38 mucilage diazotrophs were compared against a single mock-inoculated control. Each figure panel shows average values over triplicate sampling for the following response variables: A) Fresh shoot weight, B) Fresh Root Weight, C) Dry Shoot Weight, and D) Dry Root Weight. Dashed horizontal lines indicate the average measurement observed for the respective mock-inoculated control groups. Isolate numbers along the x-axis correspond to BCW-isolate identification numbers seen in other plots and tables. Numeric annotations in some plots indicate percentage of biomass for selected isolates relative to mock-inoculated controls.

**S5 Fig. Biomass weights for *in planta* inoculation screen of mucilage isolate set 2.** Mono-isolate inoculation of potato plantlets with 30 mucilage diazotrophs were compared against a single mock-inoculated control. Each figure panel shows average values over triplicate sampling for the following response variables: A) Fresh shoot weight, B) Fresh Root Weight, C) Dry Shoot Weight, and D) Dry Root Weight. Dashed horizontal lines indicate the average measurement observed for the respective mock-inoculated control groups. Isolate numbers along the x-axis correspond to BCW-isolate identification numbers seen in other plots and tables. Numeric annotations in some plots indicate percentage of biomass for selected isolates relative to mock-inoculated controls.

**S6 Fig. Biomass weights for *in planta* inoculation screen of mucilage isolate set 3.** Mono-isolate inoculation of potato plantlets with 31 mucilage diazotrophs were compared against a single mock-inoculated control. Each figure panel shows average values over triplicate sampling for the following response variables: A) Fresh shoot weight, B) Fresh Root Weight, C) Dry Shoot Weight, and D) Dry Root Weight. Dashed horizontal lines indicate the average measurement observed for the respective mock-inoculated control groups. Isolate numbers along the x-axis correspond to BCW-isolate identification numbers seen in other plots and tables. Numeric annotations in some plots indicate percentage of biomass for selected isolates relative to mock-inoculated controls.

**S7 Fig. Biomass weights for *in planta* inoculation screen of mucilage isolate set 4.** Mono-isolate inoculation of potato plantlets with 24 mucilage diazotrophs were compared against a single mock-inoculated control. Each figure panel shows average values over triplicate sampling for the following response variables: A) Fresh shoot weight, B) Fresh Root Weight, C) Dry Shoot Weight, and D) Dry Root Weight. Dashed horizontal lines indicate the average measurement observed for the respective mock-inoculated control groups. Isolate numbers along the x-axis correspond to BCW-isolate identification numbers seen in other plots and tables. Numeric annotations in some plots indicate percentage of biomass for selected isolates relative to mock-inoculated controls.

**S8 Fig. Biomass weights for *in planta* inoculation screen of mucilage isolate set 5.** Mono-isolate inoculation of potato plantlets with 35 mucilage diazotrophs were compared against a single mock-inoculated control. Each figure panel shows average values over triplicate sampling for the following response variables: Panels for all figures show average values over triplicate sampling for the following response variables: A) Fresh shoot weight, B) Fresh Root Weight, C) Dry Shoot Weight, and D) Dry Root Weight. Dashed horizontal lines indicate the average measurement observed for the respective mock-inoculated control groups. Isolate numbers along the x-axis correspond to BCW-isolate identification numbers seen in other plots and tables. Numeric annotations in some plots indicate percentage of biomass for selected isolates relative to mock-inoculated controls.

**S9 Fig. Biomass weights for *in planta* inoculation screen of mucilage isolate set 6.** Mono-isolate inoculation of potato plantlets with 35 mucilage diazotrophs were compared against a single mock-inoculated control. Each figure panel shows average values over triplicate sampling for the following response variables: A) Fresh shoot weight, B) Fresh Root Weight, C) Dry Shoot Weight, and D) Dry Root Weight. Dashed horizontal lines indicate the average measurement observed for the respective mock-inoculated control groups. Isolate numbers along the x-axis correspond to BCW-isolate identification numbers seen in other plots and tables. Numeric annotations in some plots indicate percentage of biomass for selected isolates relative to mock-inoculated controls.

**S10 Fig. Biomass weights for *in planta* inoculation screen of mucilage isolate set 7.** Mono-isolate inoculation of potato plantlets with 23 mucilage diazotrophs were compared against a single mock-inoculated control. Each figure panel shows average values over triplicate sampling for the following response variables: A) Fresh shoot weight, B) Fresh Root Weight, C) Dry Shoot Weight, and D) Dry Root Weight. Dashed horizontal lines indicate the average measurement observed for the respective mock-inoculated control groups. Isolate numbers along the x-axis correspond to BCW-isolate identification numbers seen in other plots and tables. Numeric annotations in some plots indicate percentage of biomass for selected isolates relative to mock-inoculated controls.

**S11 Fig. Biomass weights for *in planta* inoculation screen of mucilage isolate set 8.** Mono-isolate inoculation of potato plantlets with 36 mucilage diazotrophs were compared against a single mock-inoculated control. Each figure panel shows average values over triplicate sampling for the following response variables: A) Fresh shoot weight, B) Fresh Root Weight, C) Dry Shoot Weight, and D) Dry Root Weight. Dashed horizontal lines indicate the average measurement observed for the respective mock-inoculated control groups. Isolate numbers along the x-axis correspond to BCW-isolate identification numbers seen in other plots and tables. Numeric annotations in some plots indicate percentage of biomass for selected isolates relative to mock-inoculated controls.

**S12 Fig. Biomass weights for *in planta* inoculation screen of mucilage isolate set 9.** Mono-isolate inoculation of potato plantlets with 35 mucilage diazotrophs were compared against a single mock-inoculated control. Each figure panel shows average values over triplicate sampling for the following response variables: A) Fresh shoot weight, B) Fresh Root Weight, C) Dry Shoot Weight, and D) Dry Root Weight. Dashed horizontal lines indicate the average measurement observed for the respective mock-inoculated control groups. Isolate numbers along the x-axis correspond to BCW-isolate identification numbers seen in other plots and tables. Numeric annotations in some plots indicate percentage of biomass for selected isolates relative to mock-inoculated controls.

**S13 Fig. Biomass weights for *in planta* inoculation screen of mucilage isolate set 10.** Mono-isolate inoculation of potato plantlets with 19 mucilage diazotrophs were compared against a single mock-inoculated control. Each figure panel shows average values over triplicate sampling for the following response variables: A) Fresh shoot weight, B) Fresh Root Weight, C) Dry Shoot Weight, and D) Dry Root Weight. Dashed horizontal lines indicate the average measurement observed for the respective mock-inoculated control groups. Isolate numbers along the x-axis correspond to BCW-isolate identification numbers seen in other plots and tables. Numeric annotations in some plots indicate percentage of biomass for selected isolates relative to mock-inoculated controls.

**S14 Fig. Biomass weights for *in planta* inoculation screen of mucilage isolate set 11.** Mono-isolate inoculation of potato plantlets with 36 mucilage diazotrophs were compared against a single mock-inoculated control. Each figure panel shows average values over triplicate sampling for the following response variables: A) Fresh shoot weight, B) Fresh Root Weight, C) Dry Shoot Weight, and D) Dry Root Weight. Dashed horizontal lines indicate the average measurement observed for the respective mock-inoculated control groups. Isolate numbers along the x-axis correspond to BCW-isolate identification numbers seen in other plots and tables. Numeric annotations in some plots indicate percentage of biomass for selected isolates relative to mock-inoculated controls.

**S15 Fig. Biomass weights for *in planta* inoculation screen of mucilage isolate set 12.** Mono-isolate inoculation of potato plantlets with 22 mucilage diazotrophs were compared against a single mock-inoculated control. Each figure panel shows average values over triplicate sampling for the following response variables: A) Fresh shoot weight, B) Fresh Root Weight, C) Dry Shoot Weight, and D) Dry Root Weight. Dashed horizontal lines indicate the average measurement observed for the respective mock-inoculated control groups. Isolate numbers along the x-axis correspond to BCW-isolate identification numbers seen in other plots and tables. Numeric annotations in some plots indicate percentage of biomass for selected isolates relative to mock-inoculated controls.

**S16 Fig. Biomass weights for *in planta* inoculation screen of mucilage isolate set 13.** Mono-isolate inoculation of potato plantlets with 24 mucilage diazotrophs were compared against a single mock-inoculated control. Each figure panel shows average values over triplicate sampling for the following response variables: A) Fresh shoot weight, B) Fresh Root Weight, C) Dry Shoot Weight, and D) Dry Root Weight. Dashed horizontal lines indicate the average measurement observed for the respective mock-inoculated control groups. Isolate numbers along the x-axis correspond to BCW-isolate identification numbers seen in other plots and tables. Numeric annotations in some plots indicate percentage of biomass for selected isolates relative to mock-inoculated controls.

**S17 Fig. Potato growth responses to inoculation with mucilage isolate SynComs in the greenhouse.** Russet Burbank and Red La Soda potatoes were inoculated with 6 different SynComs and grown in the greenhouse alongside mock-inoculated control plants. Plants receiving complete N fertilization (100% Hoagland N) were assessed for: A) Shoot Weight, B) Root Weight, C) Tuber Weight and G) Number of Tubers. Plants receiving low N fertilization (20 % Hoagland N) were assessed for: D) Shoot Weight, E) Root Weight, F) Tuber Weight, and H) Number of tubers.

**S1 Table. PGP trait profiles of mucilage diazotrophic isolates.** Numbers represent counts for homologous sequences matching HMMs of marker genes for targeted PGP traits in each isolate’s whole genome sequence assembly. The number of significant matches to TIGRFAM [15] HMMs for marker genes of *nif* and non-*nif* PGP traits by predicted coding sequences identified in each isolate’s whole genome sequence were counted using functions from base and tidyverse 1.2.1 packages in R [18]. Predicted coding sequences were counted as significant homologous genes to HMMs if the threshold of a maximum e-value of 1e^-06^ and HMM coverage of 80% were met. NIF Group assignments for each isolate were annotated in the table alongside HMM search results and include Dos Santos Positive (DSP), Semi-Dos Santos (SDS) and Dos Santos Negative (DSN). Genus information for each isolate was determined using Sourmash [16] LCA classification with the GTDB v89 [17] and GenBank [47] databases with a k-size of 31. PGP trait profiles included possession of essential marker genes for the BNF trait proposed by the Dos Santos model (*nifH*, *nifD*, *nifK*, *nifE*, *nifN*, *nifB*), the marker gene for deamination of ACC (*acdS*), the marker gene for biosynthesis of IAA (*ipdC/ppdC*), and marker genes for phosphate solubilization using the PQQ mechanism mediated by the *pqqB*, *pqqC*, *pqqD*, *pqqE*, *pqqF* and PQQ Dehydrogenase (*pqq-DH*) genes. Data for homologous sequence counts of BNF marker genes were previously generated and reported [11].

**S2 Table. Summary of PGP assay values for each isolate.** Performance on each *in vitro* PGP assay was summarized for each isolate. Each record indicates the identification code assigned to the mucilage isolate (BCW-ID). Genus assignment based on classification of the isolate’s whole genome sequence generated with Sourmash 3.0.1 [16] and ^15^N/^14^N ratio values indicating performance on the ^15^N incorporation assay (BNF) were included using data from previous investigations with the isolate collection [11]. *In vitro* assays for non-*nif* PGP traits included the utilization of 1-Amino-1-cyclopropanecarboxylic acid as a nitrogen source for growth (ACC), the colorimetric assay for auxin biosynthesis (IAA), and the colorimetric assay to detect the liberation of soluble phosphate (PO4). Units for measurements of the ACC, IAA and PO4 assays were relative growth rate (RGR – see Methods), mg/mL and mg/L, respectively.

**S3 Table. Summary statistics for PGP assays by NIF group.** Data from the *in vitro* assays for biological nitrogen fixation (BNF), ACC utilization (ACC), indole-3-acetic acid biosynthesis (IAA) and phosphate solubilization (PO4) were summarized in R using base and tidyverse 1.2.1 packages [18]. Summary statistics included estimations generated with data from all isolates for mean, median, minimum (min) and maximum (max) values. NIF group corresponds to Dos Santos Positive (DSP), Dos Santos Negative (DSN), and Semi-Dos Santos (SDS).

