## supporting information for "Diazotrophic bacteria from maize exhibit multifaceted plant growth promotion traits in multiple hosts"

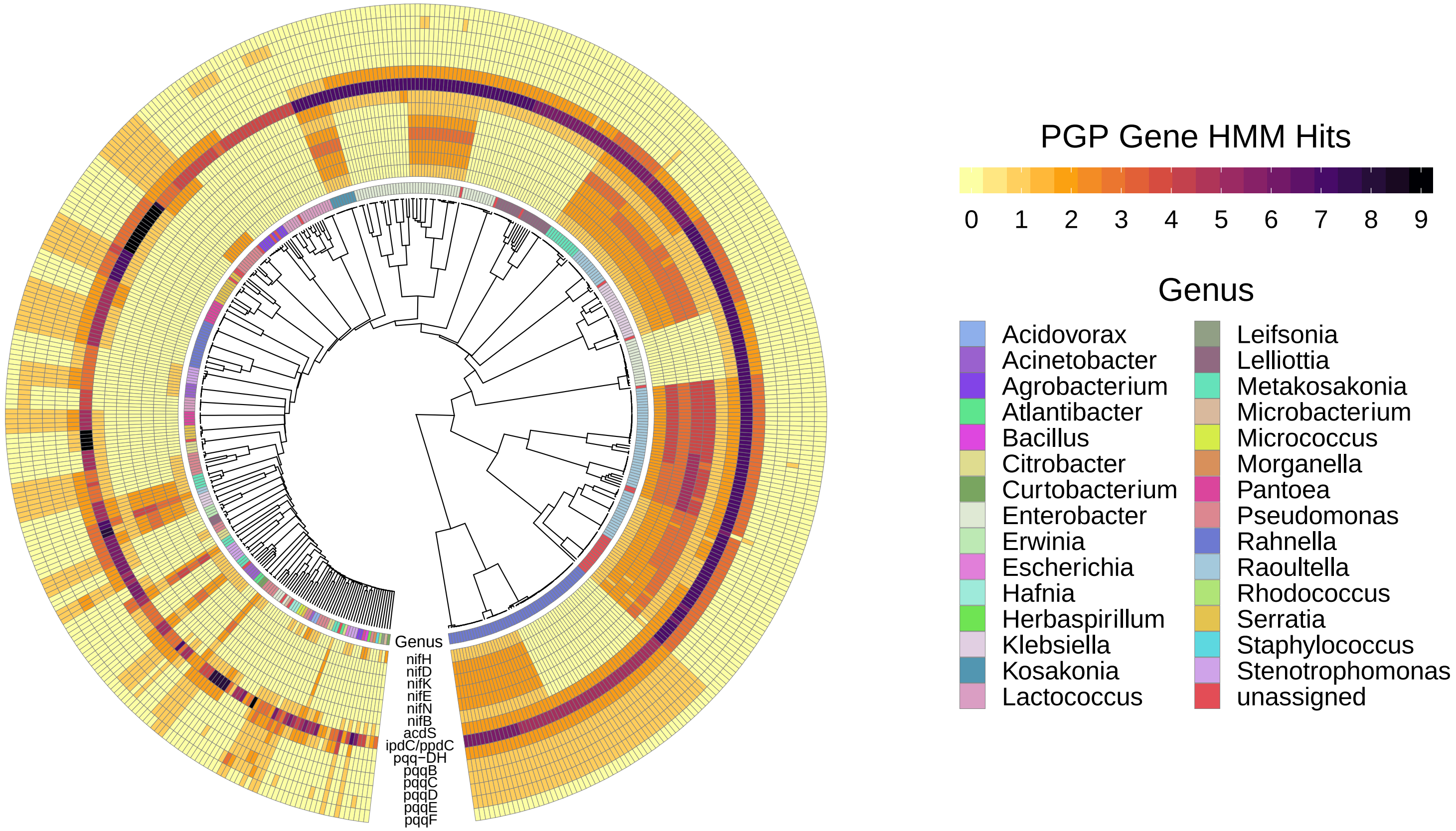

**S1 Fig. Dendrogram of diazotroph genomes with heatmaps indicating PGP gene profiles**

Marker genes for PGP functionalities were detected by scanning total amino sequences from each pure isolate genome that were predicted using Prokka against HMMs obtained from the TIGRFAM database [1]. Query sequences were considered as positive matches to the targeted HMMs if model coverage was greater than or equal to 75 % with an e-value less than or equal to 1e-9. HMMs for marker genes corresponding to the PGP traits of interest included the essential nitrogen fixation genes (*nifHDKENB*) proposed by Dos Santos et al. [10], the *acdS* gene encoding ACC deaminase, the *ipdC/ppdC* genes encoding indole pyruvate decarboxylase, individual models for the *pqqBCDEF* genes that encode corresponding subunits of the pyrroloquinoline quinone cofactor and the *pqq* associated dehydrogenase (*pqq-DH*). The PGP gene profiles for each pure isolate are presented in the context of a hierarchically clustered dendrogram corresponding to MinHash distances computed with Sourmash [2].

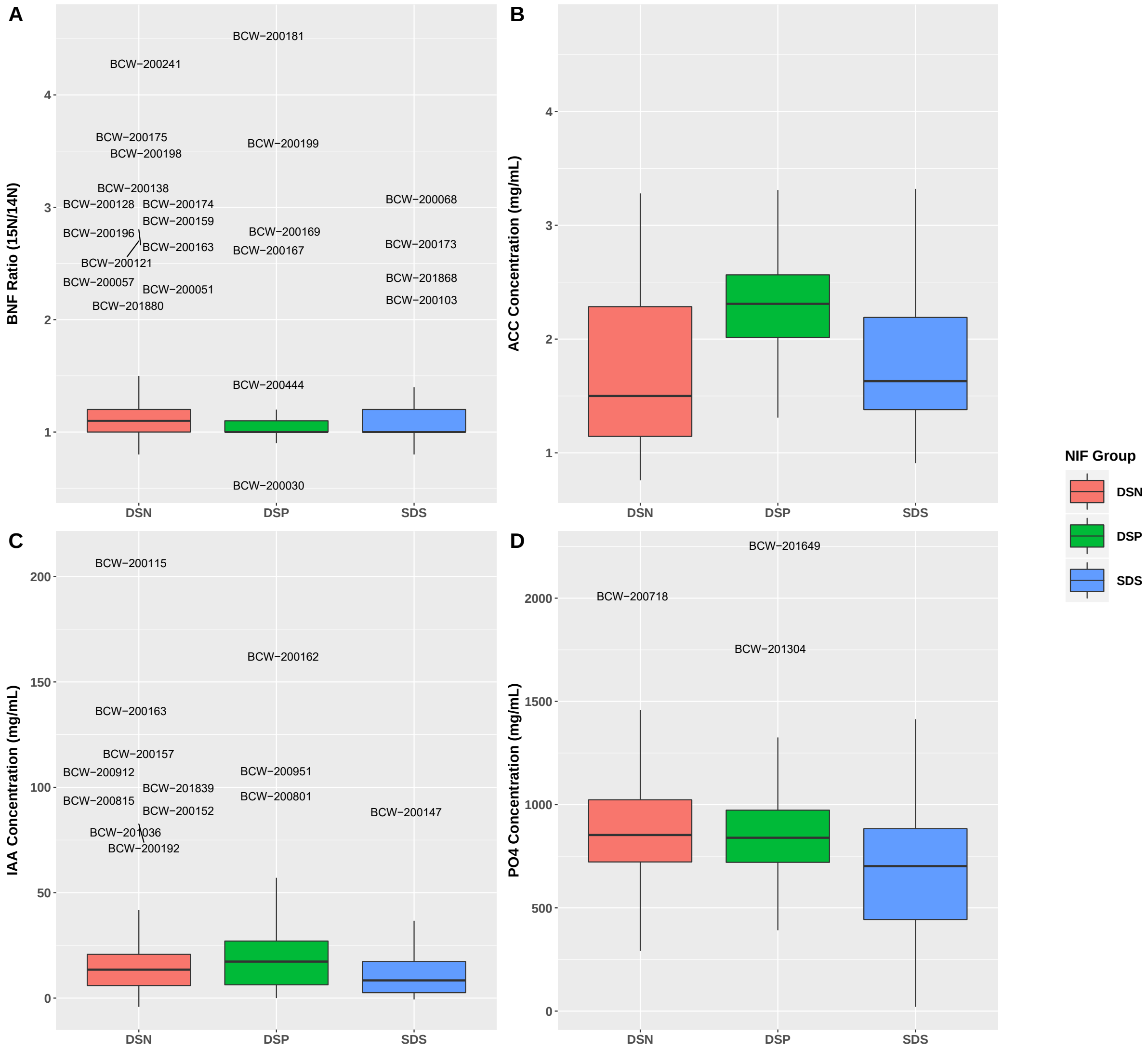


**S2 Fig. Distribution of PGP assay data from mucilage isolates across NIF groups**

Data acquired from *in vitro* assays to assess mucilage isolate phenotypes for targeted PGP traits was analyzed and plotted using R 3.5.1. Datapoints for the assays represented the average response values observed over three biological replications with each isolate. Points annotated with BCW-ID numbers indicate outliers for each assay. Outliers were determined as isolates with observed values greater than subtracting or adding 3 times the calculated interquartile range from the first or third quartile, respectively. Boxplots were made using tidyverse 1.2.1 [3], ggrepel 0.5

and cowplot 1.0.0. Code for the analysis is hosted on Github at: (https://github.com/shigdon/R-Mucilage-isolate-pgp-assay).

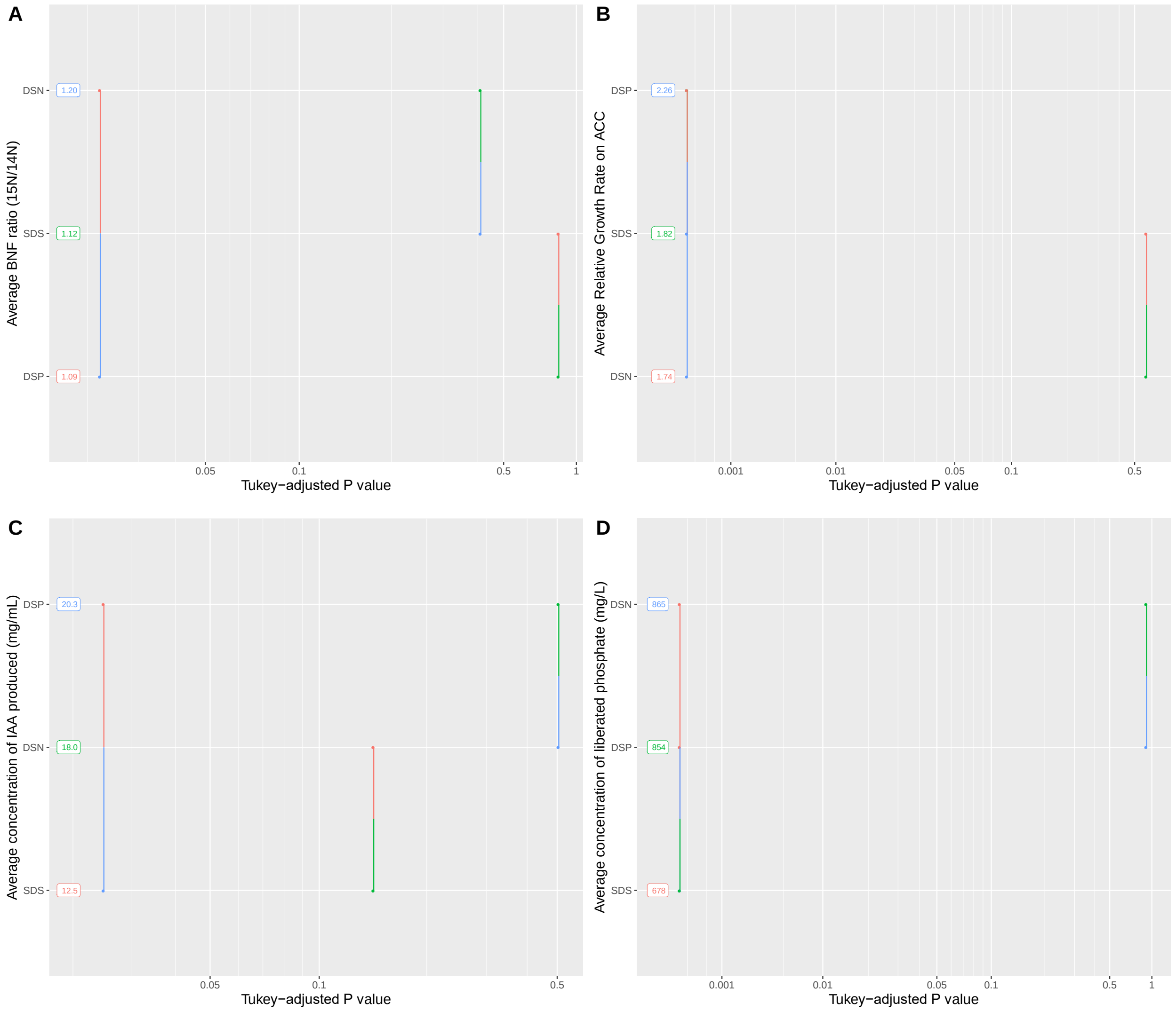


**S3 Fig. Pairwise comparisons of average PGP assay performances by NIF group**

Data obtained from *in vitro* phenotypic assays for PGP functionalities were fit to linear models to calculate mean estimates for each NIF group, compute mean difference estimates between NIF groups, and conduct multiple pairwise comparisons between the estimated means to determine statistical significance of the differences. The three NIF groups include Dos Santos Positive (DSP) isolates possessing all six essential *nif* genes, Semi-Dos Santos (SDS) isolates possessing an incomplete set of the six essential *nif* genes, and the Dos Santos Negative (DSN) isolates without any of the six essential *nif* genes. Each x-axis presents the estimated average response by the respective linear model and the y-axes depict the *p*-values of each comparison that were adjusted by the Tukey HSD method using the R package emmeans 1.4.1. A) Average BNF ratios by NIF group corresponding to the $^{15}N$ incorporation metabolomic assay; B) Average relative growth rate by NIF group for isolates cultured with 1-amino-1-cyclopropane carboxylic acid provided as the nitrogen source; C) Average production levels of indole-3-acetic acid by NIF group; D) Average liberation of soluble phosphate from hydroxyapatite by NIF group. Code for the analysis is hosted on Github at: (https://github.com/shigdon/R-Mucilage-isolate-pgp-assay).

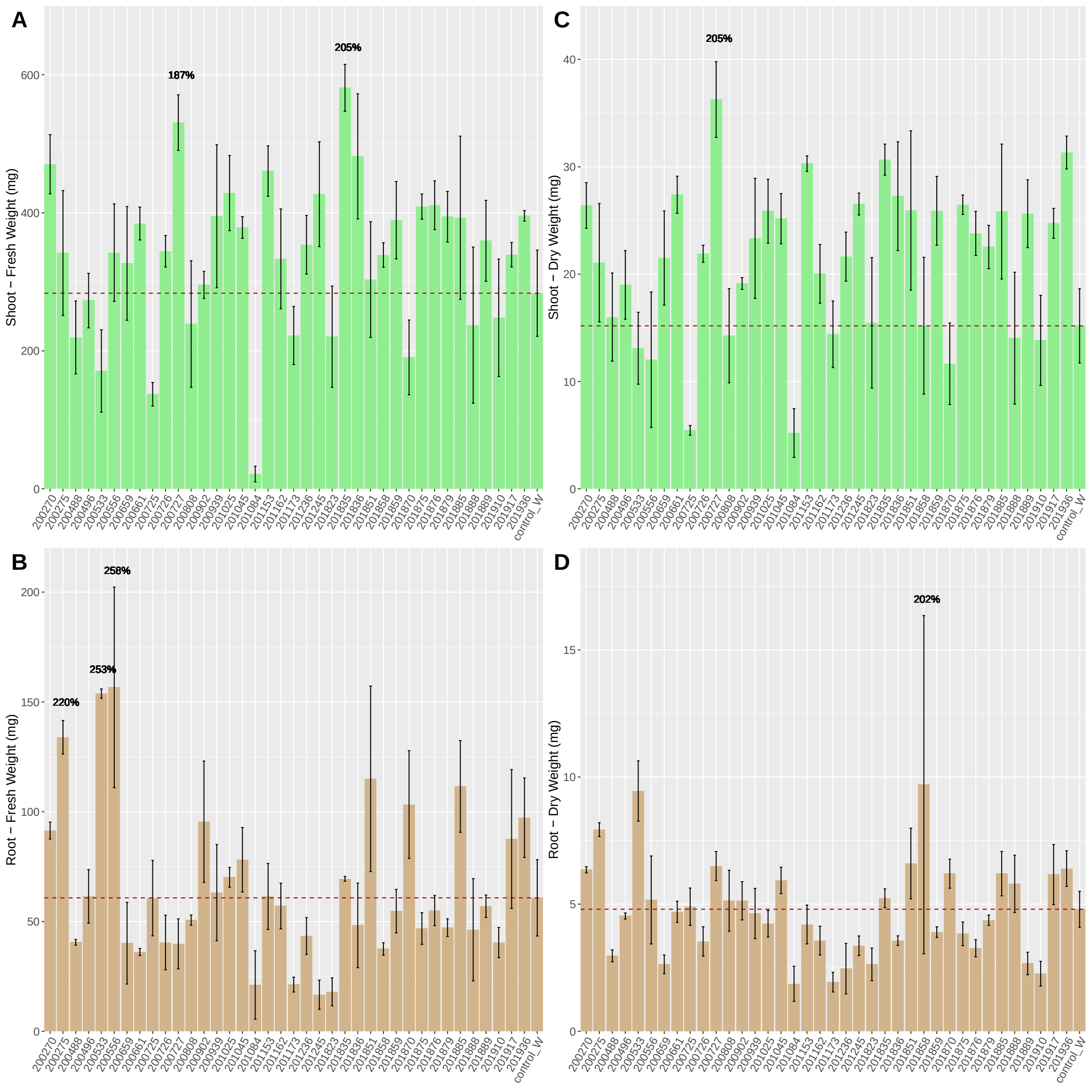


**S4 Fig. Biomass weights for *in planta* inoculation screen of mucilage isolate set 1.** Mono-isolate inoculation of potato plantlets with 38 mucilage diazotrophs were compared against a single mock-inoculated control. Each figure panel shows average values over triplicate sampling for the following response variables: A) Fresh shoot weight, B) Fresh Root Weight, C) Dry Shoot Weight, and D) Dry Root Weight. Dashed horizontal lines indicate the average measurement observed for the respective mock-inoculated control groups. Isolate numbers along the x-axis correspond to BCW-isolate identification numbers seen in other plots and tables.

Numeric annotations in some plots indicate percentage of biomass for selected isolates relative to mock-inoculated controls.

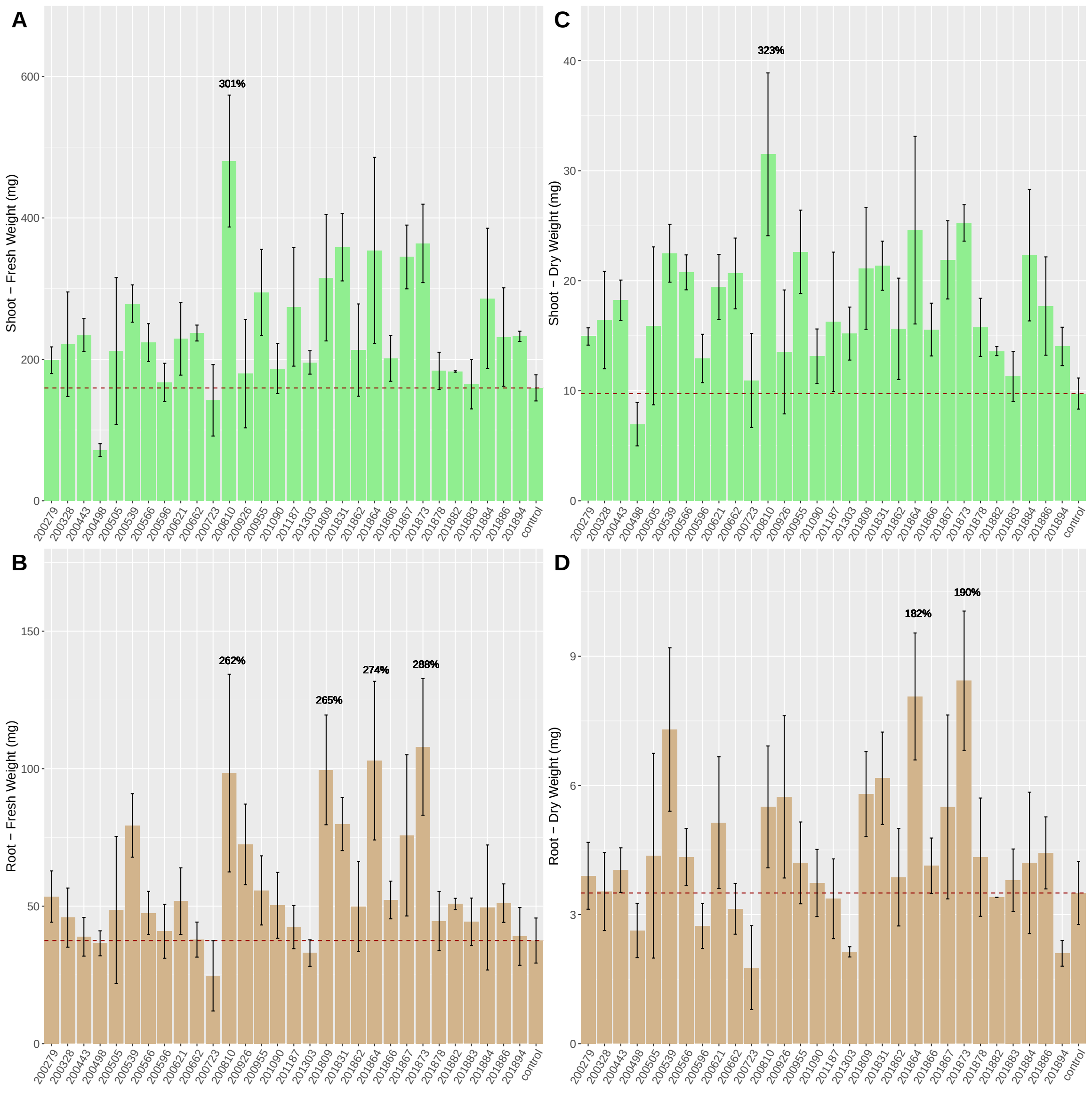


**S5 Fig. Biomass weights for *in planta* inoculation screen of mucilage isolate set 2.** Mono-isolate inoculation of potato plantlets with 30 mucilage diazotrophs were compared against a single mock-inoculated control. Each figure panel shows average values over triplicate sampling for the following response variables: A) Fresh shoot weight, B) Fresh Root Weight, C) Dry Shoot Weight, and D) Dry Root Weight. Dashed horizontal lines indicate the average measurement observed for the respective mock-inoculated control groups. Isolate numbers along the x-axis correspond to BCW-isolate identification numbers seen in other plots and tables.

Numeric annotations in some plots indicate percentage of biomass for selected isolates relative to mock-inoculated controls.

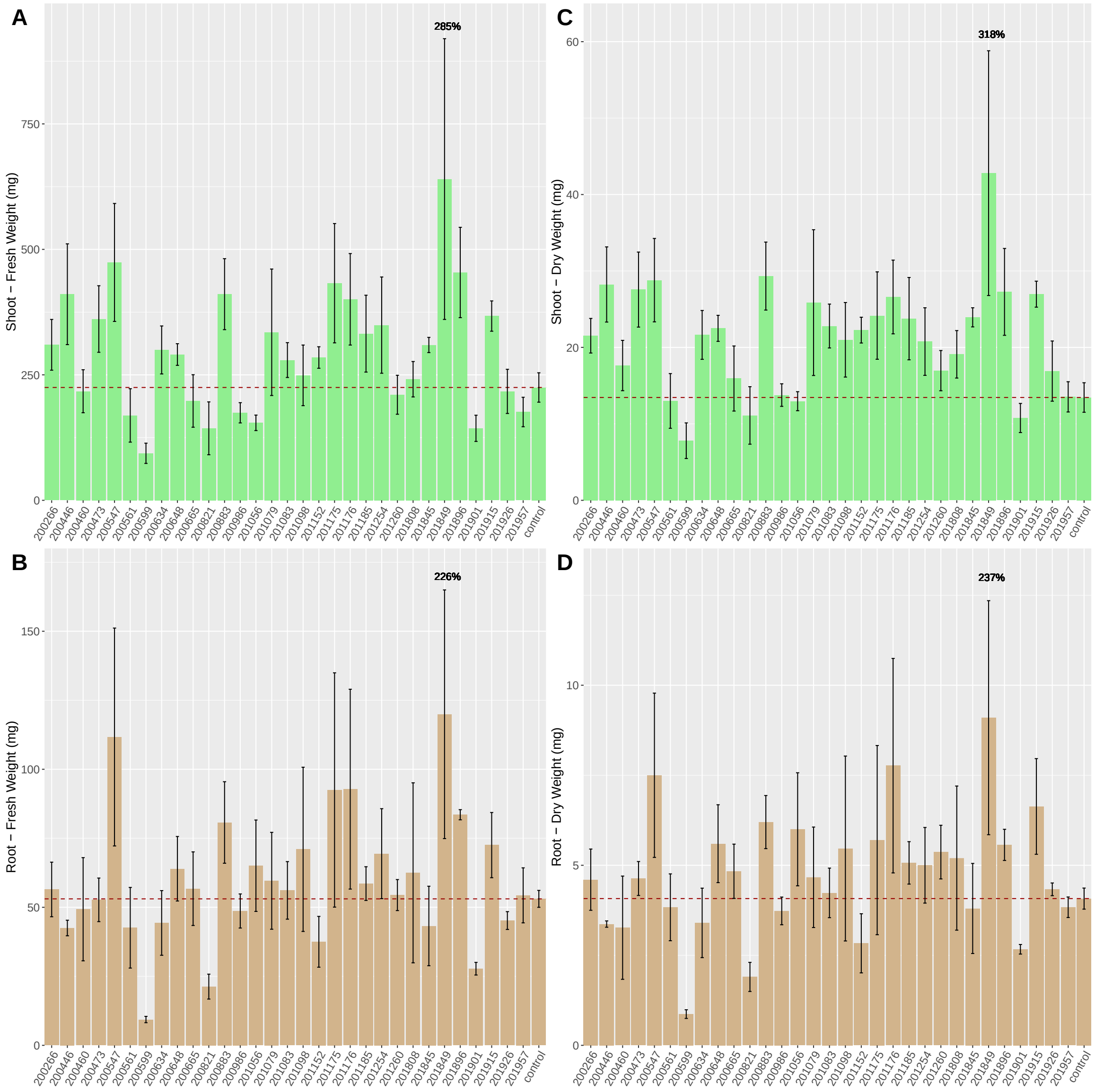


**S6 Fig. Biomass weights for *in planta* inoculation screen of mucilage isolate set 3.** Mono-isolate inoculation of potato plantlets with 31 mucilage diazotrophs were compared against a single mock-inoculated control. Each figure panel shows average values over triplicate sampling for the following response variables: A) Fresh shoot weight, B) Fresh Root Weight, C) Dry Shoot Weight, and D) Dry Root Weight. Dashed horizontal lines indicate the average measurement observed for the respective mock-inoculated control groups. Isolate numbers along the x-axis correspond to BCW-isolate identification numbers seen in other plots and tables.

Numeric annotations in some plots indicate percentage of biomass for selected isolates relative to mock-inoculated controls.

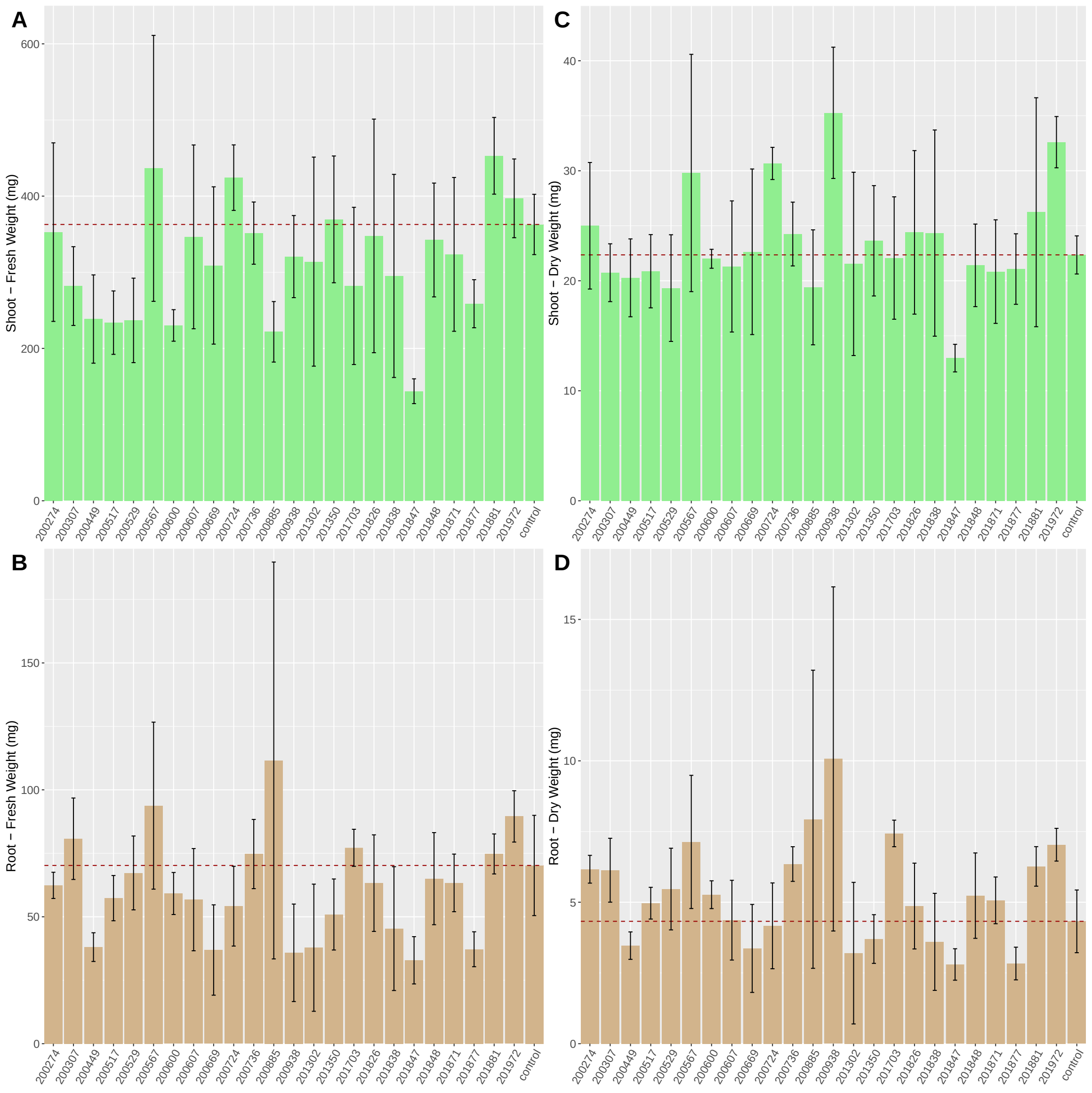


**S7 Fig. Biomass weights for *in planta* inoculation screen of mucilage isolate set 4.** Mono-isolate inoculation of potato plantlets with 24 mucilage diazotrophs were compared against a single mock-inoculated control. Each figure panel shows average values over triplicate sampling for the following response variables: A) Fresh shoot weight, B) Fresh Root Weight, C) Dry Shoot Weight, and D) Dry Root Weight. Dashed horizontal lines indicate the average measurement observed for the respective mock-inoculated control groups. Isolate numbers along the x-axis correspond to BCW-isolate identification numbers seen in other plots and tables.

Numeric annotations in some plots indicate percentage of biomass for selected isolates relative to mock-inoculated controls.

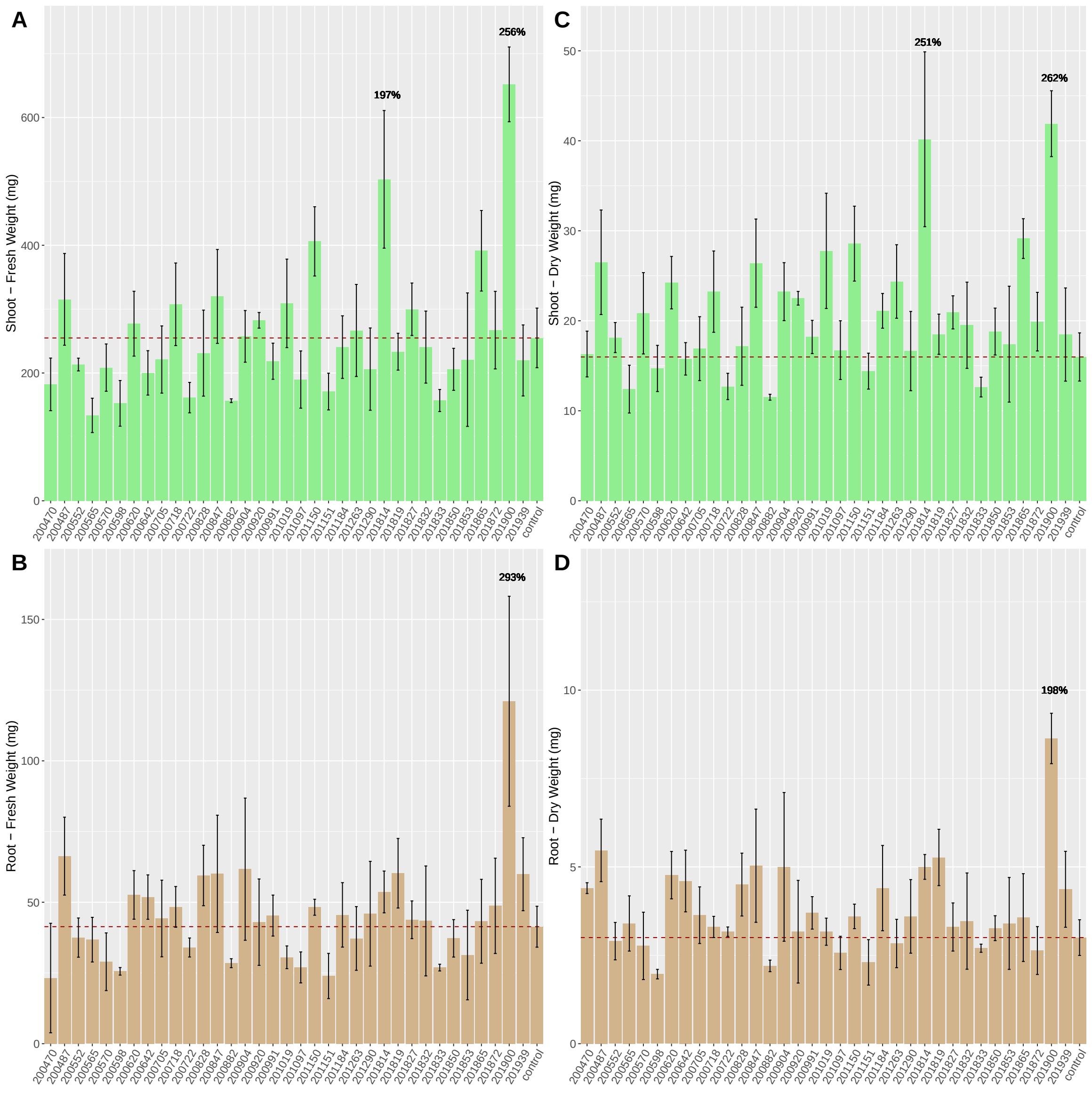


**S8 Fig. Biomass weights for *in planta* inoculation screen of mucilage isolate set 5.** Mono-isolate inoculation of potato plantlets with 35 mucilage diazotrophs were compared against a single mock-inoculated control. Each figure panel shows average values over triplicate sampling for the following response variables: A) Fresh shoot weight, B) Fresh Root Weight, C) Dry Shoot Weight, and D) Dry Root Weight. Dashed horizontal lines indicate the average measurement observed for the respective mock-inoculated control groups. Isolate numbers along the x-axis correspond to BCW-isolate identification numbers seen in other plots and tables.

Numeric annotations in some plots indicate percentage of biomass for selected isolates relative to mock-inoculated controls.

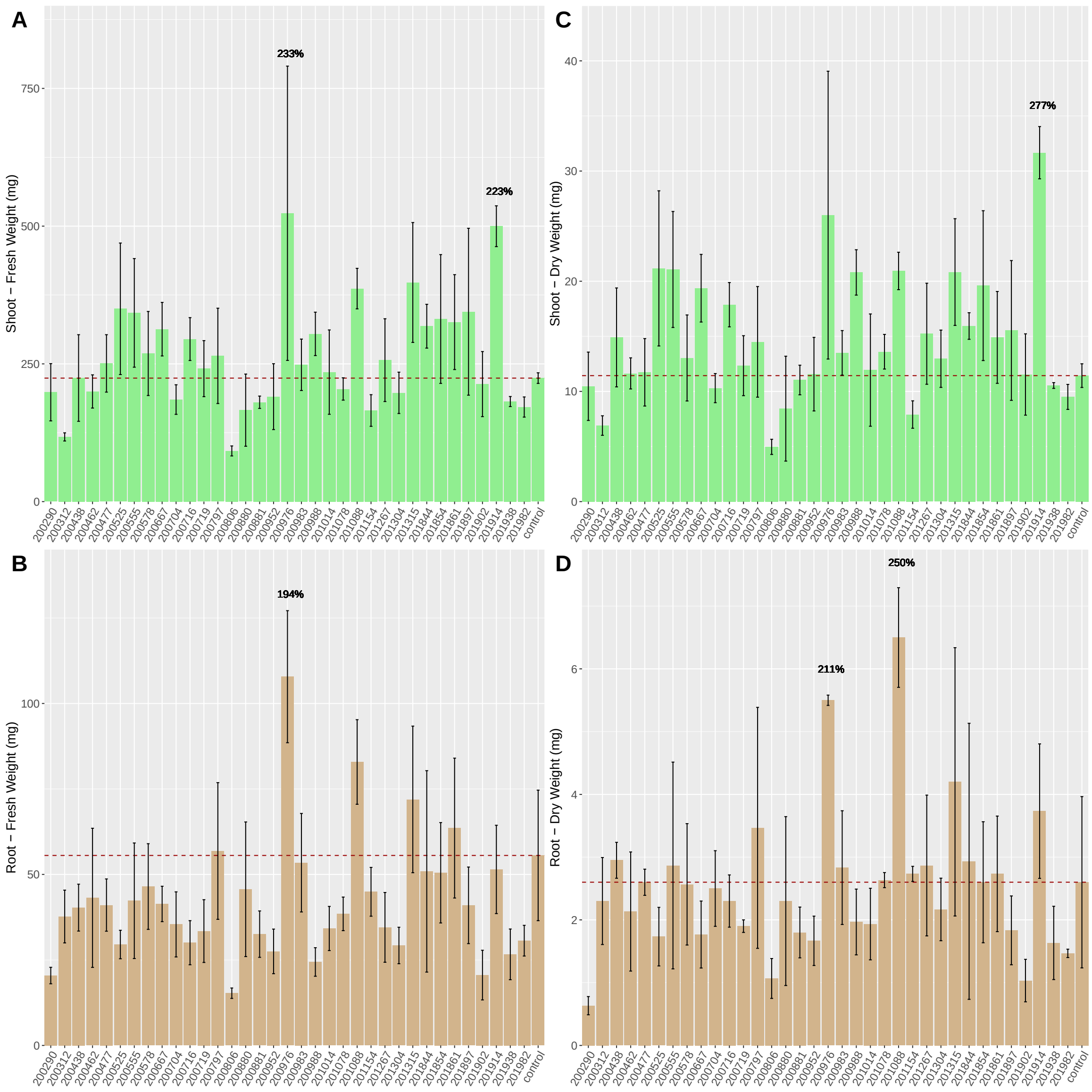


**S9 Fig. Biomass weights for *in planta* inoculation screen of mucilage isolate set 6.** Mono-isolate inoculation of potato plantlets with 35 mucilage diazotrophs were compared against a single mock-inoculated control. Each figure panel shows average values over triplicate sampling for the following response variables: A) Fresh shoot weight, B) Fresh Root Weight, C) Dry Shoot Weight, and D) Dry Root Weight. Dashed horizontal lines indicate the average measurement observed for the respective mock-inoculated control groups. Isolate numbers along the x-axis correspond to BCW-isolate identification numbers seen in other plots and tables.

Numeric annotations in some plots indicate percentage of biomass for selected isolates relative to mock-inoculated controls.

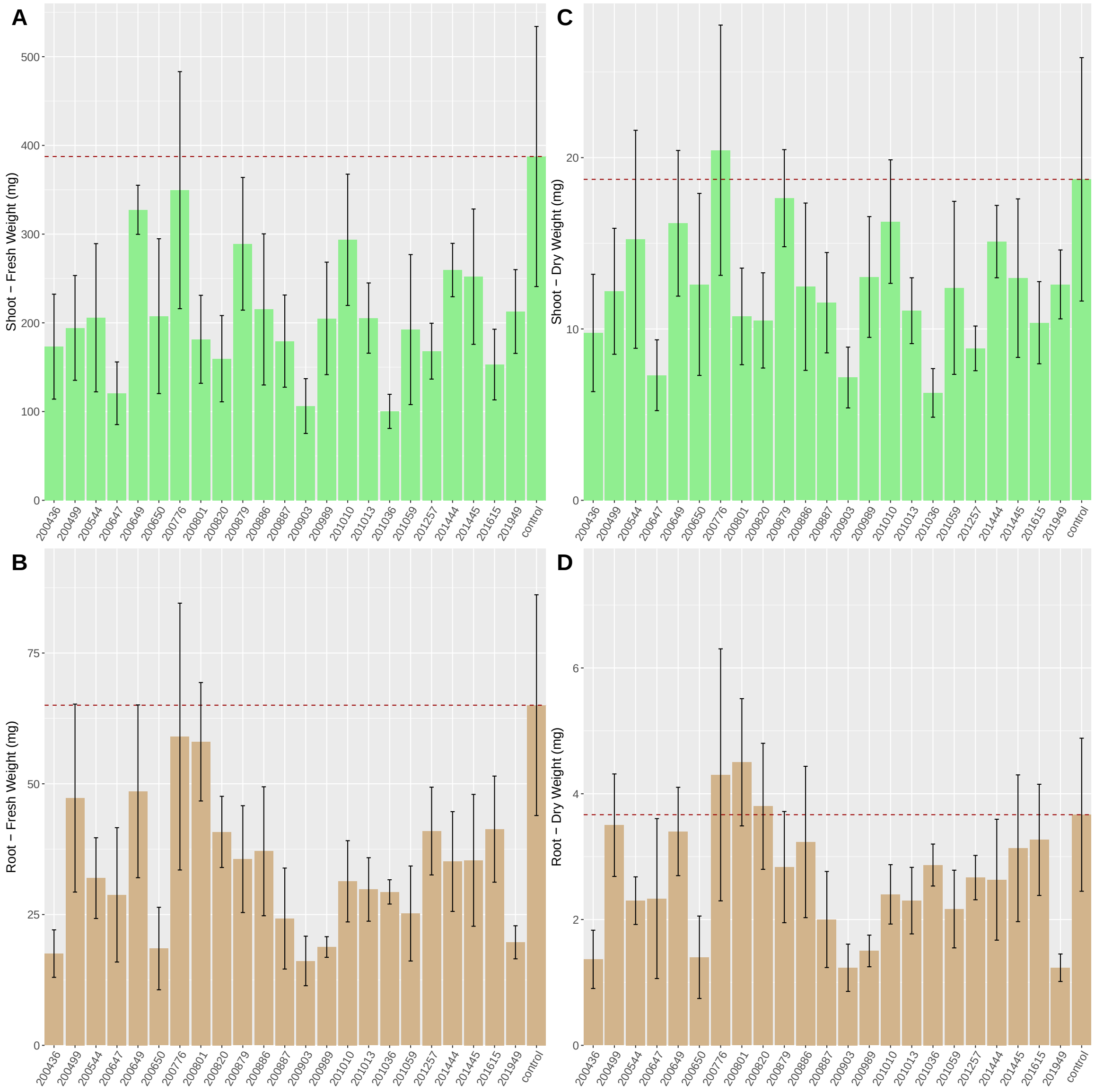


**S10 Fig. Biomass weights for *in planta* inoculation screen of mucilage isolate set 7.** Mono-isolate inoculation of potato plantlets with 23 mucilage diazotrophs were compared against a single mock-inoculated control. Each figure panel shows average values over triplicate sampling for the following response variables: A) Fresh shoot weight, B) Fresh Root Weight, C) Dry Shoot Weight, and D) Dry Root Weight. Dashed horizontal lines indicate the average measurement observed for the respective mock-inoculated control groups. Isolate numbers along the x-axis correspond to BCW-isolate identification numbers seen in other plots and tables.

Numeric annotations in some plots indicate percentage of biomass for selected isolates relative to mock-inoculated controls.

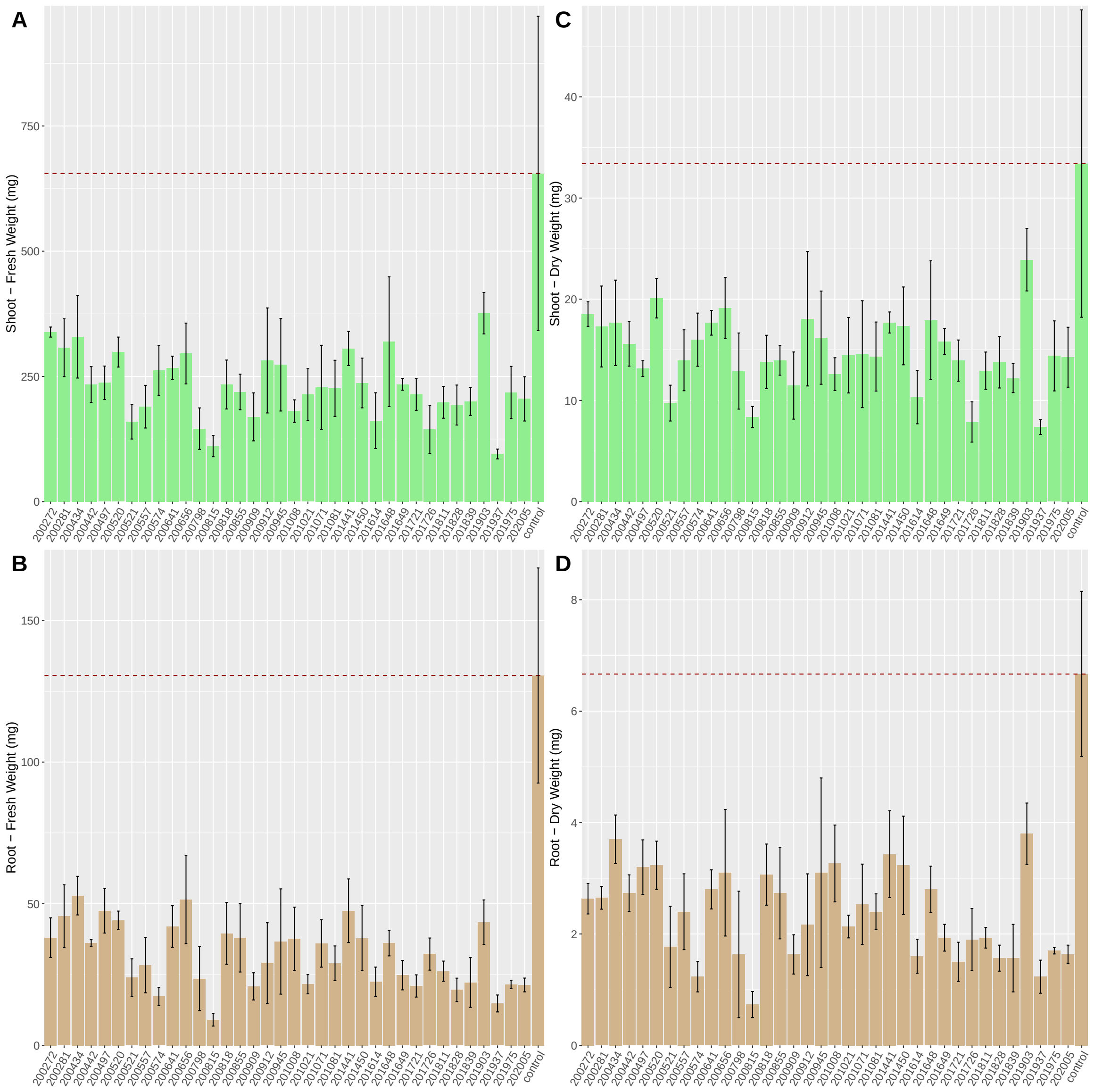


**S11 Fig. Biomass weights for *in planta* inoculation screen of mucilage isolate set 8.** Mono-isolate inoculation of potato plantlets with 36 mucilage diazotrophs were compared against a single mock-inoculated control. Each figure panel shows average values over triplicate sampling for the following response variables: A) Fresh shoot weight, B) Fresh Root Weight, C) Dry Shoot Weight, and D) Dry Root Weight. Dashed horizontal lines indicate the average measurement observed for the respective mock-inoculated control groups. Isolate numbers along the x-axis correspond to BCW-isolate identification numbers seen in other plots and tables.

Numeric annotations in some plots indicate percentage of biomass for selected isolates relative to mock-inoculated controls.

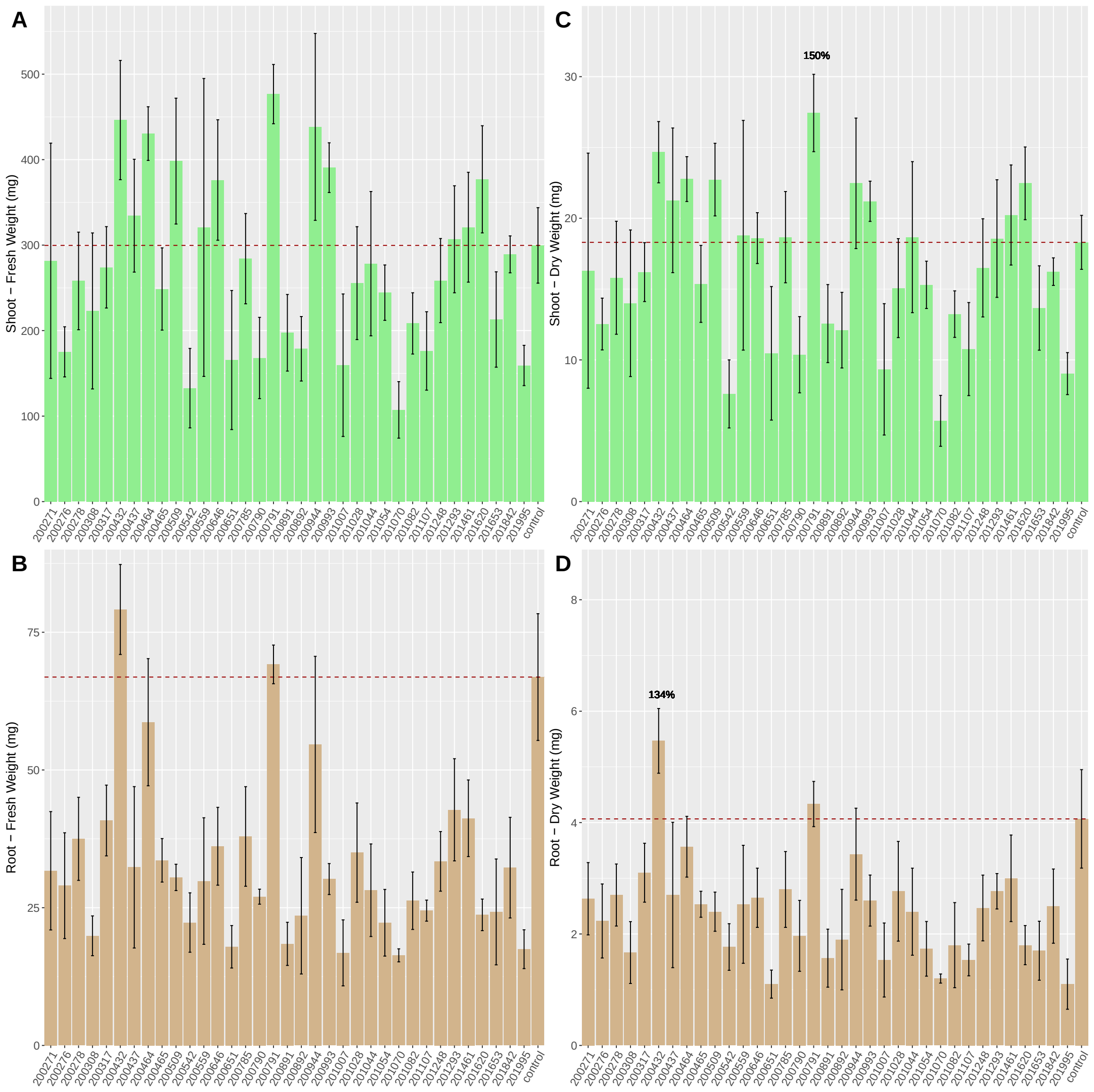


**S12 Fig. Biomass weights for *in planta* inoculation screen of mucilage isolate set 9.** Mono-isolate inoculation of potato plantlets with 35 mucilage diazotrophs were compared against a single mock-inoculated control. Each figure panel shows average values over triplicate sampling for the following response variables: A) Fresh shoot weight, B) Fresh Root Weight, C) Dry Shoot Weight, and D) Dry Root Weight. Dashed horizontal lines indicate the average measurement observed for the respective mock-inoculated control groups. Isolate numbers along the x-axis correspond to BCW-isolate identification numbers seen in other plots and tables.

Numeric annotations in some plots indicate percentage of biomass for selected isolates relative to mock-inoculated controls.

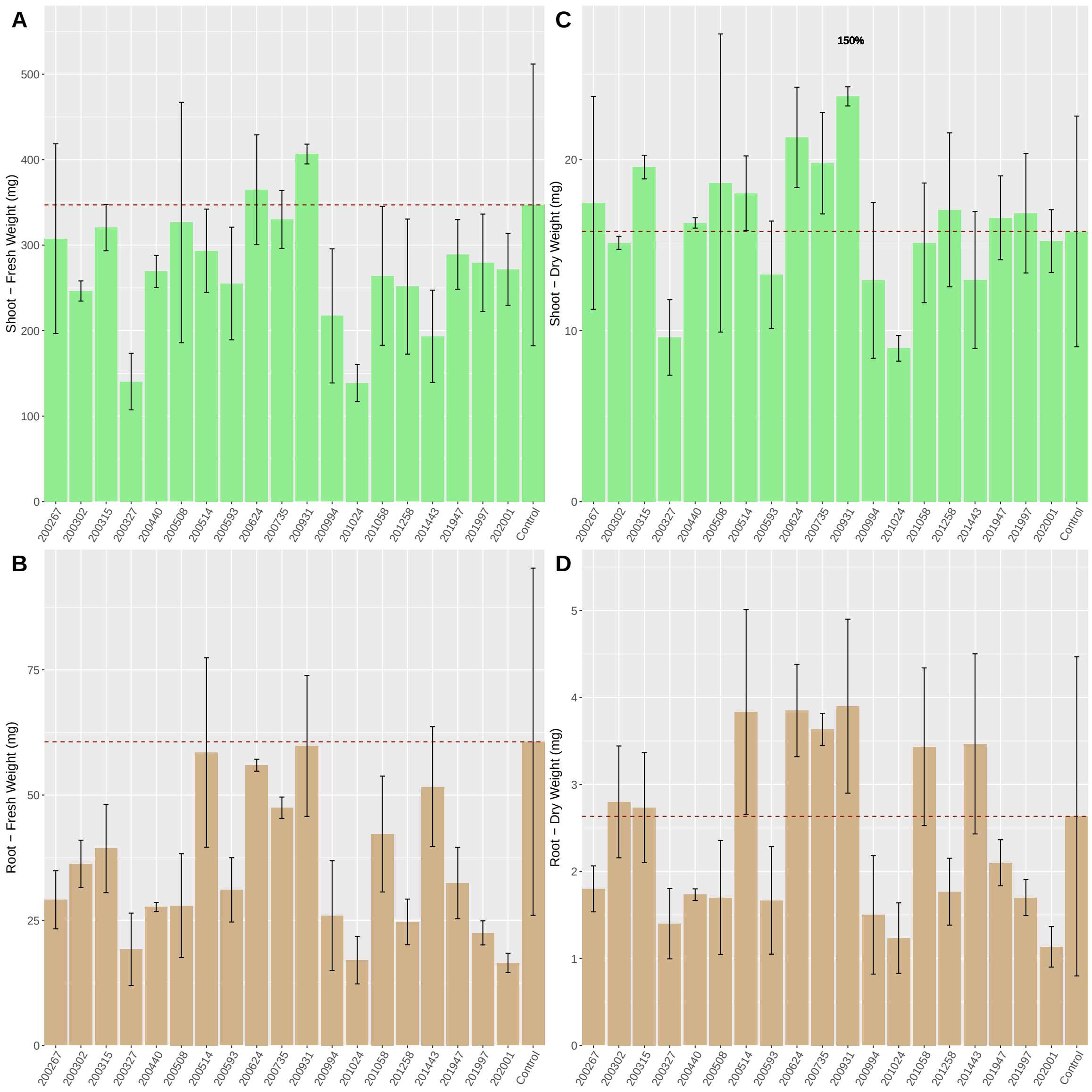


**S13 Fig. Biomass weights for *in planta* inoculation screen of mucilage isolate set 10.** Mono-isolate inoculation of potato plantlets with 19 mucilage diazotrophs were compared against a single mock-inoculated control. Each figure panel shows average values over triplicate sampling for the following response variables: A) Fresh shoot weight, B) Fresh Root Weight, C) Dry Shoot Weight, and D) Dry Root Weight. Dashed horizontal lines indicate the average measurement observed for the respective mock-inoculated control groups. Isolate numbers along the x-axis correspond to BCW-isolate identification numbers seen in other plots and tables.

Numeric annotations in some plots indicate percentage of biomass for selected isolates relative to mock-inoculated controls.

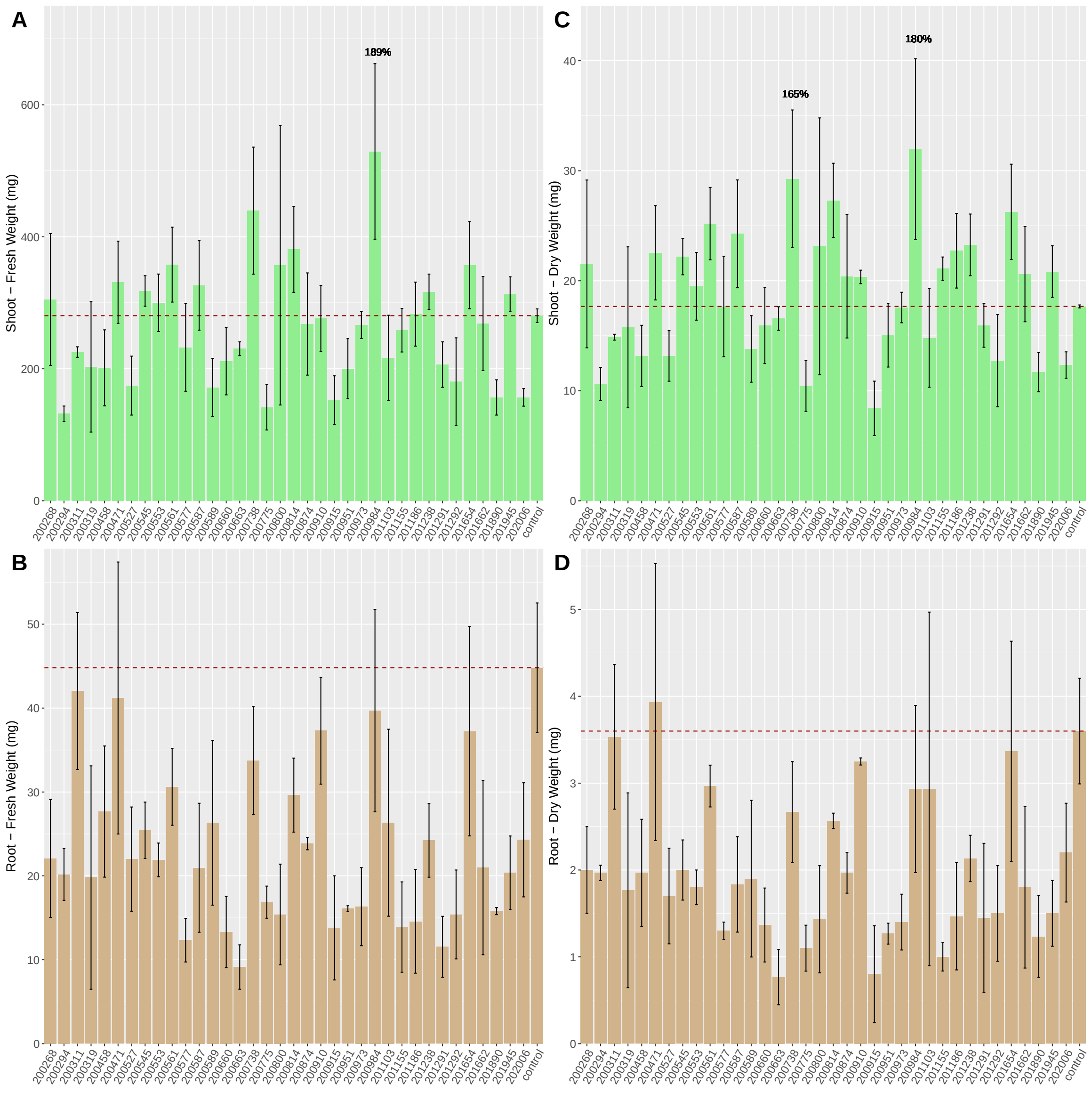


**S14 Fig. Biomass weights for *in planta* inoculation screen of mucilage isolate set 11.** Mono-isolate inoculation of potato plantlets with 36 mucilage diazotrophs were compared against a single mock-inoculated control. Each figure panel shows average values over triplicate sampling for the following response variables: A) Fresh shoot weight, B) Fresh Root Weight, C) Dry Shoot Weight, and D) Dry Root Weight. Dashed horizontal lines indicate the average measurement observed for the respective mock-inoculated control groups. Isolate numbers along the x-axis correspond to BCW-isolate identification numbers seen in other plots and tables.

Numeric annotations in some plots indicate percentage of biomass for selected isolates relative to mock-inoculated controls.

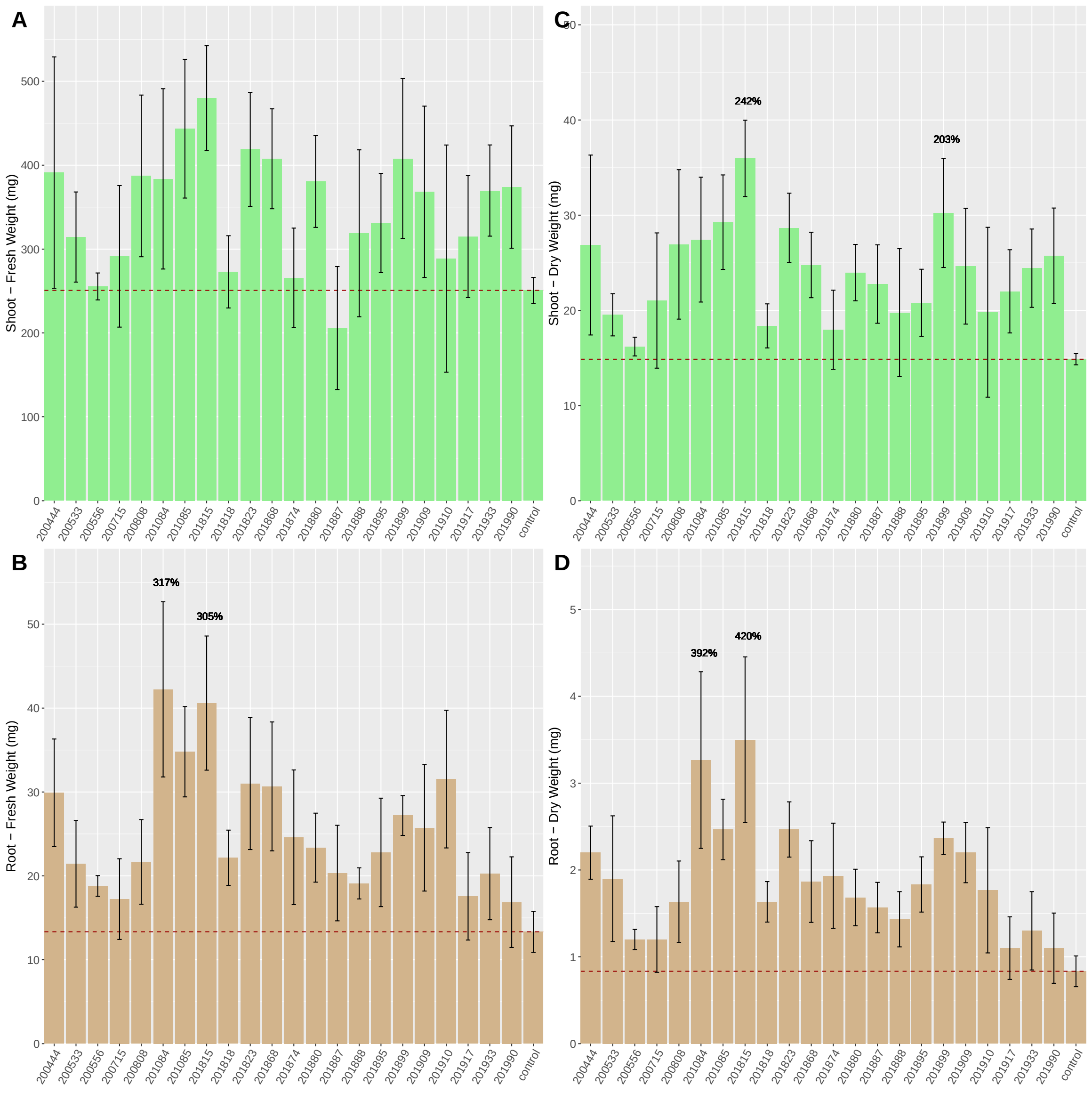


**S15 Fig. Biomass weights for *in planta* inoculation screen of mucilage isolate set 12.** Mono-isolate inoculation of potato plantlets with 22 mucilage diazotrophs were compared against a single mock-inoculated control. Each figure panel shows average values over triplicate sampling for the following response variables: A) Fresh shoot weight, B) Fresh Root Weight, C) Dry Shoot Weight, and D) Dry Root Weight. Dashed horizontal lines indicate the average measurement observed for the respective mock-inoculated control groups. Isolate numbers along the x-axis correspond to BCW-isolate identification numbers seen in other plots and tables.

Numeric annotations in some plots indicate percentage of biomass for selected isolates relative to mock-inoculated controls.

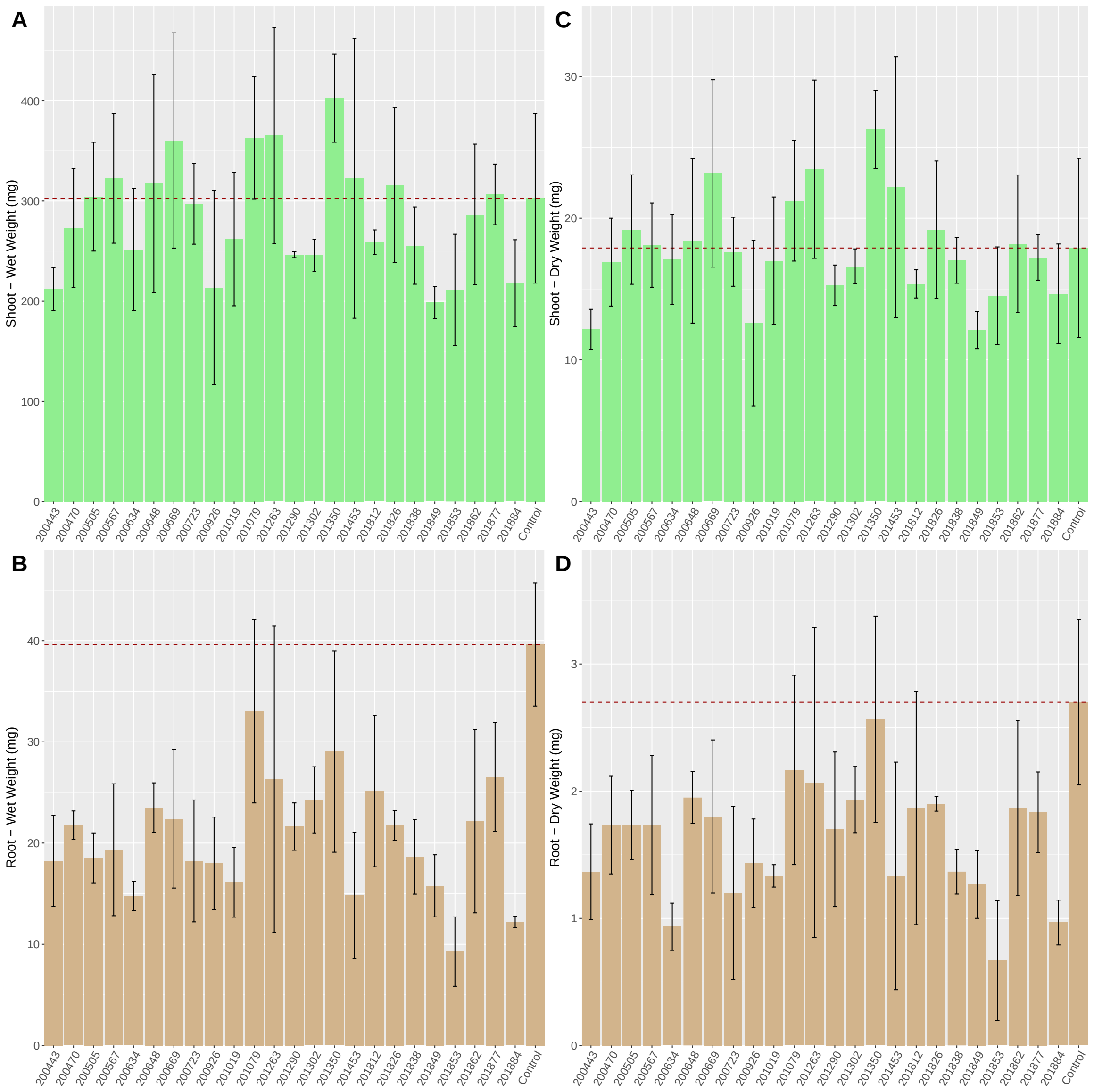


**S16 Fig. Biomass weights for *in planta* inoculation screen of mucilage isolate set 13.** Mono-isolate inoculation of potato plantlets with 24 mucilage diazotrophs were compared against a single mock-inoculated control. Each figure panel shows average values over triplicate sampling for the following response variables: A) Fresh shoot weight, B) Fresh Root Weight, C) Dry Shoot Weight, and D) Dry Root Weight. Dashed horizontal lines indicate the average measurement observed for the respective mock-inoculated control groups. Isolate numbers along the x-axis correspond to BCW-isolate identification numbers seen in other plots and tables.

Numeric annotations in some plots indicate percentage of biomass for selected isolates relative to mock-inoculated controls.

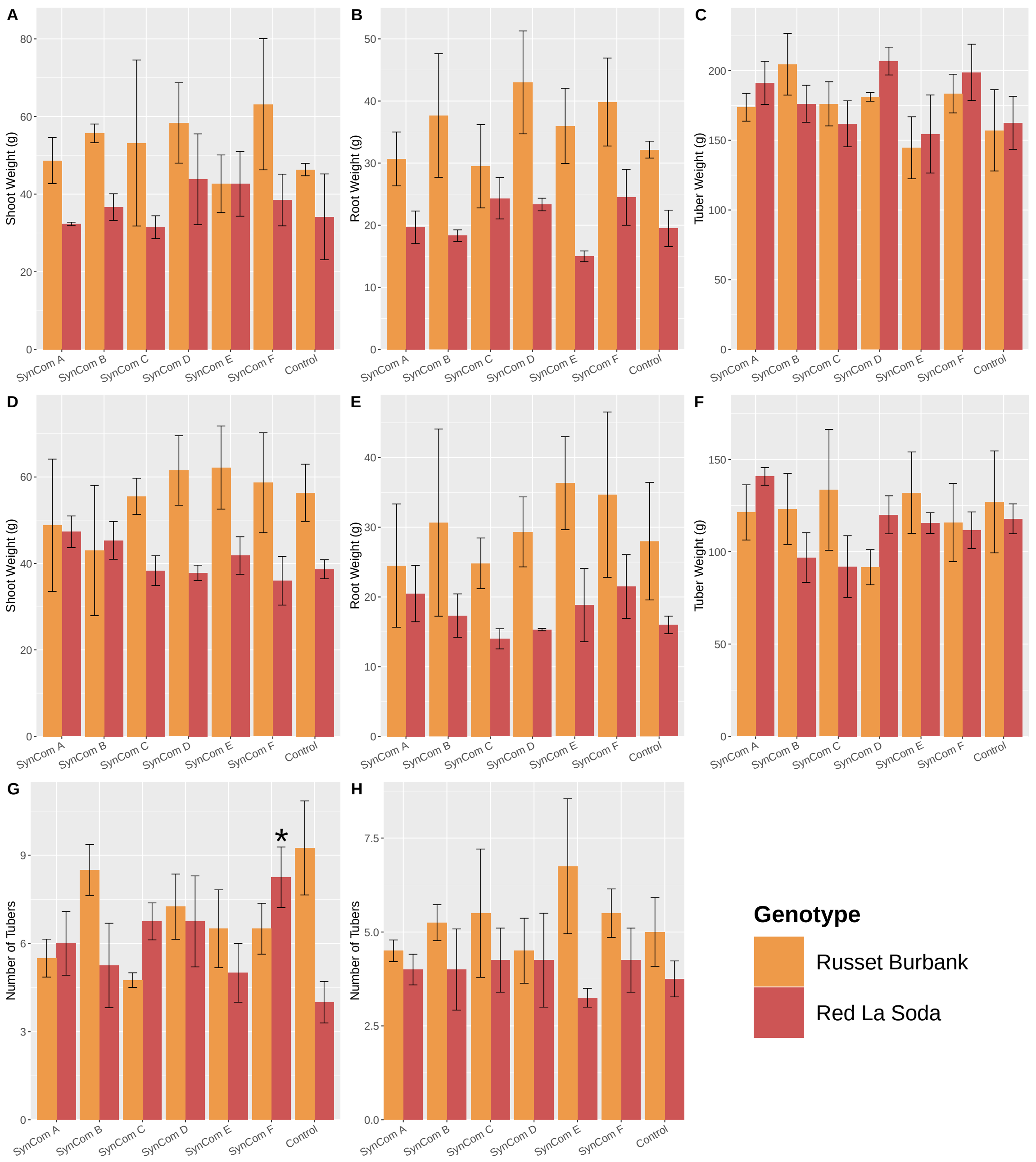


**S17 Fig. Potato growth responses to inoculation with mucilage isolate SynComs in the greenhouse.** Russet Burbank and Red La Soda potatoes were inoculated with 6 different SynComs and grown in the greenhouse alongside mock-inoculated control plants. Plants receiving complete N fertilization (100% Hoagland N) were assessed for: A) Shoot Weight, B) Root Weight, C) Tuber Weight and G) Number of Tubers. Plants receiving low N fertilization

(20 % Hoagland N) were assessed for: D) Shoot Weight, E) Root Weight, F) Tuber Weight, and H) Number of tubers.

### S1 Table. PGP trait profiles of mucilage diazotrophic isolates

| Isolate ID | Genus | NIF Grp | *nifH* | *nifD* | *nifK* | *nifE* | *nifN* | *nifB* | *acdS* | *ipdC/ ppdC* | *pqqB* | *pqqC* | *pqqD* | *pqqE* | *pqqF* | *pqq-DH* |
|---|---|---|---|---|---|---|---|---|---|---|---|---|---|---|---|---|
| BCW-200003 | *Lelliottia* | DSN | 0 | 0 | 0 | 0 | 0 | 0 | 1 | 6 | 0 | 0 | 0 | 0 | 0 | 2 |
| BCW-200009 | *Enterobacter* | DSN | 0 | 0 | 0 | 0 | 0 | 0 | 1 | 7 | 0 | 0 | 0 | 0 | 0 | 2 |
| BCW-200012 | *Enterobacter* | DSN | 0 | 0 | 0 | 0 | 0 | 0 | 1 | 7 | 0 | 0 | 0 | 0 | 0 | 2 |
| BCW-200013 | *Enterobacter* | DSN | 0 | 0 | 0 | 0 | 0 | 0 | 1 | 7 | 0 | 0 | 0 | 0 | 0 | 2 |
| BCW-200015 | *Enterobacter* | DSN | 0 | 0 | 0 | 0 | 0 | 0 | 1 | 7 | 0 | 0 | 0 | 0 | 0 | 2 |
| BCW-200018 | *Enterobacter* | DSN | 0 | 0 | 0 | 0 | 0 | 0 | 1 | 7 | 0 | 0 | 0 | 0 | 0 | 2 |
| BCW-200023 | *Enterobacter* | DSN | 0 | 0 | 0 | 0 | 0 | 0 | 1 | 7 | 0 | 0 | 0 | 0 | 0 | 2 |
| BCW-200025 | *Enterobacter* | DSN | 0 | 0 | 0 | 0 | 0 | 0 | 1 | 7 | 0 | 0 | 0 | 0 | 0 | 2 |
| BCW-200027 | *Enterobacter* | DSN | 0 | 0 | 0 | 0 | 0 | 0 | 1 | 7 | 0 | 0 | 0 | 0 | 0 | 2 |
| BCW-200029 | *Enterobacter* | DSN | 0 | 0 | 0 | 0 | 0 | 0 | 1 | 7 | 0 | 0 | 0 | 0 | 0 | 2 |
| BCW-200031 | *Citrobacter* | DSN | 0 | 0 | 0 | 0 | 0 | 0 | 1 | 6 | 0 | 0 | 0 | 0 | 0 | 1 |
| BCW-200033 | *Lelliottia* | DSN | 0 | 0 | 0 | 0 | 0 | 0 | 1 | 6 | 0 | 0 | 0 | 0 | 0 | 2 |
| BCW-200036 | *Citrobacter* | DSN | 0 | 0 | 0 | 0 | 0 | 0 | 1 | 5 | 0 | 0 | 0 | 0 | 0 | 1 |
| BCW-200040 | *Lelliottia* | DSN | 0 | 0 | 0 | 0 | 0 | 0 | 1 | 6 | 0 | 0 | 0 | 0 | 0 | 2 |
| BCW-200041 | *Enterobacter* | DSN | 0 | 0 | 0 | 0 | 0 | 0 | 1 | 7 | 0 | 0 | 0 | 0 | 0 | 2 |
| BCW-200043 | *Enterobacter* | DSN | 0 | 0 | 0 | 0 | 0 | 0 | 1 | 8 | 0 | 0 | 0 | 0 | 0 | 2 |
| BCW-200047 | *Enterobacter* | DSN | 0 | 0 | 0 | 0 | 0 | 0 | 1 | 7 | 0 | 0 | 0 | 0 | 0 | 2 |
| BCW-200051 | *Lactococcus* | DSN | 0 | 0 | 0 | 0 | 0 | 0 | 0 | 4 | 0 | 0 | 0 | 0 | 0 | 0 |
| BCW-200054 | *Enterobacter* | DSN | 0 | 0 | 0 | 0 | 0 | 0 | 1 | 7 | 0 | 0 | 0 | 0 | 0 | 2 |
| BCW-200055 | *Enterobacter* | DSN | 0 | 0 | 0 | 0 | 0 | 0 | 1 | 7 | 0 | 0 | 0 | 0 | 0 | 2 |
| BCW-200057 | *Microbacterium* | DSN | 0 | 0 | 0 | 0 | 0 | 0 | 1 | 3 | 0 | 0 | 0 | 0 | 0 | 0 |
| BCW-200060 | *Metakosakonia* | DSN | 0 | 0 | 0 | 0 | 0 | 0 | 2 | 4 | 0 | 0 | 0 | 1 | 0 | 2 |
| BCW-200061 | *Serratia* | DSN | 0 | 0 | 0 | 0 | 0 | 0 | 1 | 9 | 0 | 0 | 0 | 1 | 0 | 1 |
| BCW-200063 | *unassigned* | DSN | 0 | 0 | 0 | 0 | 0 | 0 | 1 | 1 | 0 | 0 | 1 | 0 | 0 | 0 |
| BCW-200064 | *Enterobacter* | DSN | 0 | 0 | 0 | 0 | 0 | 0 | 1 | 7 | 0 | 0 | 0 | 0 | 0 | 2 |

| Isolate ID | Genus | NIF Grp | *nifH* | *nifD* | *nifK* | *nifE* | *nifN* | *nifB* | *acdS* | *ipdC/ ppdC* | *pqqB* | *pqqC* | *pqqD* | *pqqE* | *pqqF* | *pqq-DH* |
|---|---|---|---|---|---|---|---|---|---|---|---|---|---|---|---|---|
| BCW-200066 | *Atlantibacter* | DSN | 0 | 0 | 0 | 0 | 0 | 0 | 1 | 5 | 0 | 0 | 0 | 0 | 0 | 2 |
| BCW-200067 | *Microbacterium* | DSN | 0 | 0 | 0 | 0 | 0 | 0 | 1 | 2 | 0 | 0 | 0 | 0 | 0 | 0 |
| BCW-200071 | *Lelliottia* | DSN | 0 | 0 | 0 | 0 | 0 | 0 | 1 | 6 | 0 | 0 | 0 | 0 | 0 | 1 |
| BCW-200077 | *Lactococcus* | DSN | 0 | 0 | 0 | 0 | 0 | 0 | 0 | 4 | 0 | 0 | 0 | 1 | 0 | 0 |
| BCW-200079 | *unassigned* | DSN | 0 | 0 | 0 | 0 | 0 | 0 | 0 | 4 | 0 | 0 | 0 | 1 | 0 | 0 |
| BCW-200082 | *Enterobacter* | DSN | 0 | 0 | 0 | 0 | 0 | 0 | 1 | 8 | 0 | 0 | 0 | 0 | 0 | 2 |
| BCW-200092 | *Enterobacter* | DSN | 0 | 0 | 0 | 0 | 0 | 0 | 1 | 7 | 0 | 0 | 0 | 0 | 0 | 2 |
| BCW-200094 | *unassigned* | DSN | 0 | 0 | 0 | 0 | 0 | 0 | 2 | 8 | 0 | 0 | 0 | 0 | 0 | 2 |
| BCW-200097 | *Acinetobacter* | DSN | 0 | 0 | 0 | 0 | 0 | 0 | 1 | 3 | 1 | 1 | 1 | 1 | 0 | 2 |
| BCW-200104 | *Enterobacter* | DSN | 0 | 0 | 0 | 0 | 0 | 0 | 1 | 8 | 0 | 0 | 0 | 0 | 0 | 2 |
| BCW-200107 | *Enterobacter* | DSN | 0 | 0 | 0 | 0 | 0 | 0 | 2 | 7 | 0 | 0 | 0 | 0 | 0 | 2 |
| BCW-200111 | *Citrobacter* | DSN | 0 | 0 | 0 | 0 | 0 | 0 | 2 | 5 | 0 | 0 | 0 | 0 | 0 | 1 |
| BCW-200114 | *Serratia* | DSN | 0 | 0 | 0 | 0 | 0 | 0 | 1 | 9 | 0 | 0 | 0 | 0 | 0 | 3 |
| BCW-200115 | *Morganella* | DSN | 0 | 0 | 0 | 0 | 0 | 0 | 1 | 6 | 0 | 0 | 0 | 0 | 0 | 0 |
| BCW-200121 | *Lactococcus* | DSN | 0 | 0 | 0 | 0 | 0 | 0 | 0 | 4 | 0 | 0 | 0 | 1 | 0 | 0 |
| BCW-200128 | *Lactococcus* | DSN | 0 | 0 | 0 | 0 | 0 | 0 | 0 | 4 | 0 | 0 | 0 | 1 | 0 | 0 |
| BCW-200138 | *Lactococcus* | DSN | 0 | 0 | 0 | 0 | 0 | 0 | 0 | 4 | 0 | 0 | 0 | 1 | 0 | 0 |
| BCW-200143 | *Rahnella* | DSN | 0 | 0 | 0 | 0 | 0 | 0 | 2 | 5 | 0 | 0 | 0 | 0 | 0 | 2 |
| BCW-200144 | *Rahnella* | DSN | 0 | 0 | 0 | 0 | 0 | 0 | 2 | 5 | 0 | 0 | 0 | 0 | 0 | 2 |
| BCW-200145 | *Rahnella* | DSN | 0 | 0 | 0 | 0 | 0 | 0 | 1 | 5 | 1 | 1 | 1 | 1 | 1 | 2 |
| BCW-200146 | *Rahnella* | DSN | 0 | 0 | 0 | 0 | 0 | 0 | 2 | 5 | 0 | 0 | 0 | 0 | 0 | 2 |
| BCW-200149 | *Rahnella* | DSN | 0 | 0 | 0 | 0 | 0 | 0 | 2 | 5 | 0 | 0 | 0 | 0 | 0 | 2 |
| BCW-200150 | *Lactococcus* | DSN | 0 | 0 | 0 | 0 | 0 | 0 | 0 | 4 | 0 | 0 | 0 | 1 | 0 | 0 |
| BCW-200151 | *Rahnella* | DSN | 0 | 0 | 0 | 0 | 0 | 0 | 1 | 5 | 1 | 1 | 1 | 1 | 1 | 2 |
| BCW-200152 | *Rahnella* | DSN | 0 | 0 | 0 | 0 | 0 | 0 | 2 | 5 | 0 | 0 | 0 | 0 | 0 | 2 |
| BCW-200155 | *Rahnella* | DSN | 0 | 0 | 0 | 0 | 0 | 0 | 1 | 5 | 1 | 1 | 1 | 1 | 1 | 2 |

| Isolate ID | Genus | NIF Grp | *nifH* | *nifD* | *nifK* | *nifE* | *nifN* | *nifB* | *acdS* | *ipdC/ ppdC* | *pqqB* | *pqqC* | *pqqD* | *pqqE* | *pqqF* | *pqq-DH* |
|---|---|---|---|---|---|---|---|---|---|---|---|---|---|---|---|---|
| BCW-200157 | *Rahnella* | DSN | 0 | 0 | 0 | 0 | 0 | 0 | 1 | 5 | 1 | 1 | 1 | 1 | 1 | 2 |
| BCW-200158 | *Lactococcus* | DSN | 0 | 0 | 0 | 0 | 0 | 0 | 0 | 4 | 0 | 0 | 0 | 1 | 0 | 0 |
| BCW-200159 | *Lactococcus* | DSN | 0 | 0 | 0 | 0 | 0 | 0 | 0 | 4 | 0 | 0 | 0 | 1 | 0 | 0 |
| BCW-200160 | *Lactococcus* | DSN | 0 | 0 | 0 | 0 | 0 | 0 | 0 | 4 | 0 | 0 | 0 | 1 | 0 | 0 |
| BCW-200163 | *Lactococcus* | DSN | 0 | 0 | 0 | 0 | 0 | 0 | 0 | 4 | 0 | 0 | 0 | 1 | 0 | 0 |
| BCW-200174 | *Lactococcus* | DSN | 0 | 0 | 0 | 0 | 0 | 0 | 0 | 4 | 0 | 0 | 0 | 1 | 0 | 0 |
| BCW-200175 | *Lactococcus* | DSN | 0 | 0 | 0 | 0 | 0 | 0 | 0 | 4 | 0 | 0 | 0 | 1 | 0 | 0 |
| BCW-200180 | *Lactococcus* | DSN | 0 | 0 | 0 | 0 | 0 | 0 | 0 | 4 | 0 | 0 | 0 | 1 | 0 | 0 |
| BCW-200188 | *Lactococcus* | DSN | 0 | 0 | 0 | 0 | 0 | 0 | 0 | 4 | 0 | 0 | 0 | 1 | 0 | 0 |
| BCW-200192 | *Lactococcus* | DSN | 0 | 0 | 0 | 0 | 0 | 0 | 0 | 4 | 0 | 0 | 0 | 1 | 0 | 0 |
| BCW-200196 | *Lactococcus* | DSN | 0 | 0 | 0 | 0 | 0 | 0 | 0 | 4 | 0 | 0 | 0 | 1 | 0 | 0 |
| BCW-200198 | *Lactococcus* | DSN | 0 | 0 | 0 | 0 | 0 | 0 | 0 | 4 | 0 | 0 | 0 | 0 | 0 | 0 |
| BCW-200208 | *Agrobacterium* | DSN | 0 | 0 | 0 | 0 | 0 | 0 | 0 | 3 | 0 | 0 | 0 | 0 | 0 | 2 |
| BCW-200213 | *Rahnella* | DSN | 0 | 0 | 0 | 0 | 0 | 0 | 2 | 5 | 0 | 0 | 0 | 0 | 0 | 2 |
| BCW-200215 | *Agrobacterium* | DSN | 0 | 0 | 0 | 0 | 0 | 0 | 0 | 3 | 0 | 0 | 0 | 0 | 0 | 2 |
| BCW-200218 | *Enterobacter* | DSN | 0 | 0 | 0 | 0 | 0 | 0 | 1 | 7 | 0 | 0 | 0 | 0 | 0 | 2 |
| BCW-200219 | *Enterobacter* | DSN | 0 | 0 | 0 | 0 | 0 | 0 | 1 | 7 | 0 | 0 | 0 | 0 | 0 | 2 |
| BCW-200229 | *Lactococcus* | DSN | 0 | 0 | 0 | 0 | 0 | 0 | 0 | 4 | 0 | 0 | 0 | 0 | 0 | 0 |
| BCW-200231 | *unassigned* | DSN | 0 | 0 | 0 | 0 | 0 | 0 | 1 | 6 | 0 | 0 | 0 | 0 | 0 | 2 |
| BCW-200232 | *Lactococcus* | DSN | 0 | 0 | 0 | 0 | 0 | 0 | 0 | 4 | 0 | 0 | 0 | 1 | 0 | 0 |
| BCW-200238 | *Lactococcus* | DSN | 0 | 0 | 0 | 0 | 0 | 0 | 0 | 4 | 0 | 0 | 0 | 1 | 0 | 0 |
| BCW-200241 | *Lactococcus* | DSN | 0 | 0 | 0 | 0 | 0 | 0 | 0 | 4 | 0 | 0 | 0 | 0 | 0 | 0 |
| BCW-200268 | *Enterobacter* | DSN | 0 | 0 | 0 | 0 | 0 | 0 | 2 | 8 | 0 | 0 | 0 | 0 | 0 | 2 |
| BCW-200269 | *Lelliottia* | DSN | 0 | 0 | 0 | 0 | 0 | 0 | 1 | 6 | 0 | 0 | 0 | 0 | 0 | 2 |
| BCW-200270 | *Lelliottia* | DSN | 0 | 0 | 0 | 0 | 0 | 0 | 1 | 6 | 0 | 0 | 0 | 0 | 0 | 2 |
| BCW-200271 | *Lelliottia* | DSN | 0 | 0 | 0 | 0 | 0 | 0 | 1 | 6 | 0 | 0 | 0 | 0 | 0 | 2 |

| Isolate ID | Genus | NIF Grp | *nifH* | *nifD* | *nifK* | *nifE* | *nifN* | *nifB* | *acdS* | *ipdC/ ppdC* | *pqqB* | *pqqC* | *pqqD* | *pqqE* | *pqqF* | *pqq-DH* |
|---|---|---|---|---|---|---|---|---|---|---|---|---|---|---|---|---|
| BCW-200272 | *Rahnella* | DSN | 0 | 0 | 0 | 0 | 0 | 0 | 1 | 5 | 1 | 1 | 1 | 1 | 1 | 2 |
| BCW-200275 | *Lelliottia* | DSN | 0 | 0 | 0 | 0 | 0 | 0 | 1 | 6 | 0 | 0 | 0 | 0 | 0 | 2 |
| BCW-200279 | *Enterobacter* | DSN | 0 | 0 | 0 | 0 | 0 | 0 | 2 | 7 | 0 | 0 | 0 | 0 | 0 | 2 |
| BCW-200315 | *unassigned* | DSN | 0 | 0 | 0 | 0 | 0 | 0 | 1 | 8 | 0 | 0 | 0 | 0 | 0 | 2 |
| BCW-200319 | *Erwinia* | DSN | 0 | 0 | 0 | 0 | 0 | 0 | 1 | 6 | 1 | 1 | 1 | 1 | 1 | 3 |
| BCW-200327 | *Serratia* | DSN | 0 | 0 | 0 | 0 | 0 | 0 | 1 | 9 | 0 | 0 | 0 | 0 | 0 | 3 |
| BCW-200328 | *Agrobacterium* | DSN | 0 | 0 | 0 | 0 | 0 | 0 | 0 | 4 | 0 | 0 | 0 | 0 | 0 | 2 |
| BCW-200464 | *Agrobacterium* | DSN | 0 | 0 | 0 | 0 | 0 | 0 | 0 | 3 | 0 | 0 | 0 | 0 | 0 | 2 |
| BCW-200465 | *Agrobacterium* | DSN | 0 | 0 | 0 | 0 | 0 | 0 | 0 | 4 | 0 | 0 | 0 | 0 | 0 | 2 |
| BCW-200471 | *Pseudomonas* | DSN | 0 | 0 | 0 | 0 | 0 | 0 | 1 | 5 | 1 | 1 | 1 | 1 | 1 | 2 |
| BCW-200473 | *Lelliottia* | DSN | 0 | 0 | 0 | 0 | 0 | 0 | 1 | 6 | 0 | 0 | 0 | 0 | 0 | 2 |
| BCW-200533 | *Rhodococcus* | DSN | 0 | 0 | 0 | 0 | 0 | 0 | 1 | 4 | 0 | 0 | 0 | 1 | 0 | 0 |
| BCW-200539 | *Citrobacter* | DSN | 0 | 0 | 0 | 0 | 0 | 0 | 1 | 5 | 0 | 0 | 0 | 0 | 0 | 0 |
| BCW-200542 | *unassigned* | DSN | 0 | 0 | 0 | 0 | 0 | 0 | 1 | 9 | 0 | 0 | 0 | 0 | 0 | 3 |
| BCW-200544 | *Serratia* | DSN | 0 | 0 | 0 | 0 | 0 | 0 | 1 | 9 | 0 | 0 | 0 | 0 | 0 | 3 |
| BCW-200545 | *Rahnella* | DSN | 0 | 0 | 0 | 0 | 0 | 0 | 2 | 5 | 1 | 1 | 1 | 1 | 0 | 2 |
| BCW-200547 | *Serratia* | DSN | 0 | 0 | 0 | 0 | 0 | 0 | 1 | 9 | 0 | 0 | 0 | 0 | 0 | 3 |
| BCW-200556 | *Lelliottia* | DSN | 0 | 0 | 0 | 0 | 0 | 0 | 1 | 6 | 0 | 0 | 0 | 0 | 0 | 2 |
| BCW-200561 | *Rahnella* | DSN | 0 | 0 | 0 | 0 | 0 | 0 | 1 | 5 | 1 | 1 | 1 | 1 | 1 | 2 |
| BCW-200564 | *Rahnella* | DSN | 0 | 0 | 0 | 0 | 0 | 0 | 2 | 5 | 1 | 1 | 1 | 1 | 0 | 2 |
| BCW-200565 | *Rahnella* | DSN | 0 | 0 | 0 | 0 | 0 | 0 | 2 | 5 | 1 | 1 | 1 | 1 | 0 | 2 |
| BCW-200596 | *Lelliottia* | DSN | 0 | 0 | 0 | 0 | 0 | 0 | 1 | 6 | 0 | 0 | 0 | 0 | 0 | 2 |
| BCW-200634 | *Lelliottia* | DSN | 0 | 0 | 0 | 0 | 0 | 0 | 1 | 6 | 0 | 0 | 0 | 0 | 0 | 2 |
| BCW-200641 | *Lelliottia* | DSN | 0 | 0 | 0 | 0 | 0 | 0 | 1 | 6 | 0 | 0 | 0 | 0 | 0 | 2 |
| BCW-200642 | *Rahnella* | DSN | 0 | 0 | 0 | 0 | 0 | 0 | 2 | 5 | 1 | 1 | 1 | 1 | 0 | 2 |
| BCW-200649 | *Rahnella* | DSN | 0 | 0 | 0 | 0 | 0 | 0 | 2 | 5 | 1 | 1 | 1 | 1 | 0 | 2 |

| Isolate ID | Genus | NIF Grp | *nifH* | *nifD* | *nifK* | *nifE* | *nifN* | *nifB* | *acdS* | *ipdC/ ppdC* | *pqqB* | *pqqC* | *pqqD* | *pqqE* | *pqqF* | *pqq-DH* |
|---|---|---|---|---|---|---|---|---|---|---|---|---|---|---|---|---|
| BCW-200663 | *Agrobacterium* | DSN | 0 | 0 | 0 | 0 | 0 | 0 | 0 | 3 | 0 | 0 | 0 | 0 | 0 | 2 |
| BCW-200705 | *Agrobacterium* | DSN | 0 | 0 | 0 | 0 | 0 | 0 | 0 | 4 | 0 | 0 | 0 | 0 | 0 | 2 |
| BCW-200715 | *Rahnella* | DSN | 0 | 0 | 0 | 0 | 0 | 0 | 2 | 5 | 1 | 1 | 1 | 1 | 0 | 2 |
| BCW-200716 | *Rahnella* | DSN | 0 | 0 | 0 | 0 | 0 | 0 | 2 | 5 | 1 | 1 | 1 | 1 | 0 | 2 |
| BCW-200718 | *Rahnella* | DSN | 0 | 0 | 0 | 0 | 0 | 0 | 2 | 5 | 1 | 1 | 1 | 1 | 0 | 2 |
| BCW-200723 | *Rahnella* | DSN | 0 | 0 | 0 | 0 | 0 | 0 | 2 | 5 | 1 | 1 | 1 | 1 | 0 | 2 |
| BCW-200724 | *Rahnella* | DSN | 0 | 0 | 0 | 0 | 0 | 0 | 2 | 5 | 1 | 1 | 1 | 1 | 0 | 2 |
| BCW-200725 | *Rahnella* | DSN | 0 | 0 | 0 | 0 | 0 | 0 | 2 | 5 | 1 | 1 | 1 | 1 | 0 | 2 |
| BCW-200726 | *Rahnella* | DSN | 0 | 0 | 0 | 0 | 0 | 0 | 2 | 5 | 1 | 1 | 1 | 1 | 0 | 2 |
| BCW-200736 | *Rahnella* | DSN | 0 | 0 | 0 | 0 | 0 | 0 | 2 | 5 | 1 | 1 | 1 | 1 | 0 | 2 |
| BCW-200806 | *Rahnella* | DSN | 0 | 0 | 0 | 0 | 0 | 0 | 1 | 5 | 1 | 1 | 1 | 1 | 1 | 2 |
| BCW-200808 | *Rahnella* | DSN | 0 | 0 | 0 | 0 | 0 | 0 | 1 | 5 | 1 | 1 | 1 | 1 | 1 | 2 |
| BCW-200810 | *Rahnella* | DSN | 0 | 0 | 0 | 0 | 0 | 0 | 1 | 5 | 1 | 1 | 1 | 1 | 1 | 2 |
| BCW-200814 | *Rahnella* | DSN | 0 | 0 | 0 | 0 | 0 | 0 | 2 | 5 | 1 | 1 | 1 | 1 | 0 | 2 |
| BCW-200815 | *Rahnella* | DSN | 0 | 0 | 0 | 0 | 0 | 0 | 2 | 5 | 1 | 1 | 1 | 1 | 0 | 2 |
| BCW-200883 | *Enterobacter* | DSN | 0 | 0 | 0 | 0 | 0 | 0 | 1 | 7 | 0 | 0 | 0 | 0 | 0 | 2 |
| BCW-200902 | *Agrobacterium* | DSN | 0 | 0 | 0 | 0 | 0 | 0 | 0 | 4 | 0 | 0 | 0 | 0 | 0 | 2 |
| BCW-200903 | *unassigned* | DSN | 0 | 0 | 0 | 0 | 0 | 0 | 0 | 4 | 0 | 0 | 0 | 0 | 0 | 2 |
| BCW-200904 | *Agrobacterium* | DSN | 0 | 0 | 0 | 0 | 0 | 0 | 0 | 4 | 0 | 0 | 0 | 0 | 0 | 2 |
| BCW-200910 | *Agrobacterium* | DSN | 0 | 0 | 0 | 0 | 0 | 0 | 0 | 4 | 0 | 0 | 0 | 0 | 0 | 2 |
| BCW-200912 | *unassigned* | DSN | 0 | 0 | 0 | 0 | 0 | 0 | 0 | 4 | 0 | 0 | 0 | 0 | 0 | 2 |
| BCW-200920 | *Agrobacterium* | DSN | 0 | 0 | 0 | 0 | 0 | 0 | 0 | 4 | 0 | 0 | 0 | 0 | 0 | 2 |
| BCW-200952 | *Pantoea* | DSN | 0 | 0 | 0 | 0 | 0 | 0 | 1 | 5 | 1 | 1 | 1 | 1 | 1 | 2 |
| BCW-200983 | *Citrobacter* | DSN | 0 | 0 | 0 | 0 | 0 | 0 | 1 | 5 | 0 | 0 | 0 | 0 | 0 | 0 |
| BCW-200984 | *Citrobacter* | DSN | 0 | 0 | 0 | 0 | 0 | 0 | 1 | 5 | 0 | 0 | 0 | 0 | 0 | 0 |
| BCW-200986 | *Citrobacter* | DSN | 0 | 0 | 0 | 0 | 0 | 0 | 1 | 5 | 0 | 0 | 0 | 0 | 0 | 0 |

| Isolate ID | Genus | NIF Grp | *nifH* | *nifD* | *nifK* | *nifE* | *nifN* | *nifB* | *acdS* | *ipdC/ ppdC* | *pqqB* | *pqqC* | *pqqD* | *pqqE* | *pqqF* | *pqq-DH* |
|---|---|---|---|---|---|---|---|---|---|---|---|---|---|---|---|---|
| BCW-200988 | *unassigned* | DSN | 0 | 0 | 0 | 0 | 0 | 0 | 1 | 5 | 0 | 0 | 0 | 0 | 0 | 0 |
| BCW-200989 | *Lelliottia* | DSN | 0 | 0 | 0 | 0 | 0 | 0 | 1 | 6 | 0 | 0 | 0 | 0 | 0 | 2 |
| BCW-200990 | *Lelliottia* | DSN | 0 | 0 | 0 | 0 | 0 | 0 | 1 | 6 | 0 | 0 | 0 | 0 | 0 | 2 |
| BCW-200991 | *Lelliottia* | DSN | 0 | 0 | 0 | 0 | 0 | 0 | 1 | 6 | 0 | 0 | 0 | 0 | 0 | 2 |
| BCW-200994 | *Lelliottia* | DSN | 0 | 0 | 0 | 0 | 0 | 0 | 1 | 6 | 0 | 0 | 0 | 0 | 0 | 2 |
| BCW-201007 | *Rahnella* | DSN | 0 | 0 | 0 | 0 | 0 | 0 | 2 | 5 | 1 | 1 | 1 | 1 | 0 | 2 |
| BCW-201008 | *Rahnella* | DSN | 0 | 0 | 0 | 0 | 0 | 0 | 2 | 5 | 1 | 1 | 1 | 1 | 0 | 2 |
| BCW-201010 | *Rahnella* | DSN | 0 | 0 | 0 | 0 | 0 | 0 | 2 | 5 | 1 | 1 | 1 | 1 | 0 | 2 |
| BCW-201013 | *Rahnella* | DSN | 0 | 0 | 0 | 0 | 0 | 0 | 2 | 5 | 1 | 1 | 1 | 1 | 0 | 2 |
| BCW-201014 | *Rahnella* | DSN | 0 | 0 | 0 | 0 | 0 | 0 | 2 | 5 | 1 | 1 | 1 | 1 | 0 | 2 |
| BCW-201024 | *Rahnella* | DSN | 0 | 0 | 0 | 0 | 0 | 0 | 2 | 5 | 1 | 1 | 1 | 1 | 0 | 2 |
| BCW-201025 | *Rahnella* | DSN | 0 | 0 | 0 | 0 | 0 | 0 | 2 | 5 | 1 | 1 | 1 | 1 | 0 | 2 |
| BCW-201028 | *Rahnella* | DSN | 0 | 0 | 0 | 0 | 0 | 0 | 2 | 5 | 1 | 1 | 1 | 1 | 0 | 2 |
| BCW-201036 | *Rahnella* | DSN | 0 | 0 | 0 | 0 | 0 | 0 | 2 | 5 | 1 | 1 | 1 | 1 | 0 | 2 |
| BCW-201045 | *Lelliottia* | DSN | 0 | 0 | 0 | 0 | 0 | 0 | 1 | 6 | 0 | 0 | 0 | 0 | 0 | 2 |
| BCW-201051 | *Serratia* | DSN | 0 | 0 | 0 | 0 | 0 | 0 | 1 | 9 | 0 | 0 | 0 | 0 | 0 | 1 |
| BCW-201054 | *Serratia* | DSN | 0 | 0 | 0 | 0 | 0 | 0 | 1 | 9 | 0 | 0 | 0 | 0 | 0 | 1 |
| BCW-201056 | *Hafnia* | DSN | 0 | 0 | 0 | 0 | 0 | 0 | 0 | 5 | 0 | 0 | 0 | 0 | 0 | 0 |
| BCW-201079 | *Serratia* | DSN | 0 | 0 | 0 | 0 | 0 | 0 | 1 | 9 | 0 | 0 | 0 | 0 | 0 | 1 |
| BCW-201081 | *Pantoea* | DSN | 0 | 0 | 0 | 0 | 0 | 0 | 1 | 7 | 1 | 1 | 1 | 1 | 1 | 3 |
| BCW-201083 | *Citrobacter* | DSN | 0 | 0 | 0 | 0 | 0 | 0 | 1 | 6 | 0 | 0 | 0 | 0 | 0 | 0 |
| BCW-201084 | *Lelliottia* | DSN | 0 | 0 | 0 | 0 | 0 | 0 | 1 | 6 | 0 | 0 | 0 | 0 | 0 | 2 |
| BCW-201085 | *Serratia* | DSN | 0 | 0 | 0 | 0 | 0 | 0 | 1 | 9 | 0 | 0 | 0 | 0 | 0 | 3 |
| BCW-201090 | *unassigned* | DSN | 0 | 0 | 0 | 0 | 0 | 0 | 1 | 6 | 0 | 0 | 0 | 0 | 0 | 2 |
| BCW-201103 | *Lelliottia* | DSN | 0 | 0 | 0 | 0 | 0 | 0 | 1 | 6 | 0 | 0 | 0 | 0 | 0 | 2 |
| BCW-201151 | *Lelliottia* | DSN | 0 | 0 | 0 | 0 | 0 | 0 | 1 | 6 | 0 | 0 | 0 | 0 | 0 | 2 |

| Isolate ID | Genus | NIF Grp | *nifH* | *nifD* | *nifK* | *nifE* | *nifN* | *nifB* | *acdS* | *ipdC/ ppdC* | *pqqB* | *pqqC* | *pqqD* | *pqqE* | *pqqF* | *pqq-DH* |
|---|---|---|---|---|---|---|---|---|---|---|---|---|---|---|---|---|
| BCW-201152 | *Citrobacter* | DSN | 0 | 0 | 0 | 0 | 0 | 0 | 1 | 6 | 0 | 0 | 0 | 0 | 0 | 0 |
| BCW-201153 | *Serratia* | DSN | 0 | 0 | 0 | 0 | 0 | 0 | 1 | 9 | 0 | 0 | 0 | 0 | 0 | 3 |
| BCW-201154 | *Citrobacter* | DSN | 0 | 0 | 0 | 0 | 0 | 0 | 1 | 6 | 0 | 0 | 0 | 0 | 0 | 0 |
| BCW-201173 | *Escherichia* | DSN | 0 | 0 | 0 | 0 | 0 | 0 | 1 | 6 | 0 | 0 | 0 | 0 | 0 | 1 |
| BCW-201175 | *Rahnella* | DSN | 0 | 0 | 0 | 0 | 0 | 0 | 2 | 5 | 1 | 1 | 1 | 1 | 0 | 2 |
| BCW-201176 | *Rahnella* | DSN | 0 | 0 | 0 | 0 | 0 | 0 | 2 | 5 | 1 | 1 | 1 | 1 | 0 | 2 |
| BCW-201185 | *Serratia* | DSN | 0 | 0 | 0 | 0 | 0 | 0 | 1 | 9 | 0 | 0 | 0 | 0 | 0 | 3 |
| BCW-201186 | *Rahnella* | DSN | 0 | 0 | 0 | 0 | 0 | 0 | 2 | 5 | 1 | 1 | 1 | 1 | 0 | 2 |
| BCW-201187 | *Rahnella* | DSN | 0 | 0 | 0 | 0 | 0 | 0 | 2 | 5 | 1 | 1 | 1 | 1 | 0 | 2 |
| BCW-201236 | *Hafnia* | DSN | 0 | 0 | 0 | 0 | 0 | 0 | 0 | 6 | 0 | 0 | 0 | 0 | 0 | 0 |
| BCW-201238 | *Serratia* | DSN | 0 | 0 | 0 | 0 | 0 | 0 | 1 | 9 | 0 | 0 | 0 | 0 | 0 | 3 |
| BCW-201245 | *Rahnella* | DSN | 0 | 0 | 0 | 0 | 0 | 0 | 1 | 5 | 1 | 1 | 1 | 1 | 1 | 2 |
| BCW-201248 | *Rahnella* | DSN | 0 | 0 | 0 | 0 | 0 | 0 | 1 | 5 | 1 | 1 | 1 | 1 | 1 | 2 |
| BCW-201257 | *Serratia* | DSN | 0 | 0 | 0 | 0 | 0 | 0 | 1 | 9 | 0 | 0 | 0 | 0 | 0 | 3 |
| BCW-201258 | *Lelliottia* | DSN | 0 | 0 | 0 | 0 | 0 | 0 | 1 | 6 | 0 | 0 | 0 | 0 | 0 | 2 |
| BCW-201260 | *Lelliottia* | DSN | 0 | 0 | 0 | 0 | 0 | 0 | 1 | 6 | 0 | 0 | 0 | 0 | 0 | 2 |
| BCW-201350 | *Serratia* | DSN | 0 | 0 | 0 | 0 | 0 | 0 | 1 | 9 | 0 | 0 | 0 | 0 | 0 | 3 |
| BCW-201444 | *Atlantibacter* | DSN | 0 | 0 | 0 | 0 | 0 | 0 | 1 | 5 | 0 | 0 | 0 | 0 | 0 | 1 |
| BCW-201445 | *Atlantibacter* | DSN | 0 | 0 | 0 | 0 | 0 | 0 | 1 | 5 | 0 | 0 | 0 | 0 | 0 | 1 |
| BCW-201450 | *Hafnia* | DSN | 0 | 0 | 0 | 0 | 0 | 0 | 0 | 5 | 0 | 0 | 0 | 0 | 0 | 0 |
| BCW-201453 | *Lactococcus* | DSN | 0 | 0 | 0 | 0 | 0 | 0 | 0 | 4 | 0 | 0 | 0 | 0 | 0 | 0 |
| BCW-201648 | *Rahnella* | DSN | 0 | 0 | 0 | 0 | 0 | 0 | 2 | 5 | 1 | 1 | 1 | 1 | 0 | 2 |
| BCW-201653 | *Serratia* | DSN | 0 | 0 | 0 | 0 | 0 | 0 | 1 | 9 | 0 | 0 | 0 | 0 | 0 | 1 |
| BCW-201654 | *Rahnella* | DSN | 0 | 0 | 0 | 0 | 0 | 0 | 2 | 5 | 1 | 1 | 1 | 1 | 0 | 2 |
| BCW-201662 | *Serratia* | DSN | 0 | 0 | 0 | 0 | 0 | 0 | 1 | 9 | 0 | 0 | 0 | 0 | 0 | 1 |
| BCW-201726 | *Enterobacter* | DSN | 0 | 0 | 0 | 0 | 0 | 0 | 1 | 7 | 0 | 0 | 0 | 0 | 0 | 2 |

| Isolate ID | Genus | NIF Grp | *nifH* | *nifD* | *nifK* | *nifE* | *nifN* | *nifB* | *acdS* | *ipdC/ ppdC* | *pqqB* | *pqqC* | *pqqD* | *pqqE* | *pqqF* | *pqq-DH* |
|---|---|---|---|---|---|---|---|---|---|---|---|---|---|---|---|---|
| BCW-201809 | *Serratia* | DSN | 0 | 0 | 0 | 0 | 0 | 0 | 1 | 9 | 0 | 0 | 0 | 0 | 0 | 3 |
| BCW-201811 | *Enterobacter* | DSN | 0 | 0 | 0 | 0 | 0 | 0 | 1 | 7 | 0 | 0 | 0 | 0 | 0 | 2 |
| BCW-201812 | *Enterobacter* | DSN | 0 | 0 | 0 | 0 | 0 | 0 | 1 | 7 | 0 | 0 | 0 | 0 | 0 | 2 |
| BCW-201814 | *Enterobacter* | DSN | 0 | 0 | 0 | 0 | 0 | 0 | 1 | 7 | 0 | 0 | 0 | 0 | 0 | 2 |
| BCW-201826 | *Pantoea* | DSN | 0 | 0 | 0 | 0 | 0 | 0 | 1 | 5 | 1 | 1 | 1 | 1 | 1 | 2 |
| BCW-201827 | *Pantoea* | DSN | 0 | 0 | 0 | 0 | 0 | 0 | 1 | 7 | 1 | 1 | 1 | 1 | 1 | 3 |
| BCW-201832 | *Enterobacter* | DSN | 0 | 0 | 0 | 0 | 0 | 0 | 1 | 7 | 0 | 0 | 0 | 0 | 0 | 2 |
| BCW-201833 | *Pantoea* | DSN | 0 | 0 | 0 | 0 | 0 | 0 | 1 | 5 | 1 | 1 | 1 | 1 | 1 | 4 |
| BCW-201835 | *Staphylococcus* | DSN | 0 | 0 | 0 | 0 | 0 | 0 | 0 | 4 | 0 | 0 | 0 | 0 | 0 | 0 |
| BCW-201839 | *Enterobacter* | DSN | 0 | 0 | 0 | 0 | 0 | 0 | 1 | 7 | 0 | 0 | 0 | 0 | 0 | 2 |
| BCW-201842 | *Curtobacterium* | DSN | 0 | 0 | 0 | 0 | 0 | 0 | 0 | 2 | 0 | 0 | 0 | 0 | 0 | 0 |
| BCW-201845 | *unassigned* | DSN | 0 | 0 | 0 | 0 | 0 | 0 | 1 | 7 | 0 | 0 | 0 | 0 | 0 | 2 |
| BCW-201847 | *Enterobacter* | DSN | 0 | 0 | 0 | 0 | 0 | 0 | 1 | 7 | 0 | 0 | 0 | 0 | 0 | 2 |
| BCW-201848 | *Pantoea* | DSN | 0 | 0 | 0 | 0 | 0 | 0 | 1 | 5 | 1 | 1 | 1 | 1 | 1 | 2 |
| BCW-201849 | *Enterobacter* | DSN | 0 | 0 | 0 | 0 | 0 | 0 | 1 | 7 | 0 | 0 | 0 | 0 | 0 | 2 |
| BCW-201853 | *Erwinia* | DSN | 0 | 0 | 0 | 0 | 0 | 0 | 1 | 6 | 1 | 1 | 1 | 1 | 1 | 3 |
| BCW-201854 | *Erwinia* | DSN | 0 | 0 | 0 | 0 | 0 | 0 | 1 | 6 | 1 | 1 | 1 | 1 | 1 | 3 |
| BCW-201857 | *Enterobacter* | DSN | 0 | 0 | 0 | 0 | 0 | 0 | 1 | 7 | 0 | 0 | 0 | 0 | 0 | 2 |
| BCW-201861 | *Lactococcus* | DSN | 0 | 0 | 0 | 0 | 0 | 0 | 0 | 4 | 0 | 0 | 0 | 0 | 0 | 0 |
| BCW-201864 | *Pantoea* | DSN | 0 | 0 | 0 | 0 | 0 | 0 | 1 | 7 | 1 | 1 | 1 | 1 | 1 | 3 |
| BCW-201865 | *Erwinia* | DSN | 0 | 0 | 0 | 0 | 0 | 0 | 1 | 6 | 1 | 1 | 1 | 1 | 1 | 3 |
| BCW-201866 | *Enterobacter* | DSN | 0 | 0 | 0 | 0 | 0 | 0 | 1 | 7 | 0 | 0 | 0 | 0 | 0 | 2 |
| BCW-201867 | *Pantoea* | DSN | 0 | 0 | 0 | 0 | 0 | 0 | 1 | 7 | 1 | 1 | 1 | 1 | 1 | 3 |
| BCW-201874 | *Enterobacter* | DSN | 0 | 0 | 0 | 0 | 0 | 0 | 1 | 7 | 0 | 0 | 0 | 0 | 0 | 2 |
| BCW-201877 | *Enterobacter* | DSN | 0 | 0 | 0 | 0 | 0 | 0 | 1 | 7 | 0 | 0 | 0 | 0 | 0 | 2 |
| BCW-201878 | *Enterobacter* | DSN | 0 | 0 | 0 | 0 | 0 | 0 | 1 | 7 | 0 | 0 | 0 | 0 | 0 | 2 |

| **Isolate ID** | **Genus** | **NIF Grp** | ***nifH*** | ***nifD*** | ***nifK*** | ***nifE*** | ***nifN*** | ***nifB*** | ***acdS*** | ***ipdC/ ppdC*** | ***pqqB*** | ***pqqC*** | ***pqqD*** | ***pqqE*** | ***pqqF*** | ***pqq-DH*** |
|---|---|---|---|---|---|---|---|---|---|---|---|---|---|---|---|---|
| BCW-201880 | *Enterobacter* | DSN | 0 | 0 | 0 | 0 | 0 | 0 | 1 | 7 | 0 | 0 | 0 | 0 | 0 | 2 |
| BCW-201881 | *Enterobacter* | DSN | 0 | 0 | 0 | 0 | 0 | 0 | 1 | 7 | 0 | 0 | 0 | 0 | 0 | 2 |
| BCW-201882 | *Enterobacter* | DSN | 0 | 0 | 0 | 0 | 0 | 0 | 1 | 7 | 0 | 0 | 0 | 0 | 0 | 2 |
| BCW-201883 | *unassigned* | DSN | 0 | 0 | 0 | 0 | 0 | 0 | 1 | 7 | 0 | 0 | 0 | 0 | 0 | 2 |
| BCW-201884 | *Enterobacter* | DSN | 0 | 0 | 0 | 0 | 0 | 0 | 1 | 7 | 0 | 0 | 0 | 0 | 0 | 2 |
| BCW-201885 | *Enterobacter* | DSN | 0 | 0 | 0 | 0 | 0 | 0 | 1 | 7 | 0 | 0 | 0 | 0 | 0 | 2 |
| BCW-201889 | *Enterobacter* | DSN | 0 | 0 | 0 | 0 | 0 | 0 | 1 | 7 | 0 | 0 | 0 | 0 | 0 | 2 |
| BCW-201891 | *unassigned* | DSN | 0 | 0 | 0 | 0 | 0 | 0 | 1 | 9 | 0 | 0 | 0 | 0 | 0 | 3 |
| BCW-201895 | *Enterobacter* | DSN | 0 | 0 | 0 | 0 | 0 | 0 | 1 | 7 | 0 | 0 | 0 | 0 | 0 | 2 |
| BCW-201896 | *Curtobacterium* | DSN | 0 | 0 | 0 | 0 | 0 | 0 | 0 | 2 | 0 | 0 | 0 | 0 | 0 | 0 |
| BCW-201897 | *Pantoea* | DSN | 0 | 0 | 0 | 0 | 0 | 0 | 1 | 5 | 1 | 1 | 1 | 1 | 1 | 2 |
| BCW-201899 | *Enterobacter* | DSN | 0 | 0 | 0 | 0 | 0 | 0 | 1 | 7 | 0 | 0 | 0 | 0 | 0 | 2 |
| BCW-201903 | *Enterobacter* | DSN | 0 | 0 | 0 | 0 | 0 | 0 | 1 | 7 | 0 | 0 | 0 | 0 | 0 | 2 |
| BCW-201909 | *Pantoea* | DSN | 0 | 0 | 0 | 0 | 0 | 0 | 1 | 7 | 1 | 1 | 1 | 1 | 1 | 3 |
| BCW-201914 | *Enterobacter* | DSN | 0 | 0 | 0 | 0 | 0 | 0 | 1 | 7 | 0 | 0 | 0 | 0 | 0 | 2 |
| BCW-201917 | *Pantoea* | DSN | 0 | 0 | 0 | 0 | 0 | 0 | 1 | 7 | 1 | 1 | 1 | 1 | 1 | 3 |
| BCW-201933 | *Enterobacter* | DSN | 0 | 0 | 0 | 0 | 0 | 0 | 1 | 7 | 0 | 0 | 0 | 0 | 0 | 2 |
| BCW-201938 | *Enterobacter* | DSN | 0 | 0 | 0 | 0 | 0 | 0 | 1 | 7 | 0 | 0 | 0 | 0 | 0 | 2 |
| BCW-201945 | *Enterobacter* | DSN | 0 | 0 | 0 | 0 | 0 | 0 | 1 | 7 | 0 | 0 | 0 | 0 | 0 | 2 |
| BCW-201949 | *Enterobacter* | DSN | 0 | 0 | 0 | 0 | 0 | 0 | 1 | 7 | 0 | 0 | 0 | 0 | 0 | 2 |
| BCW-201957 | *Enterobacter* | DSN | 0 | 0 | 0 | 0 | 0 | 0 | 1 | 7 | 0 | 0 | 0 | 0 | 0 | 2 |
| BCW-201975 | *Enterobacter* | DSN | 0 | 0 | 0 | 0 | 0 | 0 | 1 | 7 | 0 | 0 | 0 | 0 | 0 | 2 |
| BCW-201982 | *Rahnella* | DSN | 0 | 0 | 0 | 0 | 0 | 0 | 2 | 5 | 1 | 1 | 1 | 1 | 0 | 2 |
| BCW-201995 | *Pantoea* | DSN | 0 | 0 | 0 | 0 | 0 | 0 | 1 | 5 | 1 | 1 | 1 | 1 | 1 | 2 |
| BCW-201997 | *Pantoea* | DSN | 0 | 0 | 0 | 0 | 0 | 0 | 1 | 7 | 1 | 1 | 1 | 1 | 1 | 4 |
| BCW-202001 | *Pantoea* | DSN | 0 | 0 | 0 | 0 | 0 | 0 | 1 | 7 | 1 | 1 | 1 | 1 | 1 | 4 |

| Isolate ID | Genus | NIF Grp | *nifH* | *nifD* | *nifK* | *nifE* | *nifN* | *nifB* | *acdS* | *ipdC/ ppdC* | *pqqB* | *pqqC* | *pqqD* | *pqqE* | *pqqF* | *pqq-DH* |
|---|---|---|---|---|---|---|---|---|---|---|---|---|---|---|---|---|
| BCW-200014 | *Enterobacter* | DSP | 1 | 2 | 2 | 3 | 2 | 1 | 1 | 7 | 0 | 0 | 0 | 0 | 0 | 2 |
| BCW-200016 | *Enterobacter* | DSP | 1 | 2 | 2 | 3 | 2 | 1 | 1 | 7 | 0 | 0 | 0 | 1 | 0 | 2 |
| BCW-200017 | *Enterobacter* | DSP | 1 | 2 | 2 | 3 | 2 | 1 | 1 | 7 | 0 | 0 | 0 | 1 | 0 | 2 |
| BCW-200026 | *Enterobacter* | DSP | 1 | 2 | 2 | 3 | 2 | 1 | 1 | 7 | 0 | 0 | 0 | 1 | 0 | 2 |
| BCW-200028 | *unassigned* | DSP | 2 | 3 | 3 | 5 | 4 | 1 | 2 | 7 | 0 | 0 | 0 | 0 | 0 | 3 |
| BCW-200030 | *Raoultella* | DSP | 2 | 3 | 3 | 5 | 4 | 1 | 2 | 7 | 0 | 0 | 0 | 0 | 0 | 3 |
| BCW-200034 | *Enterobacter* | DSP | 1 | 2 | 2 | 3 | 2 | 1 | 1 | 7 | 0 | 0 | 0 | 0 | 0 | 2 |
| BCW-200035 | *Enterobacter* | DSP | 1 | 2 | 2 | 3 | 2 | 1 | 1 | 7 | 0 | 0 | 0 | 0 | 0 | 2 |
| BCW-200049 | *Klebsiella* | DSP | 1 | 2 | 2 | 3 | 2 | 1 | 1 | 8 | 0 | 0 | 0 | 0 | 0 | 3 |
| BCW-200050 | *Enterobacter* | DSP | 1 | 2 | 2 | 3 | 2 | 1 | 1 | 7 | 0 | 0 | 0 | 0 | 0 | 2 |
| BCW-200053 | *Klebsiella* | DSP | 1 | 2 | 2 | 3 | 2 | 1 | 1 | 8 | 0 | 0 | 0 | 0 | 0 | 3 |
| BCW-200069 | *Klebsiella* | DSP | 1 | 2 | 2 | 3 | 3 | 1 | 1 | 7 | 0 | 0 | 0 | 0 | 0 | 3 |
| BCW-200083 | *Klebsiella* | DSP | 1 | 2 | 2 | 3 | 3 | 1 | 1 | 7 | 0 | 0 | 0 | 0 | 0 | 3 |
| BCW-200084 | *Klebsiella* | DSP | 1 | 2 | 2 | 3 | 3 | 1 | 1 | 7 | 0 | 0 | 0 | 0 | 0 | 3 |
| BCW-200086 | *Klebsiella* | DSP | 1 | 2 | 2 | 3 | 3 | 1 | 1 | 7 | 0 | 0 | 0 | 0 | 0 | 3 |
| BCW-200093 | *Klebsiella* | DSP | 1 | 2 | 2 | 3 | 3 | 1 | 1 | 7 | 0 | 0 | 0 | 0 | 0 | 3 |
| BCW-200095 | *Enterobacter* | DSP | 1 | 2 | 2 | 3 | 2 | 1 | 1 | 7 | 0 | 0 | 0 | 0 | 0 | 2 |
| BCW-200096 | *Klebsiella* | DSP | 1 | 2 | 2 | 3 | 2 | 1 | 1 | 7 | 0 | 0 | 0 | 0 | 0 | 3 |
| BCW-200099 | *Klebsiella* | DSP | 1 | 2 | 2 | 3 | 2 | 1 | 1 | 7 | 0 | 0 | 0 | 0 | 0 | 3 |
| BCW-200102 | *Enterobacter* | DSP | 1 | 2 | 2 | 3 | 2 | 1 | 1 | 7 | 0 | 0 | 0 | 0 | 0 | 2 |
| BCW-200106 | *Raoultella* | DSP | 1 | 2 | 2 | 3 | 3 | 1 | 1 | 7 | 0 | 0 | 0 | 0 | 0 | 3 |
| BCW-200109 | *Enterobacter* | DSP | 1 | 2 | 2 | 3 | 2 | 1 | 1 | 7 | 0 | 0 | 0 | 0 | 0 | 2 |
| BCW-200113 | *unassigned* | DSP | 1 | 2 | 2 | 3 | 3 | 1 | 1 | 7 | 0 | 0 | 0 | 0 | 0 | 3 |
| BCW-200117 | *Raoultella* | DSP | 1 | 2 | 2 | 3 | 3 | 1 | 1 | 7 | 0 | 0 | 0 | 0 | 0 | 3 |
| BCW-200120 | *Raoultella* | DSP | 1 | 2 | 2 | 3 | 3 | 1 | 1 | 7 | 0 | 0 | 0 | 0 | 0 | 3 |
| BCW-200122 | *Enterobacter* | DSP | 1 | 2 | 2 | 3 | 2 | 1 | 1 | 7 | 0 | 0 | 0 | 0 | 0 | 2 |

| Isolate ID | Genus | NIF Grp | ***nifH*** | ***nifD*** | ***nifK*** | ***nifE*** | ***nifN*** | ***nifB*** | ***acdS*** | ***ipdC/ ppdC*** | ***pqqB*** | ***pqqC*** | ***pqqD*** | ***pqqE*** | ***pqqF*** | ***pqq-DH*** |
|---|---|---|---|---|---|---|---|---|---|---|---|---|---|---|---|---|
| BCW-200123 | *Klebsiella* | DSP | 1 | 2 | 2 | 3 | 3 | 1 | 1 | 7 | 0 | 0 | 0 | 0 | 0 | 3 |
| BCW-200124 | *Klebsiella* | DSP | 1 | 2 | 2 | 3 | 3 | 1 | 1 | 7 | 0 | 0 | 0 | 0 | 0 | 3 |
| BCW-200129 | *Klebsiella* | DSP | 1 | 2 | 2 | 3 | 3 | 1 | 1 | 7 | 0 | 0 | 0 | 0 | 0 | 3 |
| BCW-200132 | *Klebsiella* | DSP | 1 | 2 | 2 | 3 | 3 | 1 | 1 | 7 | 0 | 0 | 0 | 0 | 0 | 3 |
| BCW-200133 | *unassigned* | DSP | 1 | 2 | 2 | 3 | 2 | 1 | 1 | 7 | 0 | 0 | 0 | 0 | 0 | 2 |
| BCW-200136 | *Klebsiella* | DSP | 1 | 2 | 2 | 3 | 3 | 1 | 1 | 7 | 0 | 0 | 0 | 0 | 0 | 3 |
| BCW-200137 | *Klebsiella* | DSP | 1 | 2 | 2 | 3 | 3 | 1 | 1 | 7 | 0 | 0 | 0 | 0 | 0 | 3 |
| BCW-200141 | *Kosakonia* | DSP | 1 | 2 | 2 | 3 | 2 | 1 | 2 | 7 | 0 | 0 | 0 | 0 | 0 | 1 |
| BCW-200142 | *Raoultella* | DSP | 1 | 2 | 2 | 3 | 3 | 1 | 1 | 7 | 0 | 0 | 0 | 0 | 0 | 3 |
| BCW-200148 | *unassigned* | DSP | 1 | 2 | 2 | 2 | 3 | 1 | 2 | 7 | 0 | 0 | 0 | 0 | 0 | 3 |
| BCW-200156 | *unassigned* | DSP | 1 | 2 | 1 | 2 | 2 | 1 | 2 | 7 | 0 | 0 | 0 | 0 | 0 | 3 |
| BCW-200161 | *Raoultella* | DSP | 1 | 2 | 2 | 3 | 3 | 1 | 1 | 7 | 0 | 0 | 0 | 0 | 0 | 3 |
| BCW-200162 | *Metakosakonia* | DSP | 1 | 2 | 2 | 2 | 2 | 1 | 2 | 5 | 0 | 0 | 0 | 0 | 0 | 2 |
| BCW-200165 | *Raoultella* | DSP | 1 | 2 | 2 | 3 | 3 | 1 | 1 | 7 | 0 | 0 | 0 | 0 | 0 | 3 |
| BCW-200167 | *Klebsiella* | DSP | 1 | 2 | 2 | 3 | 3 | 1 | 1 | 7 | 0 | 0 | 0 | 0 | 0 | 3 |
| BCW-200168 | *Metakosakonia* | DSP | 1 | 2 | 2 | 2 | 2 | 1 | 2 | 5 | 0 | 0 | 0 | 0 | 0 | 2 |
| BCW-200169 | *Raoultella* | DSP | 1 | 2 | 2 | 3 | 3 | 1 | 1 | 7 | 0 | 0 | 0 | 0 | 0 | 3 |
| BCW-200171 | *Raoultella* | DSP | 1 | 2 | 2 | 3 | 3 | 1 | 1 | 7 | 0 | 0 | 0 | 0 | 0 | 3 |
| BCW-200172 | *Klebsiella* | DSP | 1 | 2 | 2 | 3 | 3 | 1 | 1 | 7 | 0 | 0 | 0 | 0 | 0 | 3 |
| BCW-200177 | *Klebsiella* | DSP | 1 | 2 | 2 | 3 | 3 | 1 | 1 | 7 | 0 | 0 | 0 | 0 | 0 | 3 |
| BCW-200181 | *Kosakonia* | DSP | 1 | 2 | 2 | 3 | 2 | 1 | 2 | 7 | 0 | 0 | 0 | 0 | 0 | 1 |
| BCW-200182 | *unassigned* | DSP | 1 | 2 | 2 | 2 | 3 | 1 | 2 | 7 | 0 | 0 | 0 | 0 | 0 | 3 |
| BCW-200183 | *Kosakonia* | DSP | 1 | 2 | 2 | 3 | 2 | 1 | 2 | 7 | 0 | 0 | 0 | 0 | 0 | 1 |
| BCW-200184 | *unassigned* | DSP | 1 | 2 | 2 | 2 | 3 | 1 | 2 | 7 | 0 | 0 | 0 | 0 | 0 | 3 |
| BCW-200194 | *Kosakonia* | DSP | 1 | 2 | 2 | 3 | 2 | 1 | 2 | 7 | 0 | 0 | 0 | 0 | 0 | 1 |
| BCW-200195 | *unassigned* | DSP | 1 | 2 | 2 | 2 | 3 | 1 | 2 | 7 | 0 | 0 | 0 | 0 | 0 | 3 |

| Isolate ID | Genus | NIF Grp | *nifH* | *nifD* | *nifK* | *nifE* | *nifN* | *nifB* | *acdS* | *ipdC/ ppdC* | *pqqB* | *pqqC* | *pqqD* | *pqqE* | *pqqF* | *pqq-DH* |
|---|---|---|---|---|---|---|---|---|---|---|---|---|---|---|---|---|
| BCW-200197 | *Kosakonia* | DSP | 1 | 2 | 2 | 3 | 2 | 1 | 2 | 7 | 0 | 0 | 0 | 0 | 0 | 1 |
| BCW-200199 | *unassigned* | DSP | 1 | 2 | 2 | 2 | 3 | 1 | 2 | 7 | 0 | 0 | 0 | 0 | 0 | 3 |
| BCW-200200 | *unassigned* | DSP | 1 | 2 | 2 | 2 | 3 | 1 | 2 | 6 | 0 | 0 | 0 | 0 | 0 | 3 |
| BCW-200201 | *unassigned* | DSP | 1 | 2 | 2 | 2 | 3 | 1 | 2 | 7 | 0 | 0 | 0 | 0 | 0 | 3 |
| BCW-200203 | *Raoultella* | DSP | 2 | 3 | 3 | 5 | 4 | 1 | 2 | 7 | 0 | 0 | 0 | 0 | 0 | 3 |
| BCW-200206 | *Enterobacter* | DSP | 1 | 2 | 2 | 3 | 2 | 1 | 1 | 7 | 0 | 0 | 0 | 0 | 0 | 2 |
| BCW-200210 | *Kosakonia* | DSP | 1 | 2 | 2 | 3 | 2 | 1 | 2 | 7 | 0 | 0 | 0 | 0 | 0 | 1 |
| BCW-200214 | *Kosakonia* | DSP | 1 | 2 | 2 | 3 | 2 | 1 | 2 | 7 | 0 | 0 | 0 | 0 | 0 | 1 |
| BCW-200216 | *Enterobacter* | DSP | 1 | 2 | 2 | 3 | 2 | 1 | 1 | 7 | 0 | 0 | 0 | 0 | 0 | 2 |
| BCW-200221 | *unassigned* | DSP | 1 | 2 | 2 | 2 | 3 | 1 | 2 | 7 | 0 | 0 | 0 | 0 | 0 | 3 |
| BCW-200225 | *unassigned* | DSP | 1 | 2 | 2 | 2 | 3 | 1 | 2 | 7 | 0 | 0 | 0 | 0 | 0 | 3 |
| BCW-200226 | *Kosakonia* | DSP | 1 | 2 | 2 | 3 | 2 | 1 | 2 | 7 | 0 | 0 | 0 | 0 | 0 | 1 |
| BCW-200227 | *Kosakonia* | DSP | 1 | 2 | 2 | 3 | 2 | 1 | 2 | 7 | 0 | 0 | 0 | 0 | 0 | 1 |
| BCW-200234 | *unassigned* | DSP | 1 | 1 | 2 | 1 | 2 | 1 | 2 | 7 | 0 | 0 | 0 | 0 | 0 | 3 |
| BCW-200235 | *unassigned* | DSP | 1 | 2 | 2 | 2 | 3 | 1 | 2 | 6 | 0 | 0 | 0 | 0 | 0 | 3 |
| BCW-200236 | *unassigned* | DSP | 1 | 2 | 2 | 2 | 3 | 1 | 2 | 7 | 0 | 0 | 0 | 0 | 0 | 3 |
| BCW-200237 | *unassigned* | DSP | 1 | 2 | 2 | 2 | 3 | 1 | 2 | 7 | 0 | 0 | 0 | 0 | 0 | 3 |
| BCW-200276 | *Raoultella* | DSP | 1 | 2 | 2 | 3 | 2 | 1 | 2 | 6 | 0 | 0 | 0 | 0 | 0 | 2 |
| BCW-200281 | *Raoultella* | DSP | 2 | 3 | 3 | 5 | 4 | 1 | 2 | 7 | 0 | 0 | 0 | 0 | 0 | 3 |
| BCW-200294 | *Raoultella* | DSP | 2 | 3 | 3 | 5 | 3 | 1 | 2 | 7 | 0 | 0 | 0 | 0 | 0 | 3 |
| BCW-200307 | *Metakosakonia* | DSP | 1 | 2 | 2 | 2 | 2 | 1 | 2 | 5 | 0 | 0 | 0 | 0 | 0 | 2 |
| BCW-200308 | *Metakosakonia* | DSP | 1 | 2 | 2 | 2 | 2 | 1 | 2 | 5 | 0 | 0 | 0 | 0 | 0 | 2 |
| BCW-200317 | *Enterobacter* | DSP | 1 | 2 | 2 | 3 | 2 | 1 | 1 | 7 | 0 | 0 | 0 | 0 | 0 | 2 |
| BCW-200437 | *Raoultella* | DSP | 2 | 3 | 3 | 5 | 4 | 1 | 2 | 7 | 0 | 0 | 0 | 0 | 0 | 3 |
| BCW-200438 | *Raoultella* | DSP | 2 | 4 | 3 | 5 | 3 | 1 | 2 | 7 | 0 | 0 | 0 | 0 | 0 | 3 |
| BCW-200440 | *Raoultella* | DSP | 2 | 3 | 3 | 5 | 4 | 1 | 2 | 7 | 0 | 0 | 0 | 0 | 0 | 3 |

| Isolate ID | Genus | NIF Grp | *nifH* | *nifD* | *nifK* | *nifE* | *nifN* | *nifB* | *acdS* | *ipdC/ ppdC* | *pqqB* | *pqqC* | *pqqD* | *pqqE* | *pqqF* | *pqq-DH* |
|---|---|---|---|---|---|---|---|---|---|---|---|---|---|---|---|---|
| BCW-200442 | *Raoultella* | DSP | 2 | 3 | 3 | 5 | 4 | 1 | 2 | 7 | 0 | 0 | 0 | 0 | 0 | 3 |
| BCW-200444 | *Raoultella* | DSP | 2 | 3 | 3 | 5 | 4 | 1 | 2 | 7 | 0 | 0 | 0 | 0 | 0 | 3 |
| BCW-200446 | *Raoultella* | DSP | 2 | 3 | 3 | 5 | 4 | 1 | 2 | 7 | 0 | 0 | 0 | 0 | 0 | 3 |
| BCW-200449 | *Raoultella* | DSP | 2 | 4 | 3 | 4 | 4 | 1 | 2 | 7 | 0 | 0 | 0 | 0 | 0 | 3 |
| BCW-200488 | *Raoultella* | DSP | 2 | 4 | 3 | 4 | 4 | 1 | 2 | 7 | 0 | 0 | 0 | 0 | 0 | 3 |
| BCW-200496 | *Raoultella* | DSP | 2 | 4 | 3 | 4 | 4 | 1 | 2 | 7 | 0 | 0 | 0 | 0 | 0 | 3 |
| BCW-200499 | *Raoultella* | DSP | 2 | 4 | 3 | 4 | 4 | 1 | 2 | 7 | 0 | 0 | 0 | 0 | 0 | 3 |
| BCW-200509 | *Metakosakonia* | DSP | 1 | 2 | 2 | 2 | 2 | 1 | 2 | 5 | 0 | 0 | 0 | 0 | 0 | 2 |
| BCW-200517 | *Metakosakonia* | DSP | 1 | 2 | 2 | 2 | 2 | 1 | 2 | 5 | 0 | 0 | 0 | 0 | 0 | 2 |
| BCW-200521 | *Raoultella* | DSP | 1 | 2 | 2 | 3 | 3 | 1 | 2 | 7 | 0 | 0 | 0 | 0 | 0 | 3 |
| BCW-200525 | *Raoultella* | DSP | 1 | 2 | 2 | 3 | 3 | 1 | 2 | 7 | 0 | 0 | 0 | 0 | 0 | 3 |
| BCW-200552 | *Rahnella* | DSP | 1 | 2 | 2 | 2 | 2 | 1 | 2 | 6 | 1 | 1 | 1 | 1 | 0 | 2 |
| BCW-200553 | *Raoultella* | DSP | 1 | 2 | 2 | 3 | 2 | 1 | 2 | 6 | 0 | 0 | 0 | 0 | 0 | 2 |
| BCW-200555 | *Raoultella* | DSP | 2 | 4 | 3 | 4 | 4 | 1 | 2 | 7 | 0 | 0 | 0 | 0 | 0 | 3 |
| BCW-200559 | *Rahnella* | DSP | 1 | 2 | 2 | 2 | 2 | 1 | 2 | 6 | 1 | 1 | 1 | 1 | 0 | 2 |
| BCW-200567 | *Metakosakonia* | DSP | 1 | 2 | 2 | 3 | 2 | 1 | 2 | 5 | 0 | 0 | 0 | 0 | 0 | 2 |
| BCW-200577 | *Raoultella* | DSP | 2 | 4 | 3 | 5 | 4 | 1 | 2 | 7 | 0 | 0 | 0 | 0 | 0 | 3 |
| BCW-200578 | *Rahnella* | DSP | 1 | 2 | 2 | 2 | 2 | 1 | 2 | 5 | 1 | 1 | 1 | 1 | 0 | 2 |
| BCW-200600 | *Raoultella* | DSP | 2 | 3 | 3 | 4 | 4 | 1 | 3 | 7 | 0 | 0 | 0 | 0 | 0 | 3 |
| BCW-200620 | *Raoultella* | DSP | 1 | 2 | 2 | 2 | 3 | 1 | 1 | 7 | 0 | 0 | 0 | 0 | 0 | 3 |
| BCW-200644 | *Rahnella* | DSP | 1 | 2 | 2 | 2 | 2 | 1 | 2 | 5 | 1 | 1 | 1 | 1 | 0 | 2 |
| BCW-200647 | *Rahnella* | DSP | 1 | 2 | 2 | 2 | 2 | 1 | 2 | 5 | 1 | 1 | 1 | 1 | 0 | 2 |
| BCW-200648 | *Raoultella* | DSP | 1 | 2 | 2 | 3 | 3 | 1 | 2 | 6 | 1 | 0 | 0 | 0 | 0 | 0 |
| BCW-200650 | *Rahnella* | DSP | 1 | 2 | 2 | 2 | 2 | 1 | 2 | 5 | 1 | 1 | 1 | 1 | 0 | 2 |
| BCW-200651 | *Klebsiella* | DSP | 1 | 2 | 2 | 3 | 3 | 1 | 1 | 7 | 0 | 0 | 0 | 0 | 0 | 3 |
| BCW-200656 | *Metakosakonia* | DSP | 1 | 2 | 2 | 2 | 2 | 1 | 2 | 6 | 0 | 0 | 0 | 0 | 0 | 3 |

| Isolate ID | Genus | NIF Grp | *nifH* | *nifD* | *nifK* | *nifE* | *nifN* | *nifB* | *acdS* | *ipdC/ ppdC* | *pqqB* | *pqqC* | *pqqD* | *pqqE* | *pqqF* | *pqq-DH* |
|---|---|---|---|---|---|---|---|---|---|---|---|---|---|---|---|---|
| BCW-200660 | *Klebsiella* | DSP | 1 | 2 | 2 | 3 | 3 | 1 | 1 | 7 | 0 | 0 | 0 | 0 | 0 | 3 |
| BCW-200661 | *Raoultella* | DSP | 1 | 2 | 2 | 3 | 2 | 1 | 2 | 6 | 0 | 0 | 0 | 0 | 0 | 2 |
| BCW-200662 | *Klebsiella* | DSP | 1 | 2 | 2 | 3 | 3 | 1 | 1 | 7 | 0 | 0 | 0 | 0 | 0 | 3 |
| BCW-200665 | *Raoultella* | DSP | 1 | 2 | 2 | 3 | 2 | 1 | 2 | 6 | 0 | 0 | 0 | 0 | 0 | 2 |
| BCW-200667 | *Enterobacter* | DSP | 1 | 2 | 2 | 3 | 2 | 1 | 1 | 7 | 0 | 0 | 0 | 0 | 0 | 2 |
| BCW-200669 | *Enterobacter* | DSP | 1 | 2 | 2 | 3 | 2 | 1 | 1 | 7 | 0 | 0 | 0 | 0 | 0 | 2 |
| BCW-200704 | *Raoultella* | DSP | 1 | 2 | 2 | 3 | 2 | 1 | 2 | 6 | 0 | 0 | 0 | 0 | 0 | 2 |
| BCW-200727 | *Raoultella* | DSP | 1 | 2 | 2 | 3 | 3 | 1 | 2 | 7 | 0 | 0 | 0 | 0 | 0 | 3 |
| BCW-200738 | *Raoultella* | DSP | 1 | 2 | 2 | 3 | 3 | 1 | 2 | 7 | 0 | 0 | 0 | 0 | 0 | 3 |
| BCW-200776 | *Raoultella* | DSP | 1 | 2 | 2 | 3 | 2 | 1 | 2 | 6 | 0 | 0 | 0 | 0 | 0 | 2 |
| BCW-200785 | *Pseudomonas* | DSP | 2 | 2 | 2 | 3 | 2 | 1 | 2 | 2 | 1 | 1 | 1 | 1 | 0 | 2 |
| BCW-200797 | *Rahnella* | DSP | 1 | 2 | 2 | 2 | 2 | 1 | 2 | 6 | 1 | 1 | 1 | 1 | 0 | 2 |
| BCW-200798 | *Rahnella* | DSP | 1 | 2 | 2 | 2 | 2 | 1 | 2 | 6 | 1 | 1 | 1 | 1 | 0 | 2 |
| BCW-200800 | *Rahnella* | DSP | 1 | 2 | 2 | 2 | 2 | 1 | 2 | 6 | 1 | 1 | 1 | 1 | 0 | 2 |
| BCW-200801 | *Rahnella* | DSP | 1 | 2 | 2 | 2 | 2 | 1 | 2 | 6 | 1 | 1 | 1 | 1 | 0 | 2 |
| BCW-200818 | *Rahnella* | DSP | 1 | 2 | 2 | 2 | 2 | 1 | 2 | 6 | 1 | 1 | 1 | 1 | 0 | 2 |
| BCW-200820 | *Raoultella* | DSP | 2 | 4 | 3 | 4 | 4 | 1 | 2 | 7 | 0 | 0 | 0 | 0 | 0 | 3 |
| BCW-200821 | *Rahnella* | DSP | 1 | 2 | 2 | 2 | 2 | 1 | 2 | 5 | 1 | 1 | 1 | 1 | 0 | 2 |
| BCW-200828 | *Rahnella* | DSP | 1 | 2 | 2 | 2 | 2 | 1 | 2 | 5 | 1 | 1 | 1 | 1 | 0 | 2 |
| BCW-200847 | *Raoultella* | DSP | 2 | 4 | 3 | 4 | 4 | 1 | 2 | 7 | 0 | 0 | 0 | 0 | 0 | 3 |
| BCW-200855 | *Raoultella* | DSP | 2 | 4 | 3 | 4 | 4 | 1 | 2 | 7 | 0 | 0 | 0 | 0 | 0 | 3 |
| BCW-200874 | *Raoultella* | DSP | 2 | 4 | 3 | 4 | 4 | 1 | 2 | 7 | 0 | 0 | 0 | 0 | 0 | 3 |
| BCW-200879 | *Raoultella* | DSP | 2 | 4 | 3 | 4 | 4 | 1 | 2 | 7 | 0 | 0 | 1 | 0 | 0 | 3 |
| BCW-200880 | *Raoultella* | DSP | 2 | 4 | 3 | 4 | 4 | 1 | 2 | 7 | 0 | 0 | 0 | 0 | 0 | 3 |
| BCW-200881 | *Raoultella* | DSP | 2 | 4 | 3 | 4 | 4 | 1 | 2 | 7 | 0 | 0 | 0 | 0 | 0 | 3 |
| BCW-200882 | *Raoultella* | DSP | 2 | 4 | 3 | 4 | 4 | 1 | 2 | 7 | 0 | 0 | 0 | 0 | 0 | 3 |

| Isolate ID | Genus | NIF Grp | *nifH* | *nifD* | *nifK* | *nifE* | *nifN* | *nifB* | *acdS* | *ipdC/ ppdC* | *pqqB* | *pqqC* | *pqqD* | *pqqE* | *pqqF* | *pqq-DH* |
|---|---|---|---|---|---|---|---|---|---|---|---|---|---|---|---|---|
| BCW-200885 | *Raoultella* | DSP | 2 | 4 | 3 | 4 | 4 | 1 | 2 | 7 | 0 | 0 | 0 | 0 | 0 | 3 |
| BCW-200886 | *Raoultella* | DSP | 2 | 4 | 3 | 4 | 4 | 1 | 2 | 7 | 0 | 0 | 0 | 0 | 0 | 3 |
| BCW-200887 | *Raoultella* | DSP | 2 | 4 | 3 | 4 | 4 | 1 | 2 | 7 | 0 | 0 | 0 | 0 | 0 | 3 |
| BCW-200891 | *Raoultella* | DSP | 2 | 3 | 3 | 5 | 4 | 1 | 2 | 7 | 0 | 0 | 0 | 0 | 0 | 3 |
| BCW-200892 | *Raoultella* | DSP | 2 | 3 | 3 | 5 | 4 | 1 | 2 | 7 | 0 | 0 | 0 | 0 | 0 | 3 |
| BCW-200909 | *Raoultella* | DSP | 2 | 4 | 3 | 4 | 4 | 1 | 2 | 7 | 0 | 0 | 0 | 0 | 0 | 3 |
| BCW-200915 | *Raoultella* | DSP | 2 | 4 | 3 | 4 | 4 | 1 | 2 | 7 | 0 | 0 | 0 | 0 | 0 | 3 |
| BCW-200926 | *Raoultella* | DSP | 2 | 4 | 3 | 4 | 4 | 1 | 2 | 7 | 0 | 0 | 0 | 0 | 0 | 3 |
| BCW-200951 | *Metakosakonia* | DSP | 1 | 2 | 2 | 3 | 2 | 1 | 2 | 5 | 0 | 0 | 0 | 0 | 0 | 2 |
| BCW-200955 | *Metakosakonia* | DSP | 1 | 2 | 2 | 3 | 2 | 1 | 2 | 5 | 0 | 0 | 0 | 0 | 0 | 2 |
| BCW-201019 | *Klebsiella* | DSP | 1 | 2 | 2 | 3 | 2 | 1 | 2 | 6 | 0 | 0 | 0 | 0 | 0 | 3 |
| BCW-201020 | *Klebsiella* | DSP | 1 | 2 | 2 | 3 | 2 | 1 | 2 | 6 | 0 | 0 | 0 | 0 | 0 | 3 |
| BCW-201021 | *Klebsiella* | DSP | 1 | 2 | 2 | 3 | 2 | 1 | 2 | 6 | 0 | 0 | 0 | 0 | 0 | 3 |
| BCW-201044 | *Raoultella* | DSP | 2 | 4 | 3 | 4 | 4 | 1 | 2 | 7 | 0 | 0 | 0 | 0 | 0 | 3 |
| BCW-201058 | *Metakosakonia* | DSP | 2 | 4 | 3 | 4 | 3 | 1 | 2 | 5 | 0 | 0 | 0 | 0 | 0 | 3 |
| BCW-201059 | *Rahnella* | DSP | 1 | 2 | 2 | 2 | 2 | 1 | 2 | 6 | 1 | 1 | 1 | 1 | 0 | 2 |
| BCW-201070 | *Rahnella* | DSP | 1 | 2 | 2 | 2 | 2 | 1 | 2 | 6 | 1 | 1 | 1 | 1 | 0 | 2 |
| BCW-201071 | *Raoultella* | DSP | 1 | 2 | 2 | 3 | 3 | 1 | 2 | 7 | 0 | 0 | 0 | 0 | 0 | 3 |
| BCW-201075 | *Raoultella* | DSP | 1 | 2 | 2 | 3 | 2 | 1 | 2 | 6 | 0 | 0 | 0 | 0 | 0 | 2 |
| BCW-201078 | *Raoultella* | DSP | 1 | 2 | 2 | 3 | 2 | 1 | 2 | 6 | 0 | 0 | 0 | 0 | 0 | 2 |
| BCW-201088 | *Rahnella* | DSP | 1 | 2 | 2 | 2 | 2 | 1 | 2 | 6 | 1 | 1 | 1 | 1 | 0 | 2 |
| BCW-201091 | *Rahnella* | DSP | 1 | 2 | 2 | 2 | 2 | 1 | 2 | 6 | 1 | 1 | 1 | 1 | 0 | 2 |
| BCW-201097 | *unassigned* | DSP | 1 | 2 | 2 | 3 | 2 | 1 | 2 | 6 | 0 | 0 | 0 | 0 | 0 | 2 |
| BCW-201098 | *Raoultella* | DSP | 1 | 2 | 2 | 3 | 2 | 1 | 2 | 6 | 0 | 0 | 0 | 0 | 0 | 2 |
| BCW-201107 | *Raoultella* | DSP | 1 | 2 | 2 | 3 | 2 | 1 | 2 | 5 | 1 | 1 | 0 | 0 | 0 | 2 |
| BCW-201155 | *Metakosakonia* | DSP | 2 | 4 | 3 | 4 | 3 | 1 | 2 | 5 | 0 | 0 | 0 | 0 | 0 | 3 |

| Isolate ID | Genus | NIF Grp | *nifH* | *nifD* | *nifK* | *nifE* | *nifN* | *nifB* | *acdS* | *ipdC/ ppdC* | *pqqB* | *pqqC* | *pqqD* | *pqqE* | *pqqF* | *pqq-DH* |
|---|---|---|---|---|---|---|---|---|---|---|---|---|---|---|---|---|
| BCW-201162 | *Rahnella* | DSP | 1 | 2 | 2 | 2 | 2 | 1 | 2 | 6 | 1 | 1 | 1 | 1 | 0 | 2 |
| BCW-201184 | *Rahnella* | DSP | 1 | 2 | 2 | 2 | 2 | 1 | 2 | 6 | 1 | 1 | 1 | 1 | 0 | 2 |
| BCW-201259 | *Rahnella* | DSP | 1 | 2 | 2 | 2 | 2 | 1 | 2 | 6 | 1 | 1 | 1 | 1 | 0 | 2 |
| BCW-201263 | *Klebsiella* | DSP | 1 | 2 | 2 | 3 | 3 | 1 | 1 | 7 | 0 | 0 | 0 | 0 | 0 | 3 |
| BCW-201267 | *unassigned* | DSP | 2 | 3 | 3 | 5 | 4 | 1 | 2 | 7 | 0 | 0 | 0 | 0 | 0 | 3 |
| BCW-201290 | *Pseudomonas* | DSP | 2 | 2 | 2 | 3 | 2 | 1 | 2 | 2 | 1 | 1 | 1 | 1 | 0 | 2 |
| BCW-201297 | *Rahnella* | DSP | 1 | 2 | 2 | 2 | 2 | 1 | 2 | 5 | 1 | 1 | 1 | 1 | 0 | 2 |
| BCW-201302 | *Rahnella* | DSP | 1 | 2 | 2 | 2 | 2 | 1 | 2 | 5 | 1 | 1 | 1 | 1 | 0 | 2 |
| BCW-201304 | *Rahnella* | DSP | 1 | 2 | 2 | 2 | 2 | 1 | 2 | 5 | 1 | 1 | 1 | 1 | 0 | 2 |
| BCW-201315 | *Rahnella* | DSP | 1 | 2 | 2 | 2 | 2 | 1 | 2 | 5 | 1 | 1 | 1 | 1 | 0 | 2 |
| BCW-201441 | *Klebsiella* | DSP | 1 | 2 | 2 | 3 | 3 | 1 | 1 | 7 | 0 | 0 | 0 | 0 | 0 | 3 |
| BCW-201443 | *Raoultella* | DSP | 1 | 2 | 2 | 3 | 3 | 1 | 2 | 7 | 0 | 0 | 0 | 0 | 0 | 3 |
| BCW-201461 | *Raoultella* | DSP | 1 | 2 | 2 | 3 | 3 | 1 | 2 | 7 | 0 | 0 | 0 | 0 | 0 | 3 |
| BCW-201614 | *Raoultella* | DSP | 1 | 2 | 2 | 3 | 2 | 1 | 2 | 6 | 0 | 0 | 0 | 0 | 0 | 2 |
| BCW-201615 | *Raoultella* | DSP | 1 | 2 | 2 | 3 | 2 | 1 | 2 | 6 | 0 | 0 | 0 | 0 | 0 | 2 |
| BCW-201620 | *Raoultella* | DSP | 1 | 2 | 2 | 3 | 2 | 1 | 2 | 6 | 0 | 0 | 0 | 0 | 0 | 2 |
| BCW-201649 | *Raoultella* | DSP | 2 | 4 | 3 | 5 | 3 | 1 | 2 | 7 | 0 | 0 | 0 | 0 | 0 | 3 |
| BCW-201659 | *Enterobacter* | DSP | 1 | 2 | 2 | 3 | 2 | 1 | 1 | 7 | 0 | 0 | 0 | 0 | 0 | 2 |
| BCW-201703 | *Raoultella* | DSP | 1 | 2 | 2 | 3 | 2 | 1 | 2 | 6 | 0 | 0 | 0 | 0 | 0 | 2 |
| BCW-201721 | *Enterobacter* | DSP | 1 | 2 | 2 | 3 | 2 | 1 | 1 | 7 | 0 | 0 | 0 | 0 | 0 | 2 |
| BCW-201808 | *Metakosakonia* | DSP | 1 | 2 | 2 | 2 | 3 | 1 | 2 | 6 | 0 | 0 | 0 | 0 | 0 | 3 |
| BCW-201828 | *Metakosakonia* | DSP | 1 | 2 | 2 | 2 | 3 | 1 | 2 | 6 | 0 | 0 | 0 | 0 | 0 | 3 |
| BCW-201850 | *Metakosakonia* | DSP | 1 | 2 | 2 | 2 | 3 | 1 | 2 | 6 | 0 | 0 | 0 | 0 | 0 | 3 |
| BCW-201858 | *Metakosakonia* | DSP | 1 | 2 | 2 | 2 | 3 | 1 | 2 | 6 | 0 | 0 | 0 | 0 | 0 | 3 |
| BCW-201873 | *Metakosakonia* | DSP | 1 | 2 | 2 | 2 | 3 | 1 | 2 | 6 | 0 | 0 | 0 | 0 | 0 | 3 |
| BCW-201876 | *Metakosakonia* | DSP | 1 | 2 | 2 | 2 | 3 | 1 | 2 | 6 | 0 | 0 | 0 | 0 | 0 | 3 |

| Isolate ID | Genus | NIF Grp | *nifH* | *nifD* | *nifK* | *nifE* | *nifN* | *nifB* | *acdS* | *ipdC/ ppdC* | *pqqB* | *pqqC* | *pqqD* | *pqqE* | *pqqF* | *pqq-DH* |
|---|---|---|---|---|---|---|---|---|---|---|---|---|---|---|---|---|
| BCW-201879 | *Metakosakonia* | DSP | 1 | 2 | 2 | 2 | 3 | 1 | 2 | 6 | 0 | 0 | 0 | 0 | 0 | 3 |
| BCW-201886 | *Metakosakonia* | DSP | 1 | 2 | 2 | 2 | 3 | 1 | 2 | 6 | 0 | 0 | 0 | 0 | 0 | 3 |
| BCW-201887 | *Metakosakonia* | DSP | 1 | 2 | 2 | 2 | 3 | 1 | 2 | 6 | 0 | 0 | 0 | 0 | 0 | 3 |
| BCW-201888 | *Metakosakonia* | DSP | 1 | 2 | 2 | 2 | 3 | 1 | 2 | 6 | 0 | 0 | 0 | 0 | 0 | 3 |
| BCW-201890 | *Raoultella* | DSP | 1 | 2 | 2 | 3 | 3 | 1 | 2 | 7 | 0 | 0 | 0 | 0 | 0 | 3 |
| BCW-201900 | *Raoultella* | DSP | 2 | 3 | 3 | 4 | 4 | 1 | 3 | 7 | 0 | 0 | 0 | 0 | 0 | 3 |
| BCW-201901 | *Raoultella* | DSP | 2 | 4 | 3 | 4 | 4 | 1 | 2 | 7 | 0 | 0 | 0 | 0 | 0 | 3 |
| BCW-201926 | *Raoultella* | DSP | 2 | 4 | 3 | 5 | 4 | 1 | 2 | 7 | 0 | 0 | 0 | 0 | 0 | 3 |
| BCW-201937 | *Metakosakonia* | DSP | 1 | 2 | 2 | 2 | 3 | 1 | 2 | 6 | 0 | 0 | 0 | 0 | 0 | 3 |
| BCW-201972 | *Metakosakonia* | DSP | 1 | 2 | 2 | 2 | 3 | 1 | 2 | 6 | 0 | 0 | 0 | 0 | 0 | 3 |
| BCW-201990 | *Metakosakonia* | DSP | 1 | 2 | 2 | 2 | 3 | 1 | 2 | 6 | 0 | 0 | 0 | 0 | 0 | 3 |
| BCW-200001 | *Acidovorax* | SDS | 2 | 0 | 0 | 0 | 0 | 0 | 1 | 3 | 1 | 1 | 1 | 1 | 0 | 2 |
| BCW-200002 | *unassigned* | SDS | 1 | 0 | 0 | 0 | 0 | 0 | 0 | 3 | 1 | 1 | 1 | 1 | 0 | 2 |
| BCW-200008 | *Acinetobacter* | SDS | 1 | 0 | 0 | 0 | 0 | 0 | 1 | 7 | 0 | 0 | 0 | 0 | 0 | 2 |
| BCW-200011 | *Acidovorax* | SDS | 2 | 0 | 0 | 0 | 0 | 0 | 1 | 2 | 1 | 1 | 2 | 1 | 0 | 2 |
| BCW-200032 | *Acinetobacter* | SDS | 1 | 0 | 0 | 0 | 0 | 0 | 0 | 3 | 1 | 1 | 1 | 1 | 0 | 2 |
| BCW-200044 | *Acinetobacter* | SDS | 1 | 0 | 0 | 0 | 0 | 0 | 0 | 3 | 1 | 1 | 1 | 1 | 0 | 2 |
| BCW-200046 | *Pseudomonas* | SDS | 1 | 0 | 0 | 0 | 0 | 0 | 1 | 4 | 2 | 1 | 1 | 2 | 0 | 2 |
| BCW-200056 | *Pseudomonas* | SDS | 1 | 0 | 0 | 0 | 0 | 0 | 1 | 4 | 1 | 1 | 1 | 1 | 1 | 2 |
| BCW-200065 | *Pseudomonas* | SDS | 1 | 0 | 0 | 0 | 0 | 0 | 1 | 3 | 1 | 1 | 1 | 1 | 1 | 3 |
| BCW-200068 | *Pseudomonas* | SDS | 2 | 0 | 0 | 0 | 0 | 0 | 2 | 3 | 1 | 1 | 1 | 1 | 1 | 2 |
| BCW-200073 | *Pseudomonas* | SDS | 1 | 0 | 0 | 0 | 0 | 0 | 1 | 2 | 1 | 1 | 1 | 3 | 1 | 2 |
| BCW-200078 | *Micrococcus* | SDS | 1 | 0 | 0 | 0 | 0 | 0 | 0 | 2 | 0 | 0 | 0 | 0 | 0 | 0 |
| BCW-200081 | *Acinetobacter* | SDS | 1 | 0 | 0 | 0 | 0 | 0 | 0 | 3 | 1 | 1 | 1 | 1 | 0 | 2 |
| BCW-200100 | *Acinetobacter* | SDS | 1 | 0 | 0 | 0 | 0 | 0 | 0 | 3 | 1 | 1 | 1 | 1 | 0 | 2 |
| BCW-200101 | *Bacillus* | SDS | 2 | 0 | 0 | 0 | 0 | 0 | 0 | 2 | 0 | 0 | 0 | 0 | 0 | 0 |

| Isolate ID | Genus | NIF Grp | *nifH* | *nifD* | *nifK* | *nifE* | *nifN* | *nifB* | *acdS* | *ipdC/ ppdC* | *pqqB* | *pqqC* | *pqqD* | *pqqE* | *pqqF* | *pqq-DH* |
|---|---|---|---|---|---|---|---|---|---|---|---|---|---|---|---|---|
| BCW-200103 | *Stenotrophomonas* | SDS | 1 | 0 | 0 | 0 | 0 | 0 | 0 | 2 | 0 | 0 | 0 | 0 | 0 | 0 |
| BCW-200105 | *Acinetobacter* | SDS | 1 | 0 | 0 | 0 | 0 | 0 | 0 | 3 | 1 | 1 | 1 | 1 | 0 | 2 |
| BCW-200112 | *Acinetobacter* | SDS | 1 | 0 | 0 | 0 | 0 | 0 | 0 | 3 | 1 | 1 | 1 | 1 | 0 | 2 |
| BCW-200116 | *Acinetobacter* | SDS | 1 | 0 | 0 | 0 | 0 | 0 | 0 | 3 | 1 | 1 | 1 | 1 | 0 | 2 |
| BCW-200118 | *Acinetobacter* | SDS | 1 | 0 | 0 | 0 | 0 | 0 | 0 | 3 | 1 | 1 | 1 | 1 | 0 | 2 |
| BCW-200119 | *Acinetobacter* | SDS | 1 | 0 | 0 | 0 | 0 | 0 | 0 | 3 | 1 | 1 | 1 | 1 | 0 | 2 |
| BCW-200147 | *unassigned* | SDS | 1 | 2 | 2 | 2 | 3 | 0 | 2 | 7 | 0 | 0 | 0 | 0 | 0 | 3 |
| BCW-200153 | *Stenotrophomonas* | SDS | 1 | 0 | 0 | 0 | 0 | 0 | 0 | 3 | 0 | 0 | 0 | 0 | 0 | 1 |
| BCW-200154 | *Stenotrophomonas* | SDS | 1 | 0 | 0 | 0 | 0 | 0 | 0 | 3 | 0 | 0 | 0 | 0 | 0 | 1 |
| BCW-200173 | *Micrococcus* | SDS | 1 | 0 | 0 | 0 | 0 | 0 | 0 | 3 | 0 | 0 | 0 | 0 | 0 | 0 |
| BCW-200202 | *Stenotrophomonas* | SDS | 1 | 0 | 0 | 0 | 0 | 0 | 0 | 1 | 0 | 0 | 0 | 1 | 0 | 1 |
| BCW-200209 | *Leifsonia* | SDS | 1 | 0 | 0 | 0 | 0 | 0 | 0 | 1 | 0 | 0 | 0 | 0 | 0 | 0 |
| BCW-200266 | *Stenotrophomonas* | SDS | 1 | 0 | 0 | 0 | 0 | 0 | 0 | 3 | 0 | 0 | 0 | 0 | 0 | 1 |
| BCW-200267 | *Pseudomonas* | SDS | 2 | 0 | 0 | 0 | 0 | 0 | 2 | 4 | 1 | 1 | 1 | 1 | 1 | 2 |
| BCW-200290 | *Pseudomonas* | SDS | 1 | 0 | 0 | 0 | 0 | 0 | 1 | 4 | 1 | 1 | 1 | 1 | 1 | 2 |
| BCW-200432 | *Pseudomonas* | SDS | 1 | 0 | 0 | 0 | 0 | 0 | 1 | 4 | 1 | 1 | 1 | 1 | 1 | 2 |
| BCW-200436 | *Pseudomonas* | SDS | 1 | 0 | 0 | 0 | 0 | 0 | 1 | 3 | 1 | 1 | 1 | 1 | 1 | 2 |
| BCW-200443 | *Pseudomonas* | SDS | 1 | 0 | 0 | 0 | 0 | 0 | 1 | 4 | 1 | 1 | 1 | 1 | 1 | 2 |
| BCW-200458 | *Pseudomonas* | SDS | 1 | 0 | 0 | 0 | 0 | 0 | 1 | 3 | 1 | 1 | 1 | 1 | 1 | 2 |
| BCW-200460 | *Pseudomonas* | SDS | 1 | 0 | 0 | 0 | 0 | 0 | 1 | 3 | 1 | 1 | 1 | 2 | 1 | 3 |
| BCW-200476 | *Pseudomonas* | SDS | 2 | 0 | 0 | 0 | 0 | 0 | 2 | 3 | 1 | 1 | 1 | 1 | 1 | 2 |
| BCW-200477 | *Pseudomonas* | SDS | 1 | 0 | 0 | 0 | 0 | 0 | 0 | 7 | 1 | 1 | 1 | 1 | 1 | 2 |
| BCW-200497 | *Pseudomonas* | SDS | 2 | 0 | 0 | 0 | 0 | 0 | 2 | 4 | 1 | 1 | 1 | 1 | 1 | 2 |
| BCW-200498 | *unassigned* | SDS | 2 | 0 | 0 | 0 | 0 | 0 | 2 | 4 | 1 | 1 | 1 | 1 | 1 | 2 |
| BCW-200527 | *Pseudomonas* | SDS | 2 | 0 | 0 | 0 | 0 | 0 | 2 | 3 | 1 | 1 | 1 | 1 | 1 | 2 |
| BCW-200528 | *Pseudomonas* | SDS | 2 | 0 | 0 | 0 | 0 | 0 | 2 | 3 | 1 | 1 | 1 | 1 | 1 | 2 |

| Isolate ID | Genus | NIF Grp | *nifH* | *nifD* | *nifK* | *nifE* | *nifN* | *nifB* | *acdS* | *ipdC/ ppdC* | *pqqB* | *pqqC* | *pqqD* | *pqqE* | *pqqF* | *pqq-DH* |
|---|---|---|---|---|---|---|---|---|---|---|---|---|---|---|---|---|
| BCW-200570 | *Stenotrophomonas* | SDS | 1 | 0 | 0 | 0 | 0 | 0 | 0 | 3 | 0 | 0 | 0 | 0 | 0 | 1 |
| BCW-200574 | *Stenotrophomonas* | SDS | 1 | 0 | 0 | 0 | 0 | 0 | 0 | 3 | 0 | 0 | 0 | 0 | 0 | 1 |
| BCW-200587 | *Stenotrophomonas* | SDS | 1 | 0 | 0 | 0 | 0 | 0 | 0 | 3 | 0 | 0 | 0 | 0 | 0 | 1 |
| BCW-200588 | *Pseudomonas* | SDS | 2 | 0 | 0 | 0 | 0 | 0 | 2 | 4 | 1 | 1 | 1 | 1 | 1 | 2 |
| BCW-200589 | *Stenotrophomonas* | SDS | 1 | 0 | 0 | 0 | 0 | 0 | 0 | 3 | 0 | 0 | 0 | 0 | 0 | 1 |
| BCW-200599 | *Pseudomonas* | SDS | 1 | 0 | 0 | 0 | 0 | 0 | 1 | 3 | 1 | 1 | 1 | 1 | 1 | 2 |
| BCW-200607 | *Pseudomonas* | SDS | 2 | 0 | 0 | 0 | 0 | 0 | 2 | 4 | 1 | 1 | 1 | 1 | 1 | 2 |
| BCW-200621 | *Pseudomonas* | SDS | 1 | 0 | 0 | 0 | 0 | 0 | 1 | 3 | 1 | 1 | 1 | 1 | 1 | 2 |
| BCW-200646 | *Stenotrophomonas* | SDS | 1 | 0 | 0 | 0 | 0 | 0 | 0 | 3 | 0 | 0 | 0 | 0 | 0 | 1 |
| BCW-200790 | *Pseudomonas* | SDS | 2 | 0 | 0 | 0 | 0 | 0 | 2 | 3 | 1 | 1 | 1 | 1 | 1 | 2 |
| BCW-200791 | *Pseudomonas* | SDS | 2 | 0 | 0 | 0 | 0 | 0 | 2 | 3 | 1 | 1 | 1 | 1 | 1 | 2 |
| BCW-200938 | *Stenotrophomonas* | SDS | 1 | 0 | 0 | 0 | 0 | 0 | 0 | 3 | 0 | 0 | 0 | 0 | 0 | 1 |
| BCW-200939 | *Stenotrophomonas* | SDS | 1 | 0 | 0 | 0 | 0 | 0 | 0 | 3 | 0 | 0 | 0 | 0 | 0 | 1 |
| BCW-200944 | *Stenotrophomonas* | SDS | 1 | 0 | 0 | 0 | 0 | 0 | 0 | 3 | 0 | 0 | 0 | 0 | 0 | 1 |
| BCW-200945 | *Stenotrophomonas* | SDS | 1 | 0 | 0 | 0 | 0 | 0 | 0 | 3 | 0 | 0 | 0 | 0 | 0 | 1 |
| BCW-200993 | *Herbaspirillum* | SDS | 2 | 0 | 0 | 0 | 0 | 0 | 1 | 4 | 0 | 0 | 0 | 0 | 0 | 0 |
| BCW-201292 | *Pseudomonas* | SDS | 1 | 0 | 0 | 0 | 0 | 0 | 1 | 3 | 1 | 1 | 1 | 1 | 1 | 2 |
| BCW-201293 | *Pseudomonas* | SDS | 1 | 0 | 0 | 0 | 0 | 0 | 1 | 3 | 1 | 1 | 1 | 1 | 1 | 2 |
| BCW-201819 | *Curtobacterium* | SDS | 2 | 0 | 0 | 0 | 0 | 0 | 0 | 3 | 0 | 0 | 0 | 0 | 0 | 0 |
| BCW-201851 | *Stenotrophomonas* | SDS | 1 | 0 | 0 | 0 | 0 | 0 | 0 | 2 | 0 | 0 | 0 | 0 | 0 | 1 |
| BCW-201859 | *Pseudomonas* | SDS | 1 | 0 | 0 | 0 | 0 | 0 | 0 | 6 | 1 | 1 | 2 | 1 | 1 | 2 |
| BCW-201862 | *Pseudomonas* | SDS | 1 | 0 | 0 | 0 | 0 | 0 | 1 | 3 | 1 | 1 | 1 | 1 | 1 | 2 |
| BCW-201868 | *Pseudomonas* | SDS | 1 | 0 | 0 | 0 | 0 | 0 | 0 | 6 | 1 | 1 | 2 | 1 | 1 | 3 |
| BCW-201875 | *Pseudomonas* | SDS | 1 | 0 | 0 | 0 | 0 | 0 | 0 | 5 | 1 | 1 | 2 | 1 | 1 | 2 |
| BCW-201947 | *Pseudomonas* | SDS | 1 | 0 | 0 | 0 | 0 | 0 | 0 | 5 | 1 | 1 | 2 | 1 | 1 | 2 |

Numbers represent counts for homologous sequences matching HMMs of marker genes for targeted PGP traits in each isolate's whole genome sequence assembly. The number of significant matches to TIGRFAM [1] HMMs for marker genes of *nif* and non-*nif* PGP traits by predicted coding sequences identified in each isolate's whole genome sequence were counted using functions from base and tidyverse 1.2.1 packages in R [3]. Predicted coding sequences were counted as significant homologous genes to HMMs if the threshold of a maximum e-value of $1e_{-06}$ and HMM coverage of 80% were met. NIF Group (NIF Grp) assignments for each isolate were annotated in the table alongside HMM search results and include Dos Santos Positive (DSP), Semi-Dos Santos (SDS) and Dos Santos Negative (DSN). Genus information for each isolate was determined using Sourmash [2] LCA classification with the GTDB v89 [4] and GenBank [5] databases with a k-size of 31. PGP trait profiles included possession of essential marker genes for the BNF trait proposed by the Dos Santos model (*nifH*, *nifD*, *nifK*, *nifE*, *nifN*, *nifB*), the marker gene for deamination of ACC (*acdS*), the marker gene for biosynthesis of IAA (*ipdC/ppdC*), and marker genes for phosphate solubilization using the PQQ mechanism mediated by the *pqqB*, *pqqC*, *pqqD*, *pqqE*, *pqqF* and PQQ Dehydrogenase (*pqq-DH*) genes. Data for homologous sequence counts of BNF marker genes were previously generated and reported [6].

### S2 Table. Summary of PGP assay values for each isolate

| Isolate ID | Genus | Group | ACC (RGR) | BNF (15N/14N) | IAA (mg/mL) | PO4 (mg/L) |
|---|---|---|---|---|---|---|
| BCW-200097 | *Acinetobacter* | DSN | 1.22 | 1 | 0.8 | 415.13 |
| BCW-200663 | *Agrobacterium* | DSN | 1.12 | 1 | 3.59 | 1182.06 |
| BCW-200902 | *Agrobacterium* | DSN | 1.26 | 1.5 | 11.55 | 998.98 |
| BCW-200910 | *Agrobacterium* | DSN | 1.84 | 1 | 8.35 | 911.05 |
| BCW-200465 | *Agrobacterium* | DSN | 2.54 | 1 | 20.36 | 911.05 |
| BCW-200920 | *Agrobacterium* | DSN | 2.19 | 1.1 | 10.05 | 777.1 |
| BCW-200904 | *Agrobacterium* | DSN | 1.9 | 1.2 | 15.5 | 760.74 |
| BCW-200328 | *Agrobacterium* | DSN | 2.51 | 1.5 | 19.82 | 721.88 |
| BCW-200464 | *Agrobacterium* | DSN | 1.91 | 1 | 24.45 | 413.41 |
| BCW-200208 | *Agrobacterium* | DSN | 1.67 | 1.2 | 12.73 | 320.24 |
| BCW-200705 | *Agrobacterium* | DSN | 1.13 | 1.2 | 1.86 | 238.06 |
| BCW-200215 | *Agrobacterium* | DSN | 0.97 | 1 | 14.5 | 136.8 |
| BCW-201445 | *Atlantibacter* | DSN | 1.11 | 1 | 19.08 | 879.35 |
| BCW-201444 | *Atlantibacter* | DSN | 2.1 | 1 | 17 | 835.38 |
| BCW-200066 | *Atlantibacter* | DSN | 1.04 | 1 | 37.63 | 822.09 |
| BCW-200986 | *Citrobacter* | DSN | 1.89 | 1.1 | 8.83 | 1503.07 |
| BCW-201083 | *Citrobacter* | DSN | 1.22 | 1.3 | 6.08 | 1194.27 |
| BCW-200111 | *Citrobacter* | DSN | 1.39 | 1.2 | 48.41 | 1171.92 |
| BCW-200031 | *Citrobacter* | DSN | 1.03 | 1 | -0.11 | 1062.37 |
| BCW-200539 | *Citrobacter* | DSN | 1.22 | 1.2 | 3.82 | 1049.08 |
| BCW-201154 | *Citrobacter* | DSN | 2.22 | 1 | 3.23 | 1020.45 |
| BCW-201152 | *Citrobacter* | DSN | 1.42 | 1.3 | 1.86 | 926.38 |
| BCW-200984 | *Citrobacter* | DSN | 1.76 | 1 | 13.49 | 898.77 |
| BCW-200036 | *Citrobacter* | DSN | 2.59 | 1 | 3.31 | 882.41 |
| BCW-200983 | *Citrobacter* | DSN | 2.24 | 1.1 | 2.26 | 790.39 |
| BCW-201842 | *Curtobacterium* | DSN | 1.43 | 1 | 29.69 | 511.61 |
| BCW-201896 | *Curtobacterium* | DSN | 2.45 | 1.1 | 0.74 | 151.51 |
| BCW-200219 | *Enterobacter* | DSN | 1.18 | 0.9 | 7.71 | 1339.9 |
| BCW-200104 | *Enterobacter* | DSN | 1.17 | No Data | 7.09 | 1173.23 |
| BCW-201849 | *Enterobacter* | DSN | 1.87 | 1.3 | 5.93 | 1151.33 |
| BCW-200218 | *Enterobacter* | DSN | 1.16 | 0.9 | 12.12 | 1118.11 |
| BCW-200107 | *Enterobacter* | DSN | 1.14 | 1.1 | 15.65 | 1057.74 |
| BCW-201938 | *Enterobacter* | DSN | 0.86 | 1.1 | 4.17 | 1051.12 |
| BCW-201957 | *Enterobacter* | DSN | 0.96 | 1.1 | 19.59 | 1048.06 |
| BCW-200054 | *Enterobacter* | DSN | 1.02 | 1 | 20.67 | 1015.34 |
| BCW-201839 | *Enterobacter* | DSN | 2.73 | 1.1 | 97.53 | 1012.27 |
| BCW-200012 | *Enterobacter* | DSN | 1.15 | 1 | 15.53 | 1002.04 |
| BCW-201903 | *Enterobacter* | DSN | 0.93 | 1 | 17.48 | 965.24 |
| BCW-201945 | *Enterobacter* | DSN | 1.16 | 1.1 | 14.43 | 958.08 |
| BCW-200009 | *Enterobacter* | DSN | 1.27 | 1 | 19.23 | 949.9 |
| BCW-200025 | *Enterobacter* | DSN | 0.86 | 1 | 5.9 | 939.67 |
| BCW-201811 | *Enterobacter* | DSN | 2.01 | 1 | 20.41 | 931.49 |
| BCW-201914 | *Enterobacter* | DSN | 0.76 | 1 | 7.38 | 921.27 |
| BCW-201878 | *Enterobacter* | DSN | 1.1 | 1.4 | 18.93 | 911.04 |
| BCW-200047 | *Enterobacter* | DSN | 1.08 | 0.9 | 29.6 | 891.62 |
| BCW-200023 | *Enterobacter* | DSN | 1.06 | 0.8 | 7.42 | 860.94 |
| BCW-201877 | *Enterobacter* | DSN | 0.96 | 1.2 | 43.21 | 858.9 |
| BCW-201882 | *Enterobacter* | DSN | 1.06 | 1.4 | 8.55 | 857.87 |
| BCW-201885 | *Enterobacter* | DSN | 2.3 | 1.4 | 15.04 | 854.81 |
| BCW-200013 | *Enterobacter* | DSN | 1.1 | 1 | 6.07 | 852.76 |
| BCW-201881 | *Enterobacter* | DSN | 1.04 | 1.2 | 11.53 | 850.72 |
| BCW-201857 | *Enterobacter* | DSN | 0.92 | 1 | 9.92 | 846.84 |

| Isolate ID | Genus | Group | ACC (RGR) | BNF (15N/14N) | IAA (mg/mL) | PO4 (mg/L) |
|---|---|---|---|---|---|---|
| BCW-201933 | *Enterobacter* | DSN | 1.1 | 1.4 | 14.4 | 846.63 |
| BCW-201866 | *Enterobacter* | DSN | 1.53 | 1.4 | 16.77 | 845.6 |
| BCW-200055 | *Enterobacter* | DSN | 0.98 | 0.9 | 26.52 | 843.56 |
| BCW-201814 | *Enterobacter* | DSN | 1.33 | 1.2 | 11.91 | 835.38 |
| BCW-200029 | *Enterobacter* | DSN | 1.04 | 1 | 3.76 | 834.36 |
| BCW-200268 | *Enterobacter* | DSN | 2.3 | 1 | 15.17 | 832.67 |
| BCW-200027 | *Enterobacter* | DSN | 0.9 | 1 | 3.84 | 832.31 |
| BCW-200018 | *Enterobacter* | DSN | 1.01 | 1 | 10.51 | 826.18 |
| BCW-201880 | *Enterobacter* | DSN | 1.5 | 2.2 | 26.54 | 823.11 |
| BCW-201726 | *Enterobacter* | DSN | 2.69 | 1.1 | 8.7 | 810.84 |
| BCW-200092 | *Enterobacter* | DSN | 2.11 | 1 | 22.36 | 808.79 |
| BCW-201895 | *Enterobacter* | DSN | 1.09 | 1.7 | 18.55 | 808.79 |
| BCW-200043 | *Enterobacter* | DSN | 1.08 | 1.1 | 0.76 | 804.7 |
| BCW-200041 | *Enterobacter* | DSN | 2.51 | 1 | 34.91 | 790.39 |
| BCW-200064 | *Enterobacter* | DSN | 1.01 | 1 | 21.54 | 790.39 |
| BCW-200279 | *Enterobacter* | DSN | 2.33 | 1.3 | 23.18 | 786.3 |
| BCW-201874 | *Enterobacter* | DSN | 1.01 | 1.3 | 10.15 | 783.23 |
| BCW-201832 | *Enterobacter* | DSN | 1.13 | 1.2 | 53.26 | 781.19 |
| BCW-200015 | *Enterobacter* | DSN | 1.38 | 1 | 13.68 | 780.16 |
| BCW-200082 | *Enterobacter* | DSN | 1.02 | 1 | 0.02 | 761.76 |
| BCW-201812 | *Enterobacter* | DSN | 2.49 | 1.1 | 22.67 | 688.14 |
| BCW-200883 | *Enterobacter* | DSN | 2.5 | 1.2 | 13.74 | 669.73 |
| BCW-201847 | *Enterobacter* | DSN | 2.82 | 1.2 | 14.48 | 650.31 |
| BCW-201975 | *Enterobacter* | DSN | 2.3 | 1.1 | 16.36 | 556.24 |
| BCW-201899 | *Enterobacter* | DSN | 1.32 | 1.2 | 21.81 | 532.72 |
| BCW-201949 | *Enterobacter* | DSN | 0.87 | 1 | 11.65 | 503.07 |
| BCW-201884 | *Enterobacter* | DSN | 1.42 | 1.3 | 23.46 | 434.56 |
| BCW-201889 | *Enterobacter* | DSN | 4.74 | 1.5 | 18.78 | 298.57 |
| BCW-200319 | *Erwinia* | DSN | 2.37 | 1 | 7.02 | 1390.75 |
| BCW-201865 | *Erwinia* | DSN | 1.01 | 1.1 | 2.9 | 1349.69 |
| BCW-201853 | *Erwinia* | DSN | 1.09 | 1.1 | 30.36 | 858.9 |
| BCW-201854 | *Erwinia* | DSN | 0.87 | 1.1 | 15.04 | 400.82 |
| BCW-201173 | *Escherichia* | DSN | 1.3 | 1.5 | 24.33 | 980.57 |
| BCW-201236 | *Hafnia* | DSN | 1.42 | 1.4 | 6.54 | 1262.78 |
| BCW-201056 | *Hafnia* | DSN | 1.63 | 1.1 | 13.18 | 1097.14 |
| BCW-201450 | *Hafnia* | DSN | 2.66 | 1 | 18.78 | 514.31 |
| BCW-200121 | *Lactococcus* | DSN | 3.14 | 2.7 | 20.84 | 1257.22 |
| BCW-200163 | *Lactococcus* | DSN | 3.22 | 2.8 | 140.14 | 1098.43 |
| BCW-200138 | *Lactococcus* | DSN | 3.26 | 3.1 | 23.72 | 1041.99 |
| BCW-200188 | *Lactococcus* | DSN | 1.56 | 0.9 | 2.2 | 984.25 |
| BCW-201861 | *Lactococcus* | DSN | 1.54 | 1.2 | 1.17 | 976.48 |
| BCW-201453 | *Lactococcus* | DSN | 2.72 | 1 | 18.58 | 945.99 |
| BCW-200174 | *Lactococcus* | DSN | 1.48 | 3 | 28.95 | 847.77 |
| BCW-200158 | *Lactococcus* | DSN | 1.72 | No Data | -0.77 | 808.4 |
| BCW-200160 | *Lactococcus* | DSN | 2.51 | No Data | 24.67 | 774.28 |
| BCW-200241 | *Lactococcus* | DSN | 1.78 | 4.2 | 3.02 | 742.78 |
| BCW-200198 | *Lactococcus* | DSN | 1.32 | 3.4 | 1.46 | 724.41 |
| BCW-200238 | *Lactococcus* | DSN | 1.61 | 1.1 | 5.74 | 723.1 |
| BCW-200229 | *Lactococcus* | DSN | 2.07 | No Data | 0.26 | 612.86 |
| BCW-200180 | *Lactococcus* | DSN | 1.64 | 0.8 | -4.14 | 574.8 |
| BCW-200232 | *Lactococcus* | DSN | 1.31 | No Data | 10.67 | 557.74 |
| BCW-200128 | *Lactococcus* | DSN | 2.27 | 3 | 1.13 | 526.25 |
| BCW-200150 | *Lactococcus* | DSN | 1.68 | No Data | 15.61 | 498.69 |
| BCW-200159 | *Lactococcus* | DSN | 2.27 | 2.9 | 37.67 | 329.44 |

| Isolate ID | Genus | Group | ACC (RGR) | BNF (15N/14N) | IAA (mg/mL) | PO4 (mg/L) |
|---|---|---|---|---|---|---|
| BCW-200196 | *Lactococcus* | DSN | 1.34 | 2.8 | -0.11 | 310.5 |
| BCW-200077 | *Lactococcus* | DSN | 1.22 | 0.9 | -4.06 | 226.72 |
| BCW-200051 | *Lactococcus* | DSN | 1.26 | 2.2 | 2.32 | 222.56 |
| BCW-200192 | *Lactococcus* | DSN | 1.39 | 0.9 | 82.57 | 176.3 |
| BCW-200175 | *Lactococcus* | DSN | 1.68 | 3.7 | 26.44 | 149.78 |
| BCW-200634 | *Lelliottia* | DSN | 1.74 | 1.1 | 14.76 | 1446.83 |
| BCW-200596 | *Lelliottia* | DSN | 2.27 | 1.3 | 2.54 | 1142.13 |
| BCW-200275 | *Lelliottia* | DSN | 1.2 | 1.2 | 3.74 | 907.98 |
| BCW-201260 | *Lelliottia* | DSN | 1 | 1.2 | 1.73 | 899.8 |
| BCW-200271 | *Lelliottia* | DSN | 2.29 | 1 | 16.36 | 852.5 |
| BCW-201103 | *Lelliottia* | DSN | 2.46 | 1 | 17.2 | 844 |
| BCW-200033 | *Lelliottia* | DSN | 0.95 | 0.8 | 4.38 | 803.68 |
| BCW-201258 | *Lelliottia* | DSN | 1.63 | 1 | 1.98 | 800.57 |
| BCW-200270 | *Lelliottia* | DSN | 1.36 | 1.3 | 9.41 | 798.57 |
| BCW-200994 | *Lelliottia* | DSN | 2.2 | 1 | 19.41 | 789.24 |
| BCW-200641 | *Lelliottia* | DSN | 2.29 | 1 | 30.31 | 779.14 |
| BCW-200269 | *Lelliottia* | DSN | 1.08 | 1 | 3.03 | 743.91 |
| BCW-200991 | *Lelliottia* | DSN | 0.91 | 1.1 | 7.3 | 718.81 |
| BCW-201084 | *Lelliottia* | DSN | 1.11 | 1.2 | 3.16 | 708.59 |
| BCW-200556 | *Lelliottia* | DSN | 1.06 | 1.3 | 0.69 | 705.52 |
| BCW-200989 | *Lelliottia* | DSN | 2.64 | 1 | 10.97 | 669.73 |
| BCW-200473 | *Lelliottia* | DSN | 1.05 | 1.2 | 4.99 | 649.28 |
| BCW-200040 | *Lelliottia* | DSN | 1.5 | 1 | 16.64 | 642.13 |
| BCW-200003 | *Lelliottia* | DSN | 1.33 | 0.9 | 20.14 | 641.1 |
| BCW-201151 | *Lelliottia* | DSN | 1.25 | 1.1 | 1.86 | 640.08 |
| BCW-201045 | *Lelliottia* | DSN | 2.24 | 1.3 | 16.82 | 633.95 |
| BCW-200071 | *Lelliottia* | DSN | 0.98 | 1 | 22.53 | 606.34 |
| BCW-200990 | *Lelliottia* | DSN | No Data | 1 | No Data | No Data |
| BCW-200060 | *Metakosakonia* | DSN | 1.16 | 1 | 31.99 | 757.67 |
| BCW-200067 | *Microbacterium* | DSN | 2.45 | 0.9 | 35.49 | 640.08 |
| BCW-200057 | *Microbacterium* | DSN | 1.22 | 2.4 | -3.89 | 245.62 |
| BCW-200115 | *Morganella* | DSN | 1.63 | 1 | 210.76 | 795.28 |
| BCW-201826 | *Pantoea* | DSN | 2.52 | 1.1 | 20.97 | 1561.35 |
| BCW-201827 | *Pantoea* | DSN | 1.11 | 1.1 | 27.61 | 1384.46 |
| BCW-201867 | *Pantoea* | DSN | 1.33 | 1.4 | 11.86 | 1349.69 |
| BCW-201848 | *Pantoea* | DSN | 1.47 | 1.1 | 28.73 | 1260.74 |
| BCW-201997 | *Pantoea* | DSN | 0.78 | 1 | 9.87 | 1220.77 |
| BCW-201833 | *Pantoea* | DSN | 1.25 | 1.1 | 41.78 | 1067.48 |
| BCW-201995 | *Pantoea* | DSN | 2.32 | 1 | 15.39 | 1001.7 |
| BCW-201917 | *Pantoea* | DSN | 1.93 | 1.4 | 8.47 | 990.8 |
| BCW-200952 | *Pantoea* | DSN | 2.88 | 1.1 | 6.46 | 891.62 |
| BCW-201897 | *Pantoea* | DSN | 1.33 | 1.2 | 30.18 | 873.21 |
| BCW-201864 | *Pantoea* | DSN | 2.05 | 1.2 | 19.82 | 775.05 |
| BCW-201081 | *Pantoea* | DSN | 2.47 | 1 | 9.26 | 642.13 |
| BCW-201909 | *Pantoea* | DSN | 1.93 | 1.2 | 19.95 | 541.92 |
| BCW-202001 | *Pantoea* | DSN | No Data | 1 | No Data | No Data |
| BCW-200471 | *Pseudomonas* | DSN | 2.29 | 1.1 | 6.16 | 1166.67 |
| BCW-200718 | *Rahnella* | DSN | 1.26 | 1.1 | 2.11 | 2051.12 |
| BCW-200146 | *Rahnella* | DSN | 1.73 | 0.9 | 16.89 | 1767.72 |
| BCW-200157 | *Rahnella* | DSN | 1.47 | 0.8 | 110.55 | 1574.8 |
| BCW-200151 | *Rahnella* | DSN | 1.74 | 0.8 | 20.55 | 1493.44 |
| BCW-201008 | *Rahnella* | DSN | 1.46 | 1.1 | 0.43 | 1458.08 |
| BCW-201248 | *Rahnella* | DSN | 0.93 | 1 | 9.8 | 1309.54 |
| BCW-200806 | *Rahnella* | DSN | 2.49 | 1.1 | 6.16 | 1277.1 |

| Isolate ID | Genus | Group | ACC (RGR) | BNF (15N/14N) | IAA (mg/mL) | PO4 (mg/L) |
|---|---|---|---|---|---|---|
| BCW-200642 | *Rahnella* | DSN | 1.24 | 1.1 | 6.92 | 1275.05 |
| BCW-200808 | *Rahnella* | DSN | 2.09 | 1.3 | 1.83 | 1222.9 |
| BCW-201175 | *Rahnella* | DSN | 1.24 | 1.1 | 46.62 | 1178.94 |
| BCW-200724 | *Rahnella* | DSN | 1.14 | 1.1 | 7.43 | 1169.73 |
| BCW-200815 | *Rahnella* | DSN | 2.79 | 1.1 | 92.62 | 1139.06 |
| BCW-200155 | *Rahnella* | DSN | 2.07 | 0.9 | 29.65 | 1125.98 |
| BCW-200723 | *Rahnella* | DSN | 1.34 | 1.2 | 26.72 | 1125.77 |
| BCW-200715 | *Rahnella* | DSN | 1.24 | 1.2 | 8.55 | 1096.11 |
| BCW-201025 | *Rahnella* | DSN | 1.62 | 1.2 | 12.21 | 1095.09 |
| BCW-201014 | *Rahnella* | DSN | 1.35 | 1.2 | 5.47 | 1076.69 |
| BCW-201245 | *Rahnella* | DSN | 1.21 | 1.2 | 18.32 | 1049.08 |
| BCW-200725 | *Rahnella* | DSN | 1.35 | 1.2 | 34.22 | 1046.01 |
| BCW-201982 | *Rahnella* | DSN | 1.03 | 1 | 3.61 | 1032.72 |
| BCW-200810 | *Rahnella* | DSN | 1.22 | 1.3 | 16.49 | 1027.61 |
| BCW-201036 | *Rahnella* | DSN | 3.23 | 1 | 83.54 | 1025.56 |
| BCW-200149 | *Rahnella* | DSN | 1.42 | 0.8 | 11.7 | 1023.62 |
| BCW-201007 | *Rahnella* | DSN | 1.03 | 1 | 9.8 | 1022.47 |
| BCW-200736 | *Rahnella* | DSN | 1.02 | 1.1 | 64.22 | 1006.13 |
| BCW-201654 | *Rahnella* | DSN | 1.16 | 1 | 26.97 | 1003.59 |
| BCW-201176 | *Rahnella* | DSN | 1.23 | 1.2 | 5.34 | 1000 |
| BCW-200272 | *Rahnella* | DSN | 3.22 | 1 | 15.52 | 992.26 |
| BCW-201010 | *Rahnella* | DSN | 2.56 | 1 | 12.04 | 986.71 |
| BCW-201013 | *Rahnella* | DSN | 3.15 | 1 | 36.84 | 955.01 |
| BCW-201028 | *Rahnella* | DSN | 2.37 | 1 | 4.25 | 916.71 |
| BCW-201187 | *Rahnella* | DSN | 2.4 | 1.3 | 5.39 | 913.09 |
| BCW-200716 | *Rahnella* | DSN | 1.05 | 1 | 53.44 | 894.68 |
| BCW-200814 | *Rahnella* | DSN | 1.06 | 1 | 26.41 | 827.95 |
| BCW-200649 | *Rahnella* | DSN | 2.88 | 1 | 5.73 | 824.13 |
| BCW-201024 | *Rahnella* | DSN | 1.96 | 1 | 0.05 | 791.12 |
| BCW-200152 | *Rahnella* | DSN | 2.36 | 0.9 | 85.98 | 778.22 |
| BCW-200561 | *Rahnella* | DSN | 2.21 | 1.2 | 24.25 | 766.87 |
| BCW-200565 | *Rahnella* | DSN | 2.81 | 1.1 | 9.67 | 761.76 |
| BCW-201648 | *Rahnella* | DSN | 2.41 | 1 | 11.48 | 756.19 |
| BCW-200726 | *Rahnella* | DSN | 2.71 | 1.5 | 26.54 | 721.88 |
| BCW-200213 | *Rahnella* | DSN | 1.24 | 1 | 6.97 | 695.54 |
| BCW-201186 | *Rahnella* | DSN | 2.99 | 1 | 8.8 | 640.98 |
| BCW-200545 | *Rahnella* | DSN | 2.09 | 1 | 7.1 | 565.44 |
| BCW-200143 | *Rahnella* | DSN | No Data | No Data | No Data | No Data |
| BCW-200144 | *Rahnella* | DSN | No Data | 0.8 | No Data | No Data |
| BCW-200145 | *Rahnella* | DSN | No Data | 0.8 | No Data | No Data |
| BCW-200564 | *Rahnella* | DSN | No Data | 1 | No Data | No Data |
| BCW-200533 | *Rhodococcus* | DSN | 3.28 | 1.3 | 14.61 | 457.06 |
| BCW-200544 | *Serratia* | DSN | 2.39 | 1 | 3.05 | 1581.8 |
| BCW-201257 | *Serratia* | DSN | 2.94 | 1.1 | 18.8 | 1266.87 |
| BCW-201153 | *Serratia* | DSN | 2.05 | 1.3 | 6.72 | 1208.59 |
| BCW-201079 | *Serratia* | DSN | 1.92 | 1.1 | 44.61 | 1080.78 |
| BCW-200327 | *Serratia* | DSN | 2.26 | 1 | 14.43 | 1023.42 |
| BCW-200547 | *Serratia* | DSN | 1.32 | 1.1 | 6.01 | 1002.04 |
| BCW-200114 | *Serratia* | DSN | 2.18 | 1.2 | 22.32 | 981.63 |
| BCW-201085 | *Serratia* | DSN | 2.65 | 1.4 | 13.49 | 919.22 |
| BCW-201809 | *Serratia* | DSN | 2.11 | 1.5 | 23.87 | 906.95 |
| BCW-201350 | *Serratia* | DSN | 1.83 | 1 | 6.79 | 861.96 |
| BCW-201662 | *Serratia* | DSN | 2.29 | 1 | 18.09 | 849.67 |
| BCW-201054 | *Serratia* | DSN | 2.39 | 1 | 4.22 | 839.28 |

| Isolate ID | Genus | Group | ACC (RGR) | BNF (15N/14N) | IAA (mg/mL) | PO4 (mg/L) |
|---|---|---|---|---|---|---|
| BCW-201238 | *Serratia* | DSN | 3.04 | 1 | 13.94 | 832.67 |
| BCW-201185 | *Serratia* | DSN | 1.35 | 1.2 | 15.29 | 755.62 |
| BCW-200061 | *Serratia* | DSN | 1.56 | 1 | 23.43 | 713.7 |
| BCW-201653 | *Serratia* | DSN | 1.16 | 1 | 14.25 | 677.81 |
| BCW-201051 | *Serratia* | DSN | 2.65 | 1 | 15.9 | 616.43 |
| BCW-201835 | *Staphylococcus* | DSN | 1.65 | 1.2 | 2.54 | 63.83 |
| BCW-201883 | *unassigned* | DSN | 1.14 | 1.3 | 15.01 | 1154.4 |
| BCW-201090 | *unassigned* | DSN | 2.28 | 1.2 | 4.35 | 1002.04 |
| BCW-200903 | *unassigned* | DSN | 2.11 | 1.1 | 7.18 | 937.63 |
| BCW-200231 | *unassigned* | DSN | 0.85 | 0.9 | 14.58 | 923.88 |
| BCW-201845 | *unassigned* | DSN | 1.06 | 1.1 | 15.32 | 904.91 |
| BCW-200988 | *unassigned* | DSN | 2.27 | 1 | 14.68 | 896.73 |
| BCW-200315 | *unassigned* | DSN | 1.92 | 1 | 6.49 | 892.16 |
| BCW-200542 | *unassigned* | DSN | 1.58 | 1.1 | 13.51 | 757.13 |
| BCW-200094 | *unassigned* | DSN | 0.94 | 1.1 | 11.09 | 720.86 |
| BCW-200079 | *unassigned* | DSN | 1.18 | 0.6 | -3.77 | 364.01 |
| BCW-200063 | *unassigned* | DSN | 3.11 | 0.8 | 6.89 | 292.43 |
| BCW-201891 | *unassigned* | DSN | 4.32 | 1.1 | 1.76 | 248.64 |
| BCW-200912 | *unassigned* | DSN | 3.01 | 1.1 | 103.41 | 115.23 |
| BCW-200669 | *Enterobacter* | DSP | 2.19 | 1.1 | 6.97 | 1165.64 |
| BCW-200216 | *Enterobacter* | DSP | 2.27 | 1 | 9.56 | 946.19 |
| BCW-200317 | *Enterobacter* | DSP | 2.63 | 1 | 20.38 | 936.54 |
| BCW-201659 | *Enterobacter* | DSP | 1.04 | 1.2 | 3.38 | 920.52 |
| BCW-200109 | *Enterobacter* | DSP | 2.32 | 1 | 57.09 | 909.45 |
| BCW-200102 | *Enterobacter* | DSP | 2.58 | 1.1 | 17.38 | 902.89 |
| BCW-200050 | *Enterobacter* | DSP | 2.02 | 1 | 5.86 | 866.05 |
| BCW-200034 | *Enterobacter* | DSP | 2.25 | 1 | 7.51 | 839.47 |
| BCW-200026 | *Enterobacter* | DSP | 2.25 | 0.9 | 8.78 | 834.36 |
| BCW-200014 | *Enterobacter* | DSP | 2.15 | 1 | 17.84 | 821.06 |
| BCW-201721 | *Enterobacter* | DSP | 2.4 | 1 | 7.18 | 738.24 |
| BCW-200122 | *Enterobacter* | DSP | 2.3 | 0.9 | 23.56 | 721.78 |
| BCW-200095 | *Enterobacter* | DSP | 2.11 | 1 | 11.58 | 698.36 |
| BCW-200016 | *Enterobacter* | DSP | 2.38 | 1 | 6.23 | 685.07 |
| BCW-200206 | *Enterobacter* | DSP | 3.14 | 1 | 13.88 | 683.73 |
| BCW-200667 | *Enterobacter* | DSP | 1.57 | 1 | 3.72 | 641.1 |
| BCW-200017 | *Enterobacter* | DSP | 2.12 | 1 | 23.43 | 623.72 |
| BCW-200035 | *Enterobacter* | DSP | 2.38 | 1 | 9.69 | 615.54 |
| BCW-201441 | *Klebsiella* | DSP | 3.31 | 1 | 14.66 | 1310.84 |
| BCW-200129 | *Klebsiella* | DSP | 2.74 | 1 | 38.86 | 1199.48 |
| BCW-200172 | *Klebsiella* | DSP | 2.35 | 0.9 | 36.97 | 1087.93 |
| BCW-200167 | *Klebsiella* | DSP | 2.34 | 2.7 | 44.91 | 1053.81 |
| BCW-200177 | *Klebsiella* | DSP | 2.54 | 0.9 | 40.63 | 1035.43 |
| BCW-201019 | *Klebsiella* | DSP | 2.13 | 1.1 | 33.13 | 1017.38 |
| BCW-200136 | *Klebsiella* | DSP | 2.79 | 0.8 | 43.6 | 1011.81 |
| BCW-200123 | *Klebsiella* | DSP | 1.94 | 0.9 | 43.51 | 998.69 |
| BCW-200124 | *Klebsiella* | DSP | 2.34 | 0.8 | 47.38 | 985.56 |
| BCW-200086 | *Klebsiella* | DSP | 1.99 | 1 | 32.86 | 976.48 |
| BCW-200093 | *Klebsiella* | DSP | 2.05 | 1 | 36.97 | 907.98 |
| BCW-201021 | *Klebsiella* | DSP | 2.1 | 1 | 28.12 | 907.98 |
| BCW-200083 | *Klebsiella* | DSP | 2.01 | 1 | 38.16 | 897.75 |
| BCW-200137 | *Klebsiella* | DSP | 2.7 | 0.9 | 41 | 880.58 |
| BCW-200099 | *Klebsiella* | DSP | 2.28 | 1 | 42.9 | 873.21 |
| BCW-200651 | *Klebsiella* | DSP | 2.59 | 1 | 25.45 | 855.34 |
| BCW-200096 | *Klebsiella* | DSP | 2.08 | 1 | 45.41 | 848.67 |

| Isolate ID | Genus | Group | ACC (RGR) | BNF (15N/14N) | IAA (mg/mL) | PO4 (mg/L) |
|---|---|---|---|---|---|---|
| BCW-200084 | *Klebsiella* | DSP | 2.03 | 1 | 35.61 | 830.27 |
| BCW-200069 | *Klebsiella* | DSP | 2.02 | 1 | 42.03 | 818 |
| BCW-200132 | *Klebsiella* | DSP | 2.24 | 0.9 | 38.86 | 807.09 |
| BCW-201263 | *Klebsiella* | DSP | 1.92 | 1.1 | 35.47 | 797.55 |
| BCW-200049 | *Klebsiella* | DSP | 2.47 | 1 | 40.84 | 754.6 |
| BCW-200053 | *Klebsiella* | DSP | 2.11 | 1 | 37.55 | 706.54 |
| BCW-200662 | *Klebsiella* | DSP | 2.76 | 1.2 | 32.16 | 701.43 |
| BCW-200660 | *Klebsiella* | DSP | 2.21 | 1 | 19.44 | 671.78 |
| BCW-201020 | *Klebsiella* | DSP | 2.02 | 1 | 23.26 | 558.28 |
| BCW-200183 | *Kosakonia* | DSP | 2.31 | 0.9 | 3.84 | 1325.46 |
| BCW-200181 | *Kosakonia* | DSP | 1.67 | 4.6 | 10.59 | 992.13 |
| BCW-200226 | *Kosakonia* | DSP | 1.97 | 1 | 5.98 | 784.78 |
| BCW-200227 | *Kosakonia* | DSP | 2.12 | 1 | 5.78 | 732.28 |
| BCW-200214 | *Kosakonia* | DSP | 2.09 | 0.9 | 3.88 | 729.66 |
| BCW-200141 | *Kosakonia* | DSP | 2.41 | 1 | 3.14 | 709.97 |
| BCW-200194 | *Kosakonia* | DSP | 2.48 | 0.9 | 6.77 | 703.41 |
| BCW-200210 | *Kosakonia* | DSP | 2.08 | No Data | 12.44 | 702.1 |
| BCW-200197 | *Kosakonia* | DSP | 2.11 | 0.9 | 8.08 | 674.54 |
| BCW-200162 | *Metakosakonia* | DSP | 3.7 | 0.8 | 158 | 1171.92 |
| BCW-200308 | *Metakosakonia* | DSP | 1.97 | 1 | 14.25 | 1099.91 |
| BCW-200168 | *Metakosakonia* | DSP | 2.04 | 0.8 | 38.45 | 1053.81 |
| BCW-200509 | *Metakosakonia* | DSP | 2.3 | 1 | 23.05 | 1002.64 |
| BCW-200307 | *Metakosakonia* | DSP | 1.65 | 1.1 | 36.59 | 916.16 |
| BCW-201858 | *Metakosakonia* | DSP | 1.65 | 1.3 | 23.16 | 911.04 |
| BCW-201850 | *Metakosakonia* | DSP | 2.34 | 1.2 | 26.46 | 876.28 |
| BCW-200517 | *Metakosakonia* | DSP | 2.53 | 1.2 | 32.77 | 867.08 |
| BCW-200951 | *Metakosakonia* | DSP | 3.01 | 1.1 | 103.54 | 839.47 |
| BCW-201886 | *Metakosakonia* | DSP | 1.77 | 1.2 | 23.77 | 834.36 |
| BCW-200567 | *Metakosakonia* | DSP | 1.35 | 1.1 | 16.54 | 762.78 |
| BCW-200955 | *Metakosakonia* | DSP | 2.47 | 1.2 | 22.75 | 722.9 |
| BCW-201155 | *Metakosakonia* | DSP | 1.15 | 1 | 21.86 | 670.25 |
| BCW-201058 | *Metakosakonia* | DSP | 2.12 | 1 | 25.78 | 639.09 |
| BCW-201972 | *Metakosakonia* | DSP | 2.16 | 1.1 | 19.47 | 600.2 |
| BCW-201828 | *Metakosakonia* | DSP | 2.47 | 1 | 22.26 | 582.82 |
| BCW-200656 | *Metakosakonia* | DSP | 2.75 | 1.1 | 22.16 | 550.1 |
| BCW-201937 | *Metakosakonia* | DSP | 1.83 | 1 | 21.4 | 528.63 |
| BCW-201876 | *Metakosakonia* | DSP | 1.47 | 1.3 | 18.96 | 519.43 |
| BCW-201873 | *Metakosakonia* | DSP | 1.7 | 1.3 | 22.93 | 512.27 |
| BCW-201990 | *Metakosakonia* | DSP | 1.69 | 1.2 | 22.21 | 507.16 |
| BCW-201887 | *Metakosakonia* | DSP | 1.31 | 1.3 | 21.7 | 480.57 |
| BCW-201888 | *Metakosakonia* | DSP | 1.69 | 1.3 | 17.33 | 480.57 |
| BCW-201879 | *Metakosakonia* | DSP | 1.45 | 1.4 | 21.32 | 448.88 |
| BCW-201808 | *Metakosakonia* | DSP | 2.74 | 1.2 | 33.56 | 444.79 |
| BCW-200785 | *Pseudomonas* | DSP | 3.31 | 1 | 15.22 | 823.23 |
| BCW-201290 | *Pseudomonas* | DSP | 1.05 | 1.1 | 12.7 | 391.62 |
| BCW-201304 | *Rahnella* | DSP | 2.11 | 1.1 | 0.18 | 1798.57 |
| BCW-201315 | *Rahnella* | DSP | 2.11 | 1.1 | 3.38 | 1111.45 |
| BCW-200647 | *Rahnella* | DSP | 1.83 | 1 | 0.28 | 1086.91 |
| BCW-200578 | *Rahnella* | DSP | 0.8 | 1.1 | 1.02 | 1038.85 |
| BCW-201297 | *Rahnella* | DSP | 1.54 | 1 | 6.36 | 1032.86 |
| BCW-200828 | *Rahnella* | DSP | 2.38 | 1.1 | 4.86 | 1022.49 |
| BCW-200650 | *Rahnella* | DSP | 2.04 | 1.1 | 48.22 | 902.86 |
| BCW-200818 | *Rahnella* | DSP | 2.54 | 1 | 22.62 | 887.53 |
| BCW-200801 | *Rahnella* | DSP | 2.46 | 1.1 | 99.62 | 871.17 |

| Isolate ID | Genus | Group | ACC (RGR) | BNF (15N/14N) | IAA (mg/mL) | PO4 (mg/L) |
|---|---|---|---|---|---|---|
| BCW-200552 | *Rahnella* | DSP | 1.57 | 1.1 | 34.66 | 870.14 |
| BCW-201302 | *Rahnella* | DSP | 2.04 | 1.1 | 2.67 | 866.05 |
| BCW-200559 | *Rahnella* | DSP | 2.36 | 1 | 0.41 | 853.45 |
| BCW-201091 | *Rahnella* | DSP | 1.81 | 1 | 21.53 | 830.78 |
| BCW-200800 | *Rahnella* | DSP | 3.26 | 1.1 | 8.14 | 816.97 |
| BCW-201184 | *Rahnella* | DSP | 1.45 | 1.1 | 2.54 | 811.86 |
| BCW-201162 | *Rahnella* | DSP | 2.79 | 1.2 | 10.56 | 796.52 |
| BCW-201070 | *Rahnella* | DSP | 1.97 | 1 | 12.62 | 759.96 |
| BCW-200797 | *Rahnella* | DSP | 2.23 | 1.1 | 26.9 | 755.62 |
| BCW-201059 | *Rahnella* | DSP | 2.33 | 1 | 39.69 | 720.86 |
| BCW-200821 | *Rahnella* | DSP | 2.45 | 1.1 | 24.38 | 687.12 |
| BCW-201088 | *Rahnella* | DSP | 3.28 | 1 | 47.23 | 672.8 |
| BCW-200798 | *Rahnella* | DSP | 2.25 | 1 | 8.09 | 548.06 |
| BCW-200644 | *Rahnella* | DSP | No Data | 1 | No Data | No Data |
| BCW-201259 | *Rahnella* | DSP | No Data | 1.2 | No Data | No Data |
| BCW-201649 | *Raoultella* | DSP | 1.6 | 1 | 16.51 | 2214.72 |
| BCW-200600 | *Raoultella* | DSP | 1.58 | 1.2 | 3.89 | 1246.42 |
| BCW-201703 | *Raoultella* | DSP | 1.91 | 1.1 | 13.33 | 1195.3 |
| BCW-200555 | *Raoultella* | DSP | 2.76 | 1.1 | 18.88 | 1148.26 |
| BCW-200449 | *Raoultella* | DSP | 1.17 | 1.1 | 10.46 | 1102.25 |
| BCW-200117 | *Raoultella* | DSP | 2.57 | 1.1 | 1.87 | 1099.74 |
| BCW-200444 | *Raoultella* | DSP | 2.5 | 1.5 | 37.68 | 1095.09 |
| BCW-200169 | *Raoultella* | DSP | 2.39 | 2.7 | 30.92 | 1078.74 |
| BCW-200142 | *Raoultella* | DSP | 2.43 | 0.9 | 5.08 | 1076.12 |
| BCW-201900 | *Raoultella* | DSP | 1.44 | 1.2 | 43.03 | 1071.57 |
| BCW-200120 | *Raoultella* | DSP | 2.45 | 0.9 | 20.14 | 1052.49 |
| BCW-200171 | *Raoultella* | DSP | 2.45 | 0.8 | 23.1 | 1036.75 |
| BCW-200704 | *Raoultella* | DSP | 1.33 | 1 | 2.09 | 1024.54 |
| BCW-200892 | *Raoultella* | DSP | 2.68 | 1 | 4.48 | 1018.7 |
| BCW-200577 | *Raoultella* | DSP | 2.22 | 1 | 17.74 | 995.09 |
| BCW-200891 | *Raoultella* | DSP | 2.89 | 1 | 12.06 | 992.26 |
| BCW-200203 | *Raoultella* | DSP | 2.87 | 1 | 6.6 | 981.63 |
| BCW-200885 | *Raoultella* | DSP | 1.51 | 1.2 | 25.9 | 981.6 |
| BCW-200030 | *Raoultella* | DSP | 1.49 | 0.6 | 6.02 | 970.35 |
| BCW-200281 | *Raoultella* | DSP | 2.68 | 1.1 | 22.37 | 956.03 |
| BCW-200648 | *Raoultella* | DSP | 2.11 | 1.1 | 10.38 | 952.97 |
| BCW-200437 | *Raoultella* | DSP | 2.84 | 1 | 3.1 | 952.6 |
| BCW-200442 | *Raoultella* | DSP | 2.53 | 1 | 13.41 | 944.1 |
| BCW-200440 | *Raoultella* | DSP | 1.78 | 1 | 16.36 | 910.1 |
| BCW-200874 | *Raoultella* | DSP | 2.05 | 1 | 2.75 | 897.83 |
| BCW-201098 | *Raoultella* | DSP | 1.59 | 1.2 | 17.35 | 897.75 |
| BCW-200438 | *Raoultella* | DSP | 2.58 | 1 | 3.54 | 893.66 |
| BCW-201443 | *Raoultella* | DSP | 2.28 | 1 | 19.19 | 892.16 |
| BCW-200738 | *Raoultella* | DSP | 2.78 | 1 | 17.51 | 873.28 |
| BCW-201461 | *Raoultella* | DSP | 2.41 | 1 | 20.97 | 866.67 |
| BCW-201901 | *Raoultella* | DSP | 0.93 | 1.2 | 29.52 | 852.76 |
| BCW-200820 | *Raoultella* | DSP | 2.33 | 1 | 27.66 | 851.74 |
| BCW-200106 | *Raoultella* | DSP | 2.46 | 0.8 | 27.22 | 838.58 |
| BCW-200525 | *Raoultella* | DSP | 2.47 | 1.1 | 60.61 | 824.13 |
| BCW-200776 | *Raoultella* | DSP | 2.95 | 1.1 | 5.73 | 814.93 |
| BCW-200886 | *Raoultella* | DSP | 2.12 | 1.1 | 6.79 | 813.91 |
| BCW-200161 | *Raoultella* | DSP | 2.97 | 0.8 | 15.78 | 812.34 |
| BCW-200521 | *Raoultella* | DSP | 1.34 | 1 | 20.03 | 809.82 |
| BCW-201071 | *Raoultella* | DSP | 2.38 | 1 | 17.23 | 804.7 |

| Isolate ID | Genus | Group | ACC (RGR) | BNF (15N/14N) | IAA (mg/mL) | PO4 (mg/L) |
|---|---|---|---|---|---|---|
| BCW-200727 | *Raoultella* | DSP | 1.96 | 1.4 | 4.07 | 793.46 |
| BCW-200880 | *Raoultella* | DSP | 1.87 | 1.1 | 0.46 | 781.19 |
| BCW-200926 | *Raoultella* | DSP | 2.54 | 1.4 | 22.57 | 781.19 |
| BCW-201075 | *Raoultella* | DSP | 2.26 | 1 | 6.28 | 780.74 |
| BCW-200915 | *Raoultella* | DSP | 2.54 | 1 | 0.05 | 780.16 |
| BCW-201890 | *Raoultella* | DSP | 2.61 | 1 | 13.23 | 776.07 |
| BCW-200294 | *Raoultella* | DSP | 2.38 | 1 | 1.73 | 768.46 |
| BCW-200446 | *Raoultella* | DSP | 2.72 | 1.1 | 10.81 | 767.89 |
| BCW-200855 | *Raoultella* | DSP | 2.47 | 1.1 | 4.45 | 757.67 |
| BCW-200553 | *Raoultella* | DSP | 2.37 | 1.1 | 17 | 738.24 |
| BCW-201044 | *Raoultella* | DSP | 2.15 | 1 | 3 | 736.36 |
| BCW-201107 | *Raoultella* | DSP | 2.58 | 1 | 17.48 | 736.36 |
| BCW-200909 | *Raoultella* | DSP | 2.41 | 1 | 3.56 | 736.2 |
| BCW-200879 | *Raoultella* | DSP | 1.76 | 1 | 9.19 | 734.15 |
| BCW-200496 | *Raoultella* | DSP | 1.7 | 1.2 | 3.1 | 725.97 |
| BCW-200881 | *Raoultella* | DSP | 2.56 | 1.1 | 0.69 | 719.84 |
| BCW-201614 | *Raoultella* | DSP | 2.89 | 1 | 23.56 | 713.7 |
| BCW-200499 | *Raoultella* | DSP | 2.52 | 1 | 23.74 | 709.61 |
| BCW-200665 | *Raoultella* | DSP | 3.08 | 1.2 | 7.61 | 695.3 |
| BCW-201615 | *Raoultella* | DSP | 2.56 | 1 | 1.73 | 687.12 |
| BCW-201078 | *Raoultella* | DSP | 2.67 | 1 | 3.97 | 677.91 |
| BCW-200882 | *Raoultella* | DSP | 2.79 | 1.2 | 3.38 | 675.87 |
| BCW-200847 | *Raoultella* | DSP | 2.46 | 1.1 | 3.61 | 665.64 |
| BCW-200887 | *Raoultella* | DSP | 2.71 | 1 | 0.56 | 652.35 |
| BCW-200620 | *Raoultella* | DSP | 3 | 1.2 | 25.83 | 650.31 |
| BCW-200488 | *Raoultella* | DSP | 3.12 | 1.3 | 1.5 | 644.17 |
| BCW-200276 | *Raoultella* | DSP | 2.14 | 1 | 3.23 | 607.93 |
| BCW-201620 | *Raoultella* | DSP | 3.23 | 1 | 11.07 | 587.16 |
| BCW-200165 | *Raoultella* | DSP | 2.37 | 0.9 | 26.68 | 559.06 |
| BCW-201926 | *Raoultella* | DSP | 3.16 | 1.2 | 31.55 | 541.92 |
| BCW-200661 | *Raoultella* | DSP | 3.26 | 1.3 | 45.73 | 540.9 |
| BCW-200182 | *unassigned* | DSP | 2.24 | 0.8 | 34.26 | 1463.25 |
| BCW-200113 | *unassigned* | DSP | 2.26 | 1 | 61.74 | 1148.29 |
| BCW-200234 | *unassigned* | DSP | 2.31 | 0.9 | 24.26 | 1099.74 |
| BCW-200133 | *unassigned* | DSP | 2.09 | 0.8 | 17.09 | 1041.99 |
| BCW-200195 | *unassigned* | DSP | 2.61 | 0.9 | 4.5 | 988.19 |
| BCW-200148 | *unassigned* | DSP | 2.69 | 0.8 | 53.02 | 979 |
| BCW-200200 | *unassigned* | DSP | 2.54 | 0.8 | 8.04 | 976.38 |
| BCW-200156 | *unassigned* | DSP | 2.22 | 1 | 10.55 | 952.76 |
| BCW-200225 | *unassigned* | DSP | 1.85 | 0.9 | 24.5 | 952.76 |
| BCW-200199 | *unassigned* | DSP | 2.89 | 3.5 | 8.74 | 952.76 |
| BCW-200201 | *unassigned* | DSP | 2.59 | 1 | 4.54 | 917.32 |
| BCW-200236 | *unassigned* | DSP | 2.4 | 0.9 | 4.5 | 913.39 |
| BCW-200237 | *unassigned* | DSP | 2.35 | 1 | 5.37 | 898.95 |
| BCW-200221 | *unassigned* | DSP | 1.81 | 0.8 | 10.02 | 877.95 |
| BCW-201267 | *unassigned* | DSP | 2.88 | 1 | 21.07 | 865.03 |
| BCW-200184 | *unassigned* | DSP | 2.64 | 1 | 11.37 | 863.52 |
| BCW-200028 | *unassigned* | DSP | 0.84 | 1 | 7.05 | 815.95 |
| BCW-200235 | *unassigned* | DSP | 2.9 | No Data | 27.3 | 808.4 |
| BCW-201097 | *unassigned* | DSP | 2.26 | 1.1 | 30.92 | 593.05 |
| BCW-200011 | *Acidovorax* | SDS | 1.15 | 0.8 | 3.6 | 665.64 |
| BCW-200001 | *Acidovorax* | SDS | 2.64 | 0.9 | 0.1 | 414.11 |
| BCW-200112 | *Acinetobacter* | SDS | 3.14 | 0.8 | 33.19 | 1320.21 |
| BCW-200100 | *Acinetobacter* | SDS | 2.28 | 0.8 | 7.79 | 1270.34 |

| Isolate ID | Genus | Group | ACC (RGR) | BNF (15N/14N) | IAA (mg/mL) | PO4 (mg/L) |
|---|---|---|---|---|---|---|
| BCW-200119 | *Acinetobacter* | SDS | 2.33 | 0.8 | 10.26 | 988.19 |
| BCW-200032 | *Acinetobacter* | SDS | 1.26 | 0.8 | 13.64 | 757.67 |
| BCW-200105 | *Acinetobacter* | SDS | 2.86 | 0.9 | 8.86 | 738.85 |
| BCW-200116 | *Acinetobacter* | SDS | 1.45 | No Data | 1.66 | 618.11 |
| BCW-200118 | *Acinetobacter* | SDS | 3.08 | 0.9 | -0.64 | 603.67 |
| BCW-200008 | *Acinetobacter* | SDS | 2.88 | 1 | 28.82 | 553.17 |
| BCW-200081 | *Acinetobacter* | SDS | 2.19 | 0.9 | 11.17 | 472.39 |
| BCW-200044 | *Acinetobacter* | SDS | 2.02 | 1 | 4.71 | 469.33 |
| BCW-200101 | *Bacillus* | SDS | 1.51 | 0.9 | 9.11 | 702.1 |
| BCW-201819 | *Curtobacterium* | SDS | 2.11 | 1.2 | 1.04 | 949.9 |
| BCW-200993 | *Herbaspirillum* | SDS | 1.33 | 1 | 1.88 | 890.27 |
| BCW-200209 | *Leifsonia* | SDS | 1.15 | 1 | 2.57 | 872.7 |
| BCW-200078 | *Micrococcus* | SDS | 0.99 | 0.8 | 4.13 | 800.61 |
| BCW-200173 | *Micrococcus* | SDS | 1.47 | 2.6 | 0.39 | 443.57 |
| BCW-200436 | *Pseudomonas* | SDS | 1.9 | 1 | 15.06 | 1297.55 |
| BCW-200621 | *Pseudomonas* | SDS | 3.32 | 1.3 | 10.23 | 1161.55 |
| BCW-201292 | *Pseudomonas* | SDS | 1.14 | 1 | 11.3 | 1136.73 |
| BCW-200527 | *Pseudomonas* | SDS | 2.05 | 1 | 9.06 | 1086.69 |
| BCW-200599 | *Pseudomonas* | SDS | 1.4 | 1.1 | 17.94 | 1065.44 |
| BCW-201868 | *Pseudomonas* | SDS | 1.51 | 2.3 | 5.78 | 993.87 |
| BCW-200528 | *Pseudomonas* | SDS | 2.06 | 1 | 8.4 | 942.21 |
| BCW-200460 | *Pseudomonas* | SDS | 1.38 | 1.1 | 7.46 | 936.61 |
| BCW-200290 | *Pseudomonas* | SDS | 1.45 | 1.2 | 19.97 | 831.29 |
| BCW-201862 | *Pseudomonas* | SDS | 1.34 | 1.2 | 6.62 | 791.41 |
| BCW-200477 | *Pseudomonas* | SDS | 1.78 | 1.1 | 29.9 | 755.62 |
| BCW-200065 | *Pseudomonas* | SDS | 1.25 | 0.9 | 27.88 | 723.93 |
| BCW-201293 | *Pseudomonas* | SDS | 1.19 | 1 | 21.48 | 704.25 |
| BCW-200267 | *Pseudomonas* | SDS | 1.05 | 1 | 36.72 | 699.53 |
| BCW-200588 | *Pseudomonas* | SDS | 1.5 | 1.1 | 0.1 | 684.05 |
| BCW-201875 | *Pseudomonas* | SDS | 2.51 | 1.2 | 8.04 | 653.37 |
| BCW-200497 | *Pseudomonas* | SDS | 1.5 | 1 | 2.6 | 591.88 |
| BCW-200443 | *Pseudomonas* | SDS | 2.67 | 1.4 | 26.13 | 445.81 |
| BCW-200791 | *Pseudomonas* | SDS | 1.63 | 1 | 14.96 | 409.63 |
| BCW-201947 | *Pseudomonas* | SDS | 1.13 | 1 | 34.73 | 386.97 |
| BCW-200046 | *Pseudomonas* | SDS | 1.4 | 0.9 | 20.67 | 354.81 |
| BCW-200068 | *Pseudomonas* | SDS | 3.17 | 3 | 23.39 | 310.84 |
| BCW-200432 | *Pseudomonas* | SDS | 1.15 | 1 | 10.81 | 307.65 |
| BCW-200056 | *Pseudomonas* | SDS | 1.6 | 1.3 | 8.08 | 260.74 |
| BCW-200458 | *Pseudomonas* | SDS | 0.97 | 1.1 | 7.07 | 260.36 |
| BCW-201859 | *Pseudomonas* | SDS | 1.87 | 1.3 | 10.33 | 250.53 |
| BCW-200073 | *Pseudomonas* | SDS | 2.34 | 0.9 | 4.13 | 230.88 |
| BCW-200607 | *Pseudomonas* | SDS | 1.38 | 1.2 | 1.37 | 168.9 |
| BCW-200790 | *Pseudomonas* | SDS | 1.74 | 1 | 17.35 | 20.59 |
| BCW-200476 | *Pseudomonas* | SDS | No Data | 1 | No Data | No Data |
| BCW-200574 | *Stenotrophomonas* | SDS | 1.34 | 1.1 | 17.33 | 1414.11 |
| BCW-200587 | *Stenotrophomonas* | SDS | 2.2 | 1 | 14.15 | 1049.86 |
| BCW-200945 | *Stenotrophomonas* | SDS | 1.41 | 1 | 12.09 | 1017.75 |
| BCW-200154 | *Stenotrophomonas* | SDS | 2.86 | 1 | 50.76 | 883.2 |
| BCW-200938 | *Stenotrophomonas* | SDS | 1.38 | 1.2 | 16.56 | 820.04 |
| BCW-200266 | *Stenotrophomonas* | SDS | 1.89 | 1.1 | 1.35 | 816.97 |
| BCW-200202 | *Stenotrophomonas* | SDS | 2.19 | 1 | 4.01 | 775.59 |
| BCW-200103 | *Stenotrophomonas* | SDS | 1.68 | 2.1 | 7.01 | 770.34 |
| BCW-200570 | *Stenotrophomonas* | SDS | 2.45 | 1.2 | 20.48 | 724.95 |
| BCW-201851 | *Stenotrophomonas* | SDS | 1.89 | 1.3 | 2.54 | 716.77 |

| Isolate ID | Genus | Group | ACC (RGR) | BNF (15N/14N) | IAA (mg/mL) | PO4 (mg/L) |
|---|---|---|---|---|---|---|
| BCW-200153 | *Stenotrophomonas* | SDS | 2.11 | 0.8 | 0.96 | 686.35 |
| BCW-200589 | *Stenotrophomonas* | SDS | 1.49 | 1 | -0.08 | 677.81 |
| BCW-200944 | *Stenotrophomonas* | SDS | 0.91 | 1 | 1.3 | 329.37 |
| BCW-200646 | *Stenotrophomonas* | SDS | 1.59 | 1 | 1.76 | 66.86 |
| BCW-200939 | *Stenotrophomonas* | SDS | 1.13 | 1.2 | 0 | 47.58 |
| BCW-200147 | *unassigned* | SDS | 3.05 | 1 | 92.2 | 682.41 |
| BCW-200002 | *unassigned* | SDS | 1.94 | 1 | 5.12 | 451.94 |
| BCW-200498 | *unassigned* | SDS | 1.67 | 1.2 | 1.98 | 146.98 |

Performance on each *in vitro* PGP assay was summarized for each isolate. Each record indicates the identification code assigned to the mucilage isolate (BCW-ID). Genus assignment based on classification of the isolate's whole genome sequence generated with Sourmash 3.0.1 [2] and $^{15}N/^{14}N$ ratio values indicating performance on the $^{15}N$ incorporation assay (BNF) were included using data from previous investigations with the isolate collection [6]. *In vitro* assays for non-*nif* PGP traits included the utilization of 1-Amino-1-cyclopropanecarboxylic acid as a nitrogen source for growth (ACC), the colorimetric assay for auxin biosynthesis (IAA), and the colorimetric assay to detect the liberation of soluble phosphate (PO4). Units for measurements of the ACC, IAA and PO4 assays were relative growth rate (RGR – see Methods), mg/mL and mg/L, respectively.

**S3 Table. Summary statistics for PGP assays by NIF group**

| NIF Group | Assay | Units | Mean | Median | Min | Max |
|---|---|---|---|---|---|---|
| DSN | ACC | RGR | 1.74 | 1.5 | 0.76 | 4.74 |
| DSP | ACC | RGR | 2.26 | 2.31 | 0.8 | 3.7 |
| SDS | ACC | RGR | 1.82 | 1.63 | 0.91 | 3.32 |
| DSN | BNF | 15N/14N | 1.2 | 1.1 | 0.6 | 4.2 |
| DSP | BNF | 15N/14N | 1.09 | 1 | 0.6 | 4.6 |
| SDS | BNF | 15N/14N | 1.12 | 1 | 0.8 | 3 |
| DSN | IAA | mg/mL | 18.01 | 13.51 | -4.14 | 210.76 |
| DSP | IAA | mg/mL | 20.29 | 17.35 | 0.05 | 158 |
| SDS | IAA | mg/mL | 12.45 | 8.4 | -0.64 | 92.2 |
| DSN | PO4 | mg/L | 864.9 | 852.76 | 63.83 | 2051.12 |
| DSP | PO4 | mg/L | 853.68 | 839.47 | 391.62 | 2214.72 |
| SDS | PO4 | mg/L | 678.05 | 702.1 | 20.59 | 1414.11 |

Data from the *in vitro* assays for biological nitrogen fixation (BNF), ACC utilization (ACC), indole-3-acetic acid biosynthesis (IAA) and phosphate solubilization (PO4) were summarized in R using base and tidyverse 1.2.1 packages [3]. Summary statistics included estimations generated with data from all isolates for mean, median, minimum (min) and maximum (max) values. NIF group corresponds to Dos Santos Positive (DSP), Dos Santos Negative (DSN), and Semi-Dos Santos (SDS).
